# Frequencies of house fly proto-Y chromosomes across populations are predicted by temperature heterogeneity within populations

**DOI:** 10.1101/2024.05.15.594357

**Authors:** Patrick D. Foy, Sara R. Loetzerich, David Boxler, Edwin R. Burgess, R. T. Trout Fryxell, Alec C. Gerry, Nancy C. Hinkle, Erika T. Machtinger, Cassandra Olds, Aaron M. Tarone, Wes Watson, Jeffrey G. Scott, Richard P. Meisel

## Abstract

Sex chromosomes often differ between closely related species and can even be polymorphic within populations. Species with polygenic sex determination segregate for multiple different sex determining loci within populations, making them uniquely informative of the selection pressures that drive the evolution of sex chromosomes. The house fly (*Musca domestica*) is a model species for studying polygenic sex determination because male determining genes have been identified on all six of the chromosomes, which means that any chromosome can be a “proto-Y” chromosome. In addition, chromosome IV can carry a female-determining locus, making it a W chromosome. The different proto-Y chromosomes are distributed along latitudinal clines on multiple continents, their distributions can be explained by seasonality in temperature, and they have temperature-dependent effects on physiological and behavioral traits. It is not clear, however, how the clinal distributions interact with the effect of seasonality on the frequencies of house fly proto-Y chromosomes across populations. To address this question, we measured the frequencies of house fly Y and W chromosomes across nine populations in the United States of America. We confirmed the clinal distribution along the eastern coast of North America, but it is limited to the eastern coast. In contrast, annual mean daily temperature range is significantly correlated with proto-Y chromosome frequencies across the entire continent. Our results therefore suggest that temperature heterogeneity can explain the distributions of house fly proto-Y chromosomes in a way that does not depend on the cline. These results contribute to our understanding of how ecological factors affect sex chromosome evolution.

## Introduction

Sex chromosomes and sex determining genes often differ between even closely related species (Bachtrog et al. 2014; Beukeboom and Perrin 2014) . Evolutionary changes in sex chromosomes typically occur via one of two methods (Abbott et al. 2017) . First, an autosome can fuse to a sex chromosome, turning the autosome into a neo-sex chromosome. Second, an autosome can acquire a new master sex determining locus, turning the autosome into a proto-sex chromosome (and allowing the ancestral sex chromosome to “revert” to an autosome). Sex-specific selection pressures are thought to be important for the invasion and fixation of both neo- and proto-sex chromosomes because sex-linkage can resolve inter-sexual conflicts (van Doorn and Kirkpatrick 2007; Roberts et al. 2009; van Doorn 2014; Mank et al. 2014) . In addition, if sex ratios are distorted from their evolutionary stable equilibrium, a new sex determiner on a proto-sex chromosome can be favored if it increases the frequency of the sex that is below its equilibrium value (Bull 1983; Werren and Beukeboom 1998) . Notably, ecological factors can modulate the effects of these sex-specific selection pressures, but the extent to which ecological selection pressures drive sex chromosome evolution are not yet resolved (Meisel 2022) .

We used the house fly (*Musca domestica*) as a model species to explore how ecological factors affect sex chromosome evolution. House fly is well-suited for this purpose because it has a highly polymorphic multifactorial sex determination system (Hamm et al. 2015) . Male determining factors have been genetically mapped to all six of the house fly chromosome pairs, and a single gene (*Mdmd*) has been implicated as the male-determiner on at least four of the six chromosomes (Sharma et al. 2017) . Each of these *Mdmd*-bearing chromosomes is a young “proto-Y” chromosome (Meisel et al. 2017; Son and Meisel 2021) . Nearly every male house fly in North America carries one or both the two most abundant proto-Y chromosomes, the Y chromosome (Y ^M^) and third chromosome (III ^M^), and males with other proto-Y chromosomes are rarely found (Hamm et al. 2015) . Y ^M^ and III ^M^ form latitudinal clines in Europe, Japan, and North America, with Y ^M^ most common in northern populations and III ^M^ predominating in the south (Franco et al. 1982; Denholm et al. 1986; Tomita and Wada 1989; Hamm et al. 2005) . Consistent with this geographical distribution, the Y ^M^ chromosome confers greater tolerance to extreme cold and preference for cooler temperatures, while III ^M^ confers improved tolerance to extreme heat and preference for warmer temperatures (Delclos et al. 2021) . In addition, higher Y ^M^ frequency is associated with locations with low seasonality of temperatures, or small differences between minimum and maximum values of monthly high and low temperatures (Feldmeyer et al. 2008) . Moreover, in some house fly populations, males can carry multiple proto-Y chromosomes (e.g., both Y ^M^ and III ^M^, or homozygous for III ^M^), which could create male-biased sex ratios and selection in favor of a female-determining factor (Eshel 1975; Bull and Charnov 1977; Bulmer and Bull 1982) . Indeed, such a female-determiner exists in house fly populations, in the form of a dominant allele of the house fly ortholog of *transformer* (*Md-tra*^*D*^), which causes embryos to develop into females even if they carry multiple male-determining chromosomes (Hediger et al. 2010) . The frequency of *Md-tra*^*D*^ is correlated with the frequency of males with multiple male-determining chromosomes across populations, suggesting that there is selection for balanced sex-ratios (Meisel et al. 2016) .

We aimed to test if the frequencies of Y ^M^, III ^M^, and *Md-tra*^*D*^ across North American populations could be explained by climatic variables. Previous studies linking the frequencies of male-determining chromosomes to climatic variation in North America have been limited to a latitudinal cline along the eastern coast (Hamm et al. 2005; Hamm and Scott 2008), and only Japanese and African populations were sampled to study how *Md-tra*^*D*^ frequencies vary across climates (Feldmeyer et al. 2008) . However, the frequencies of Y ^M^, III ^M^ and *Md-tra*^*D*^ vary in non-clinal patterns across regions of North America outside the eastern coast (McDonald et al. 1975; Meisel et al. 2016), suggesting that different climatic variables may predict their distribution outside the cline. To address this question, we genotyped male and female house flies from nine different locations across the United States of America, and we tested if the frequencies of Y ^M^, III ^M^ and *Md-tra*^*D*^ were correlated with a variety of climatic variables.

## Materials and Methods

### House fly collections

House flies were collected from dairy farms, poultry farms, and other locations where flies are present in nine different populations across the United States of America (Supplemental Table S1) . Collections were performed in May–June 2021 using sweep nets. Flies were allowed to lay eggs in laboratories near the collection sites. Resulting pupae were then shipped to Cornell University (Ithaca, NY), where colonies from each collection site were established. Pupae from each of those colonies were then shipped to the University of Houston (Houston, TX) within 4 generations of establishing the laboratory colonies. Those pupae were raised into adults in laboratory conditions (22°C) at the University of Houston, and the emerging adults were frozen for genotyping.

### DNA extraction and genotyping

DNA was extracted from individual frozen house fly heads using the hot sodium hydroxide and tris, HotSHOT, protocol (Truett et al. 2000) . We performed PCR to test for the presence of Y ^M^ using the A12CMF1 and A12CMR1 primer pair (Hamm et al. 2009) . We were unable to design a PCR primer pair that could reliably identify the III ^M^ chromosome. We used the GM2IIIF1 and GM2IIIR2 primer pair as a positive control to confirm successful DNA amplification from males (Hamm et al. 2009) . We tested for *Md-tra*^*D*^ in females using a primer pair that amplifies a region of exon 3 of the *Md-tra* gene containing the diagnostic deletion (Hediger et al. 2010; Meisel et al. 2016) . We tested for the presence of *Mdmd* in females using the Mdmd_F1 and Mdmd_R4 primer pair (Sharma et al. 2017) .

Each male was genotyped for the presence of Y ^M^. From these data, we determined the frequencies of males with Y ^M^ and males without Y ^M^ (Figure 1A) . The latter group (males without Y ^M^) presumably consists of III ^M^ males, but some males without Y ^M^ may also carry III ^M^. We used population genetic simulations (see below) to estimate the frequency of III ^M^ along with the frequencies of all possible male genotypes in each population (Figure 1B) .

**Figure 1.**
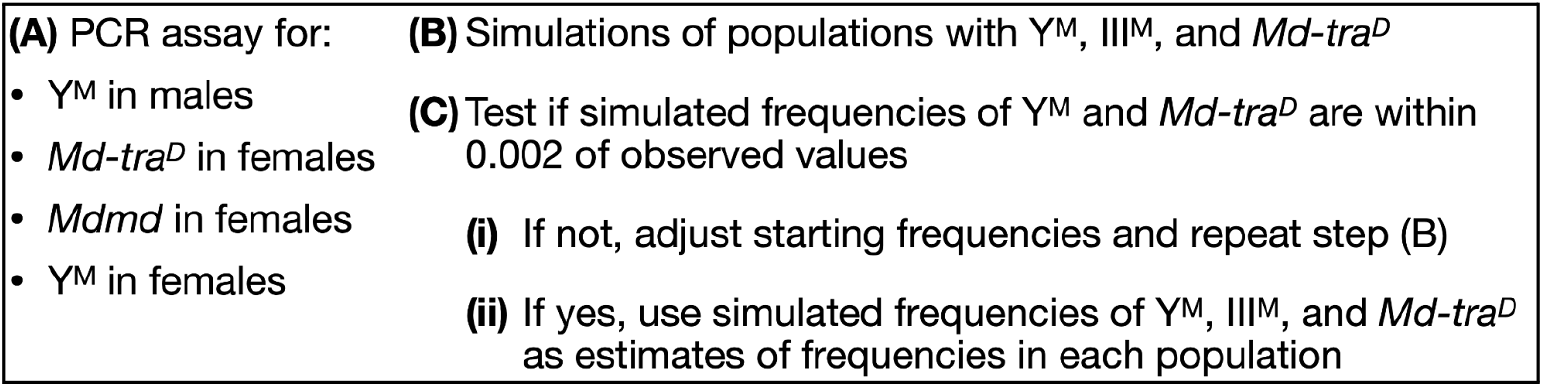
Approach to estimate the frequencies of Y ^M^, III ^M^, and *Md-tra*^*D*^ in each of the nine sampled populations. **A**. PCR was used to determine the frequencies of proto-sex chromosomes and sex determining loci/alleles. **B**. Simulations were performed to determine equilibrium frequencies of proto-sex chromosomes in randomly mating populations. **C**. If those equilibrium frequencies deviated from the observed frequencies (i), then the starting frequencies were adjusted and the simulations repeated. If the equilibrium frequencies were similar to the observed frequencies in a population (ii), then the equilibrium frequencies were used as estimates of the frequencies of Y ^M^, III ^M^, and *Md-tra*^*D*^ in a population.

Each female was genotyped for the presence of *Md-tra* ^*D*^, *Mdmd*, and Y ^M^. For each population, we determined the frequencies of: females without *Md-tra* ^*D*^; females with *Md-tra* ^*D*^ but not *Mdmd*; females with *Md-tra* ^*D*^ and Y ^M^; and females with *Md-tra* ^*D*^ and *Mdmd* but not Y ^M^ (Figure 1A) . We also used these data to determine the frequencies of *Md-tra* ^*D*^, *Mdmd* in females, and *Mdmd* in *Md-tra* ^*D*^ females.

### Population genetic simulations to determine genotype frequencies

We used population genetic simulations to predict the frequencies of 18 possible sex chromosome genotypes in each sampled population (Figure 1B) . Our PCR genotyping method is unable to detect the III ^M^ chromosome nor is it able to diagnose specific genotypes, which means our genotype data are incomplete. There are a total of 18 possible genotypes (8 male and 10 female) when considering all possible combinations of Y ^M^, III ^M^, and *Md-tra* ^*D*^ (Meisel 2021) . To help overcome the deficiency in our genotyping protocol, we performed population genetic simulations in order to identify genotype frequencies that would produce the frequencies of Y ^M^ and *Md-tra* ^*D*^ observed in our data (Figure 1C) . Our simulations used the same recursion equations that we have previously used to model randomly mating house fly populations, without selection (Meisel et al. 2016) . We performed separate simulations for each of the nine sampled populations in order to estimate the frequencies of Y ^M^, III ^M^, *Md-tra* ^*D*^, and each genotype in each population.

In the simulations, we calculated the frequency of a chromosome (Y ^M^ or III ^M^) or allele (*Md-tra* ^*D*^) as follows. The frequency of Y ^M^ was calculated as the number of Y ^M^ chromosomes in a population divided by the sum of the number of Y ^M^ and X chromosomes in that population. The frequency of the III ^M^ chromosome was calculated as the number of third chromosomes with *Mdmd* (i.e., III ^M^) divided by the total number of third chromosomes. The frequency of *Md-tra* ^*D*^ was calculated as the number of *Md-tra* ^*D*^ alleles divided by the total number of *Md-tra* genes. The frequencies of Y ^M^ and III ^M^ can take values between 0 and 1, while the *Md-tra* ^*D*^ frequency can take values between 0 and 0.25 (because it is a W chromosome).

We started each simulation with estimates of the frequencies of Y ^M^, III ^M^, and *Md-tra* ^*D*^ from our PCR assay. The initial frequency of Y ^M^ (*f*_*YM*_) was estimated as half of the frequency of males carrying a Y ^M^ chromosome. This initial frequency assumes that all males carrying Y ^M^ also carry an X chromosome, and all copies of Y ^M^ are found in heterozygous individuals. The initial frequency of III ^M^ (*f*_*IIIM*_) was estimated as 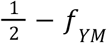, which assumes all males without Y ^M^ carry one copy of III ^M^. Both of these calculations assume that *f*_*YM*_ and *f*_*IIIM*_ are equal in males and females. The initial frequency of *Md-tra* ^*D*^ (*f*_*traD*_) was estimated as one quarter of the number of females carrying *Md-tra* ^*D*^, which requires no assumptions about genotypes because each female carrying *Md-tra* ^*D*^ must be heterozygous and males cannot carry *Md-tra* ^*D*^.

From the initial estimated frequencies of Y ^M^, III ^M^, and *Md-tra* ^*D*^, we calculated initial frequencies of each of the 18 possible genotypes. We first calculated the initial frequencies of each single chromosome genotype assuming random mating. For example, the frequency of the X/X genotype was estimated as (1 − *f*_*YM*_)^2^; the X/Y ^M^ frequency was estimated as 2*f*_*YM*_(1 − *f*_*YM*_); and the Y ^M^/Y ^M^ frequency was estimated as *f*_*YM*_^2^. Similar calculations were performed to estimate the frequencies of the third chromosome genotypes. The frequency of the *Md-tra* ^*D*^/*Md-tra*^*+*^ genotype was estimated as *f*_*traD*_(1 − *f*_*traD*_), and the frequency of the *Md-tra*^*+*^/*Md-tra*^*+*^ genotype was estimated as (1 − *f*_*traD*_)^2^. The initial frequencies of the 18 multi-chromosome genotypes were then calculated using the product of each single chromosome genotype, and each of the 18 frequencies were divided by the sum to obtain a new estimate that summed to one.

We next performed simulations for 10 generations of random mating using those initial genotype frequencies and previously developed recursion equations (Meisel et al. 2016) to determine the equilibrium frequencies of each chromosome and genotype, given the initial genotype frequencies. We compared the resulting values of *f*_*YM*_ and *f*_*traD*_ after 10 generations with the observed values measured in the respective natural population. We tested if the simulated values were within 0.002 of the observed values (Figure 1C) . If the simulated frequency of a chromosome was less than the observed frequency, we increased the initial frequency and repeated the simulation. Conversely, if the simulated frequency was greater than the observed frequency, we decreased the initial frequency and repeated the simulation. We repeated this process until the simulated frequencies of *f*_*YM*_ and *f*_*traD*_ matched the observed frequencies within 0.002. We used these simulated frequencies at equilibrium (i.e., after 10 generations) as estimates of the chromosome and genotype frequencies in downstream analyses.

### Climate data

We tested if climatic features were associated with the frequencies of sex chromosomes and genotypes across the sampled populations. To do so, we obtained weather data from the nearest NOAA station to the collection site measured between 1991–2020 (Table 1; Supplemental Table S1) . From these data, we extracted many of the same features as a previous analysis comparing the frequencies of house fly sex chromosomes and climatic data (Feldmeyer et al. 2008), and we used annual precipitation (Precip) instead of humidity measurements. We additionally calculated mean temperatures for only summer months (May–July) because that was when the house flies in our collections were sampled. All temperatures were provided in Fahrenheit, and we converted the values to Celcius for analysis.

**Table 1.** Climate features analyzed.

| Name | Description |
| --- | --- |
| $T_{\text{mean}}$ | Annual mean average daily temperature |
| $T_{\text{min}}$ | Annual mean minimum daily temperature |
| $T_{\text{max}}$ | Annual mean maximum daily temperature |
| Daily<sub>TR</sub> | Annual mean daily temperature range |
| Precip | Annual precipitation |
| $T_{\text{active}}$ | Mean temperature of warmest month |
| Season<sub>1</sub> | Coefficient of variation of average monthly temperatures |
| Season<sub>2</sub> | Average of difference between highest and lowest monthly maximum temperature and difference between highest and lowest monthly minimum temperature |
| Summer<sub>mean</sub> | Mean average monthly temperature during May-July |
| Summer<sub>max</sub> | Mean maximum monthly temperature during May-July |
| Summer<sub>min</sub> | Mean minimum monthly temperature during May-July |

We performed two separate analyses to test if climate features were associated with the frequencies of Y ^M^, III ^M^, *Md-tra* ^*D*^, males with multiple proto-Y chromosomes, and males with both Y ^M^ and III ^M^. First, we used the prcomp() function in R to perform a principal component analysis (PCA) on the annual climate data (with variables scaled to have unit variance), excluding the measurements that only sampled summer months (May–June). We then tested if each principal component (PC) is correlated or associated with Y ^M^ or III ^M^ frequencies across populations. We also calculated pairwise rank-order (Spearman) correlations between chromosome frequencies and individual climate features.

## Results

We used PCR assays to determine the frequencies of male house flies carrying the Y ^M^ chromosome across nine locations (populations) sampled in 2021 in the United State of America (Figure 2A) . We also used PCR to determine the frequencies of female house flies carrying *Md-tra* ^*D*^, *Mdmd*, and Y ^M^ in the same nine populations (Figure 2A) . We then performed population genetic simulations to identify frequencies of Y ^M^, III ^M^, and *Md-tra* ^*D*^ in each population that could produce the observed frequencies we observed in our PCR assays (Figure 1B; Supplemental Figures S1-S9). The III ^M^ chromosome was at the highest frequency in two of the three southernmost populations (CA and FL). In the FL population, males were predicted to be almost entirely III ^M^, and there were very few *Md-tra* ^*D*^ females. In contrast, the northernmost population (PA) was predicted to have almost entirely Y ^M^ males. In addition, there was a positive correlation between the predicted frequencies of males with multiple male-determining chromosomes (Y ^M^ and/or III ^M^) and females carrying *Md-tra* ^*D*^ (*r*^2^ = 0. 975, *p* = 4. 5 × 10^−7^; Figure 2C) .

**Figure 2.**
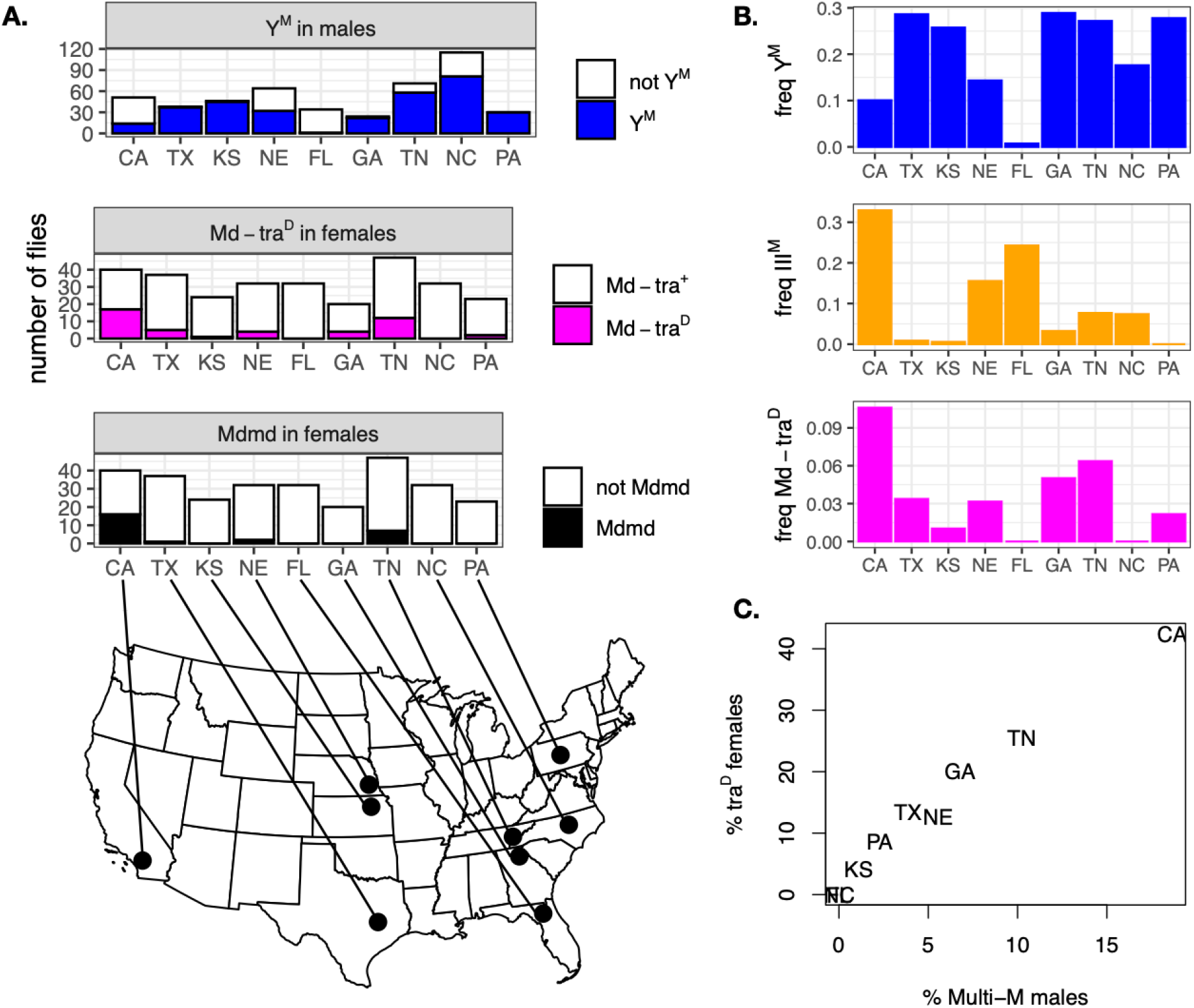
Observed and inferred frequencies of sex chromosomes, determiners, and alleles across nine populations. **A**. The number of males genotyped with Y ^M^ (blue bars), females with *Md-tra* ^*D*^ (magenta bars), and females with *Mdmd* (black bars) from each of nine populations are plotted. The number of flies without each chromosome, allele, or gene are shown in white bars. The sampling locations for each population are indicated by dots on the map. **B**. The estimated frequencies of Y ^M^, III ^M^, and *Md-tra* ^*D*^ in each of the nine populations are plotted. Estimated frequencies are from population genetics simulations which produced observed frequencies shown in panel A. **C**. The relationship between the percent of females carrying *Md-tra* ^*D*^ and the percent of males with multiple male-determining chromosomes (Y ^M^ and/or III ^M^) are plotted for the nine populations based on estimated genotype frequencies.

We compared our estimates of the frequencies of Y ^M^, III ^M^, *Md-tra* ^*D*^, and four sex chromosome genotypes in the CA and NC populations with previous measurements in nearby populations from California and North Carolina (Table 2) . A population was sampled from Chino, CA in 1982 and 2014 (Meisel et al. 2016), ∼50 km from our CA collection site (sampled in 2021). There was a high frequency of *Md-tra* ^*D*^ in both populations. However, the Chino population had a higher frequency of Y ^M^ chromosomes, while the CA population we sampled in 2021 had a higher III ^M^ frequency. In contrast, we observed similar frequencies of Y ^M^, III ^M^, and *Md-tra* ^*D*^ in the NC population we sampled in 2021 and the populations sampled in 2002, 2006, and 2007. All of the NC populations were sampled in close proximity within Wake County.

**Table 2.**
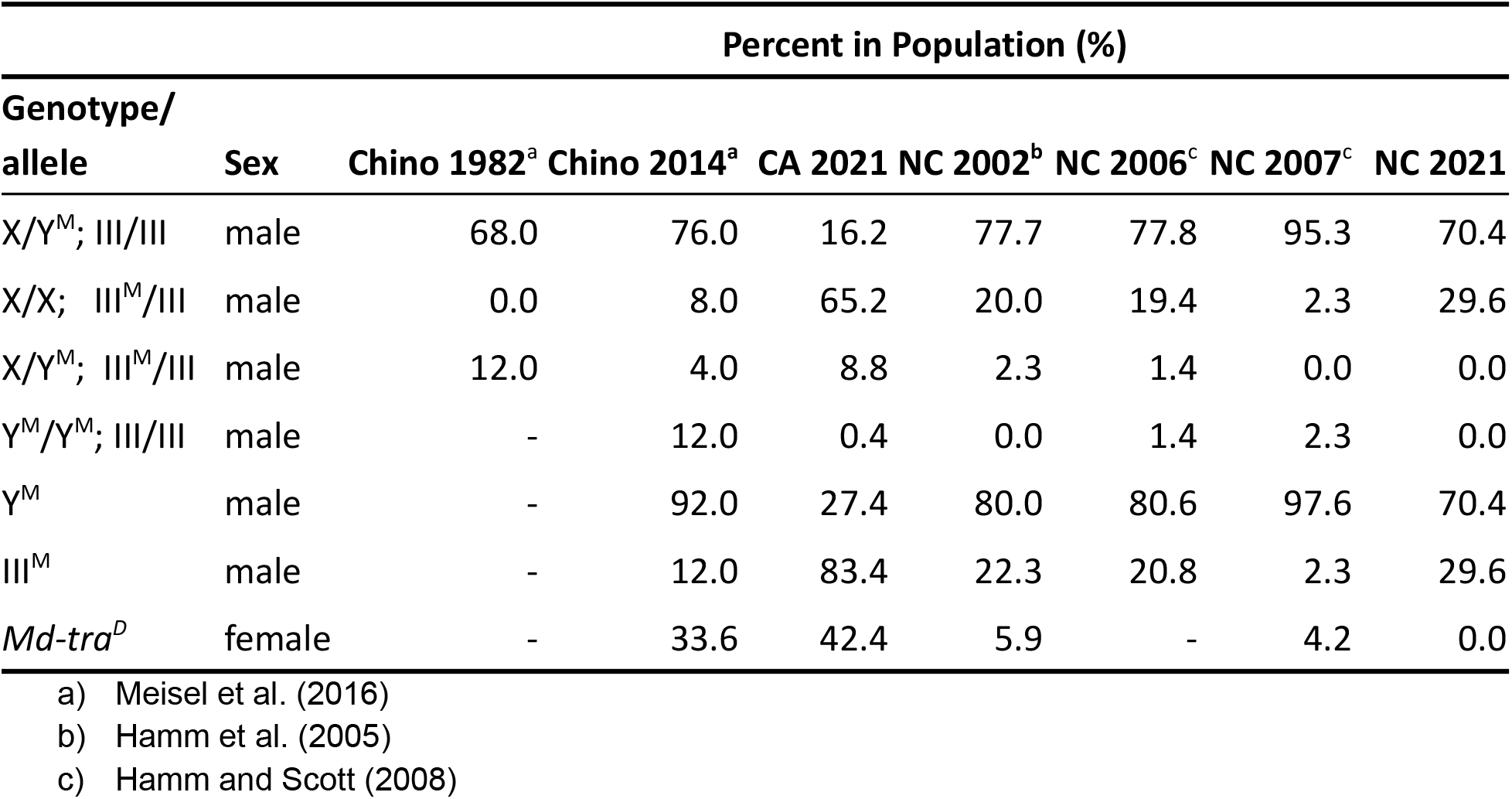

We tested for associations between climatic variables and the frequencies of sex chromosomes across the nine populations we sampled. To those ends, we first performed a principal component analysis (PCA) using eight climate features measured across the nine populations. The first three PCs explain >98% of the variance in the data (Supplemental Table S2), with PC1 and PC2 explaining nearly 90% of variance (Figure 3A) . PC1 captured variation in seasonality (Season_1_ and Season_2_), along with minimum, maximum, and average temperatures (T_mean_, T_min_, T_max_, and T_active_) across populations. PC2 captured variation in precipitation (Precip) and annual mean daily temperature range (Daily_TR_) across populations. We tested for correlations between PC1 or PC2 and the frequencies of sex chromosomes across populations. Only PC2 and Y ^M^ frequency had a significant correlation, with Y ^M^ frequency decreasing as PC2 values increased (*ρ* = -0.767, *p* = 0.0214). We additionally constructed linear models in which we tested if PC1, PC2, and their interaction predicted the frequencies of Y ^M^ or III ^M^. The only significant relationship in these models was between PC2 and III ^M^ frequency (*F* = 8.372, *p* = 0.034). Therefore, there is evidence that Y ^M^ and III ^M^ frequencies across populations were associated with PC2, which captured variation in precipitation and daily temperature range.

**Figure 3.**
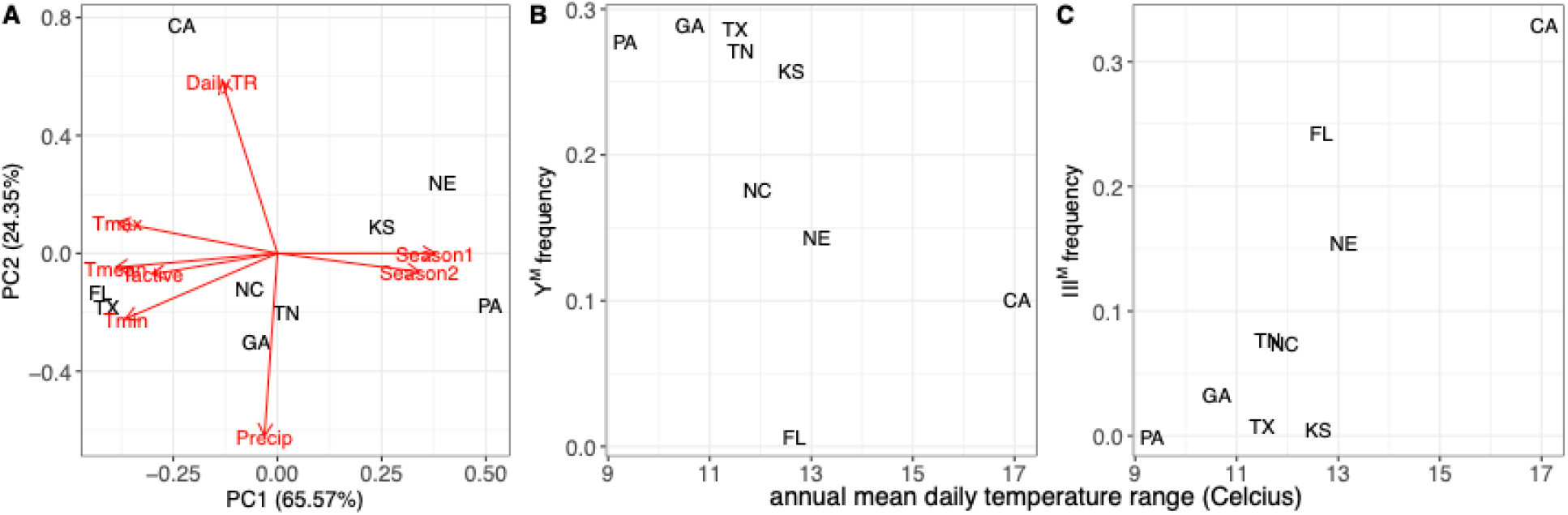
Associations between climate features and proto-Y chromosome frequencies across the nine sampled populations. Each population is represented by the two letter abbreviation of the state from which it was collected. **A**. Populations are plotted according to the first two principal components (PCs) based on climate features. The loadings of each climate feature are indicated by labeled red vectors. Vector labels are described in Table 1. **B-C**. The relationships between the predicted frequency of Y ^M^ or III ^M^ and the annual mean daily temperature range are plotted for each population.

We observed similar patterns when we calculated pairwise correlations between climatic features and sex chromosome frequencies (Supplemental Table S3). The only significant correlations were between the frequencies of Y ^M^ or III ^M^ and the annual mean daily temperature range (Figure 3C) . Specifically, the frequency of Y ^M^ was negatively correlated with the daily temperature range (*ρ =* -0.883, *p* = 0.003), and the frequency of III ^M^ was positively correlated with the daily temperature range (*ρ =* 0.783, *p* = 0.017). These results provide consistent evidence that daily temperature range is associated with proto-Y chromosome frequencies.

The CA population appeared to be an outlier in many respects, which could have driven some of the patterns we observed. For example, the CA population had higher frequencies of III ^M^ chromosomes, males with multiple male determiners, and females with *Md-tra* ^*D*^, when compared to all other populations (Figure 2) . In addition, the CA site was an outlier along PC2 because it had a higher daily temperature range and lower precipitation than the other populations (Figure 3) . When we excluded the CA site from our climate PCA, we observed similar loadings of the climate variables: PC1 explained 71.69% of the variance and captured variation in seasonality and minimum/maximum temperature, while PC2 explained 22.13% of variance and captured variation in daily temperature range and precipitation (Supplemental Figure S10) . We also observed a significant negative correlation between daily temperature range and Y ^M^ frequency (*ρ =* -0.881, *p* = 0.007) when the CA population was excluded. Therefore, the relationships between climate features and proto-Y chromosome frequencies was not driven solely by the CA population.

## Discussion

We observed substantial variation in the frequencies of Y ^M^, III ^M^, and *Md-tra* ^*D*^ across populations of house flies in North America (Figure 2) . Along the eastern coast, Y ^M^ was most common in the north (PA), III ^M^ was most common in the south (FL), and both Y ^M^ and III ^M^ were found in the central (NC) population, consistent with the previously documented cline (Hamm et al. 2005) . However, moving west, we found that the clinal distribution eroded, and latitude was not associated with the frequencies of Y ^M^ and III ^M^. For example, the GA, TN, NE, and CA populations all had moderate to high frequencies of III ^M^, Y ^M^, and *Md-tra* ^*D*^, without any relationship to latitude. In addition, the presence of all three chromosomes/alleles in CA is consistent with previous observations (Meisel et al. 2016) .

We used population genetic simulation models to estimate the frequencies of Y ^M^, III ^M^, *Md-tra* ^*D*^, and all 18 genotypes in each population based on PCR assays for the presence of Y ^M^, *Mdmd*, and *Md-tra* ^*D*^ from individual flies (Figure 1) . Our PCR assays likely do not measure allele, chromosome, and genotype frequencies as accurately as more direct genotyping assays that were used in prior studies (e.g., Hamm et al. 2005; Feldmeyer et al. 2008; Meisel et al. 2016) . In addition, we sampled flies after multiple generations of lab breeding, which could lead to deviations from the frequencies in natural populations. Nonetheless, we believe that our estimates of allele, chromosome, and genotype frequencies were sufficiently accurate for the analyses we performed. First, our estimated frequencies of Y ^M^, III ^M^, and *Md-tra* ^*D*^ are largely concordant with prior estimates from the same county in North Carolina (Table 2) . Second, previous work found that Y ^M^ is more common than III ^M^ in Texas (McDonald et al. 1975), consistent with our results (Figure 2) . Furthermore, we predicted a positive correlation between the frequencies of *Md-tra* ^*D*^ and males with multiple male-determining chromosomes (Figure 2C), as expected if Y ^M^, III ^M^, and *Md-tra* ^*D*^ frequencies are under selection to maintain balanced sex-ratios (Meisel et al. 2016) .

Despite the concordance with prior results, there are discrepancies between the frequencies we predicted and those previously observed in California. The CA population we sampled was estimated to have a high frequency of III ^M^ (Figure 2B), but previous collections from a nearby population found that Y ^M^ was at a higher frequency than III ^M^ (Meisel et al. 2016) . This discrepancy between our observations and prior measurements from California can likely be explained by a >50 km distance between our site and the previous sampling locations. Similar differences in the frequencies of house fly male-determining chromosomes have been observed over relatively short distances in Japan and Spain (Tomita and Wada 1989; Li et al. 2022) . Therefore, small-scale variations in Y ^M^ and III ^M^ frequencies appear to be a global phenomenon, in addition to the large-scale variation observed across the entire continent (Figure 2) .

Our results contribute to the body of evidence that climatic factors affect the frequencies of Y ^M^ and III ^M^ in natural populations of house fly. In addition to the clinal distribution we confirmed in eastern North America (Figure 2), we also detected consistent associations between Y ^M^ or III ^M^ frequencies and the annual mean daily temperature range, Daily_TR_ (Figure 3) . This climate metric captures the extent of temperature heterogeneity within days, averaged over the entire year. We predicted that Y ^M^ was at the highest frequency when daily temperature heterogeneity was lowest, and III ^M^ frequency was higher with more daily temperature heterogeneity. The same general patterns held when the CA population, which had extreme values relative to other populations, was excluded.

Our results differ from a prior observation that the frequencies of non-Y ^M^, male-determining chromosomes (e.g., III ^M^) in Africa and Europe were higher when seasonality in temperature was highest (Feldmeyer et al. 2008) . Feldmeyer et al. (2008) measured seasonality as the difference between the minimum and maximum values of the monthly minimum and maximum temperatures. We found this measure of seasonality was orthogonal to daily temperature range (Figure 3A) and not significantly correlated with III ^M^ or Y ^M^ frequency (Supplemental Table S3). However, both seasonality and daily temperature range are measures of temperature heterogeneity across time, suggesting that temperature variation more generally may be an important selection pressure that affects proto-Y chromosome frequencies across house fly populations.

There is growing evidence that ecological factors contribute to sex chromosome evolution (Meisel 2022) . Our results contribute to evidence that temperature variation across the species range predicts the frequencies of house fly proto-Y chromosomes (Franco et al. 1982; Denholm et al. 1986; Tomita and Wada 1989; Hamm et al. 2005; Feldmeyer et al. 2008) . In addition, the proto-Y chromosomes affect thermal traits in ways that are consistent with their clinal distributions (Delclos et al. 2021) . These temperature-dependent phenotypic effects likely create variation in the fitness effects of the proto-Y chromosomes, which in turn allow for the maintenance of multiple male-determining loci across populations. This is a special case of local adaptation maintaining genetic variation across populations (Wadgymar et al. 2022) . It may also be possible for these differences in fitness effects to promote divergence between populations and subsequent speciation. These links between ecological adaptation and proto-Y chromosomes could therefore provide a mechanism to explain the disproportionate effects of sex chromosomes on speciation (Payseur et al. 2018) .

It remains unclear if or how ecological selection pressures that affect sex chromosome evolution are related to sex-specific selection pressures that are predicted to be important for sex chromosome evolution. The house fly proto-Y chromosomes are disproportionately found in males relative to females, but they can also be carried by females who have an *Md-tra* ^*D*^ allele. Population genetic models predict that the proto-Y chromosomes could have male-beneficial, female detrimental sexually antagonistic fitness effects, which could contribute to the maintenance of the polymorphism within populations (Meisel et al. 2016; Meisel 2021) . However, there is no direct evidence for sexually antagonistic effects of the proto-Y chromosomes, let alone sexual antagonism that depends on temperature or any other ecological factor. Future work is therefore needed to evaluate if the well-documented temperature-dependent fitness effects of house fly proto-Y chromosomes have any relationship to their hypothesized sexually antagonistic effects. Such evidence would provide an important link between the effects of sexual antagonism and ecological variation on sex chromosome evolution.

## Acknowledgements

This material is based upon work supported by the National Science Foundation under Grant No. DEB-1845686. This research was supported in part by the U.S. Department of Agriculture under multistate agreement S-1076.

## Supplemental Material

**Supplemental Table S1.**
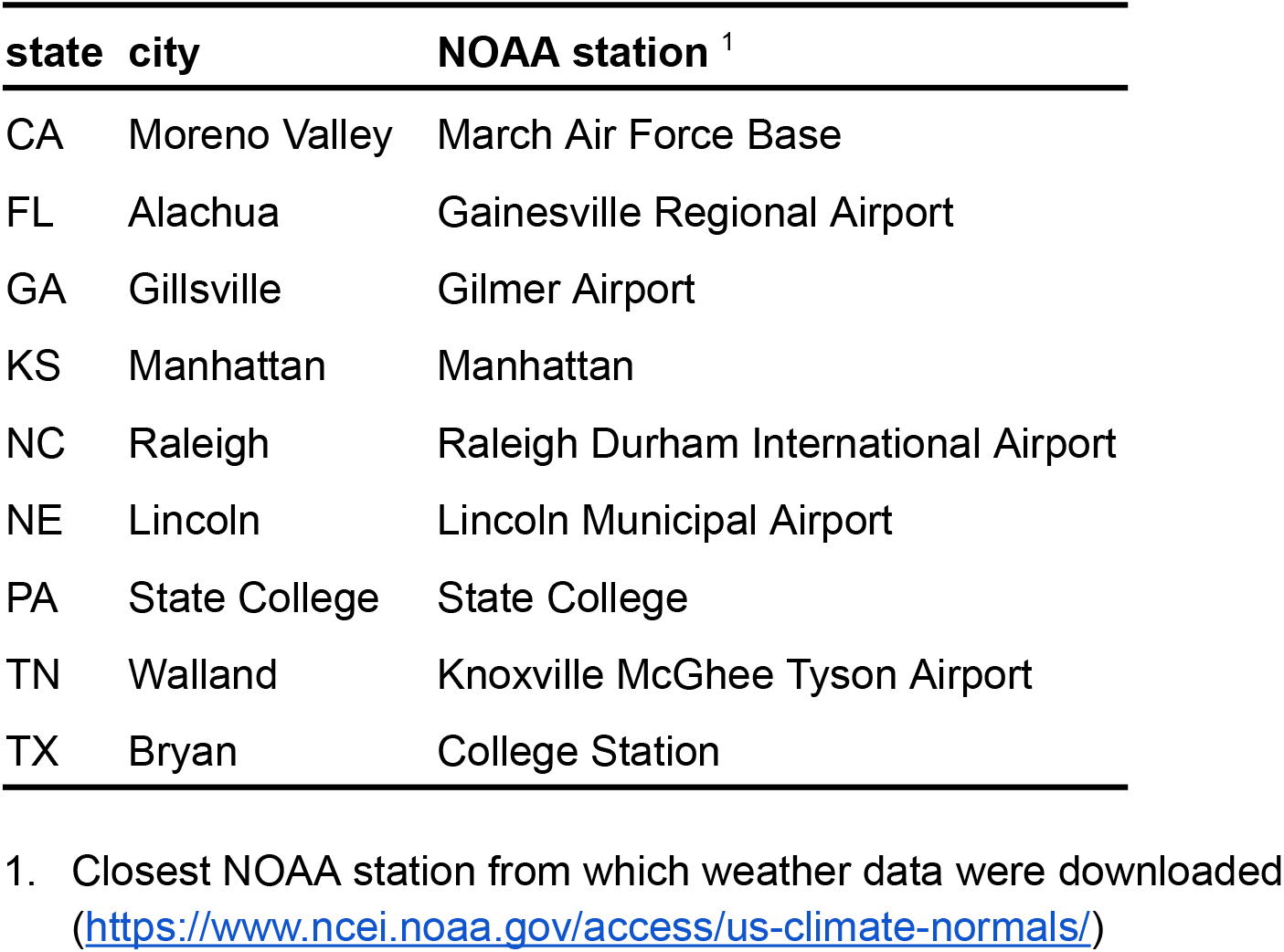
Collection sites for house flies and closest NOAA station

| state | city | NOAA station <sup>1</sup> |
| --- | --- | --- |
| CA | Moreno Valley | March Air Force Base |
| FL | Alachua | Gainesville Regional Airport |
| GA | Gillsville | Gilmer Airport |
| KS | Manhattan | Manhattan |
| NC | Raleigh | Raleigh Durham International Airport |
| NE | Lincoln | Lincoln Municipal Airport |
| PA | State College | State College |
| TN | Walland | Knoxville McGhee Tyson Airport |
| TX | Bryan | College Station |

1. Closest NOAA station from which weather data were downloaded
(https://www.ncei.noaa.gov/access/us-climate-normals/)

**Supplemental Figures S1–S9**. Simulation results to predict chromosome and genotype frequencies in each of the nine sampled populations. Graphs show the frequencies of males carrying III ^M^ (orange M), males carrying Y ^M^ (blue Y), females carrying *Md-tra* ^*D*^ (magenta D), and males (black m) across ten generations of the simulation. Dashed blue and magenta lines show the observed frequencies of males carrying Y ^M^ and females carrying *Md-tra* ^*D*^, respectively. Only the results of the final simulation that accurately predicted the observed chromosome frequencies are shown. Code to generate graphs from intermediate simulations is provided in the Supplemental Material.

**Supplemental Figure S10.**
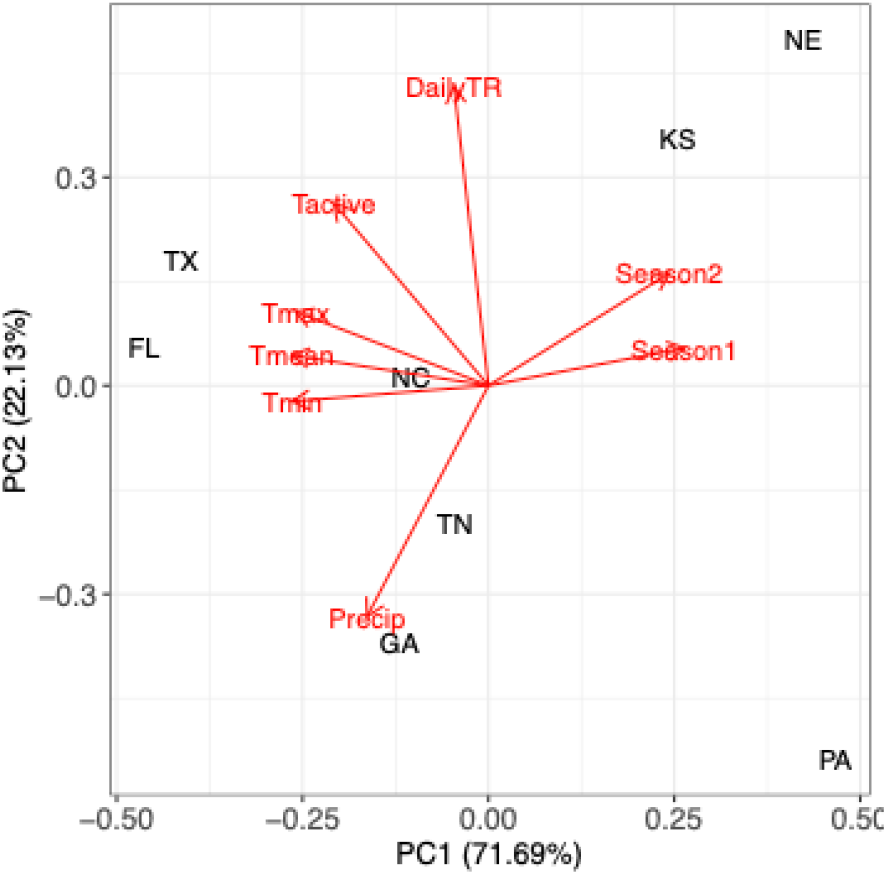
Principal component analysis (PCA) of climate features across sampling locations. Populations are plotted according to the first two principal components (PCs) based on climate features. The loadings of each climate feature are indicated by labeled red vectors. Vector labels are described in Table 1. Each population is represented by the two letter abbreviation of the state from which it was collected.

