## Supplemental Figures S1-9 for "Frequencies of house fly proto-Y chromosomes across populations are predicted by temperature heterogeneity within populations": _readme.docx

Graphs show the frequencies of Y^M^ (blue Y), III^M^ (orange M), *Md-tra^D^* (magenta D), and males (black m) across 10 generations in a simulation that predicts the frequencies of Y^M^ and *Md-tra^D^* observed in a natural population. Observed frequencies of Y^M^ and *Md-tra^D^* are shown with dashed lines of the same color as the simulated values, respectively.
