## Supplementary figures and images for "Frequencies of house fly proto-Y chromosomes across populations are predicted by temperature heterogeneity within populations"

### CA_simulation.pdf

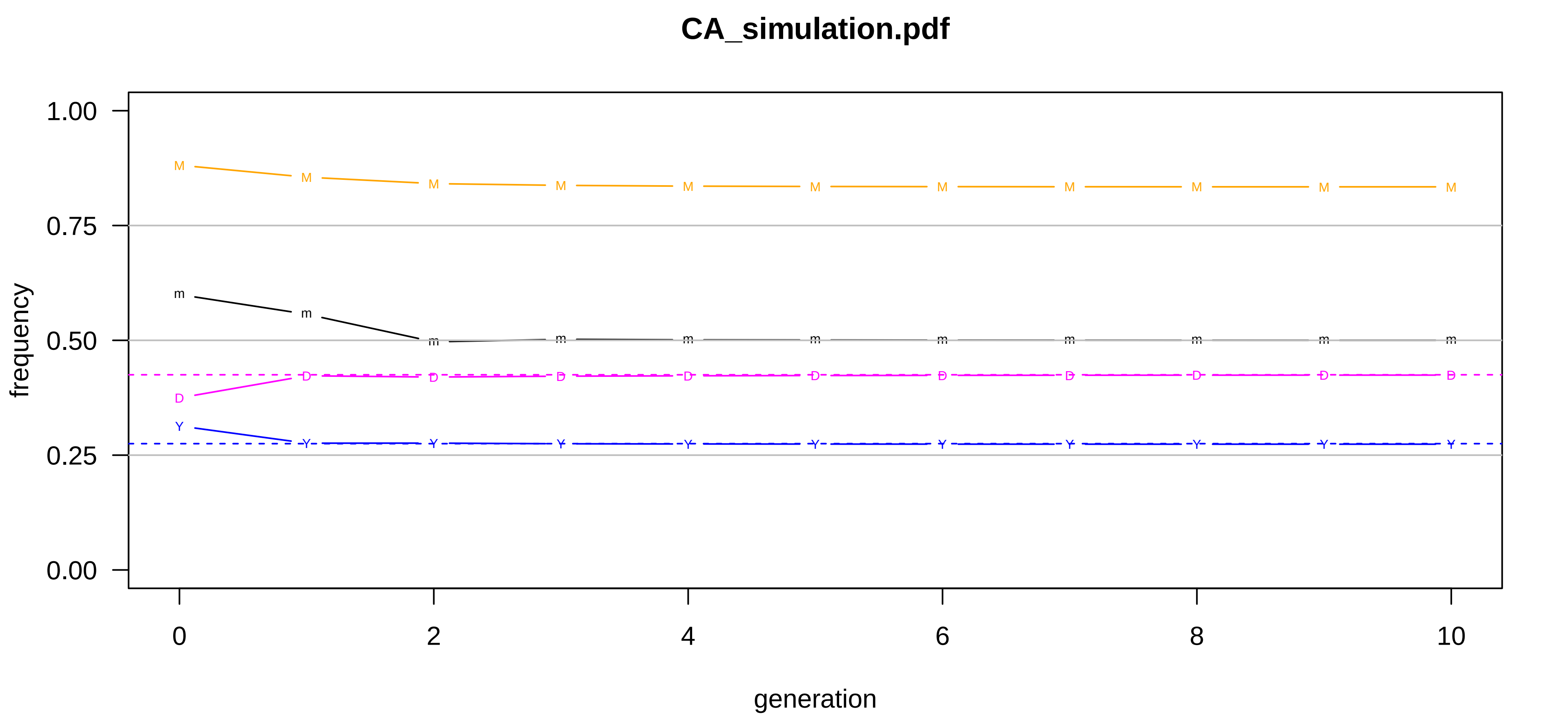

### FL_simulation.pdf

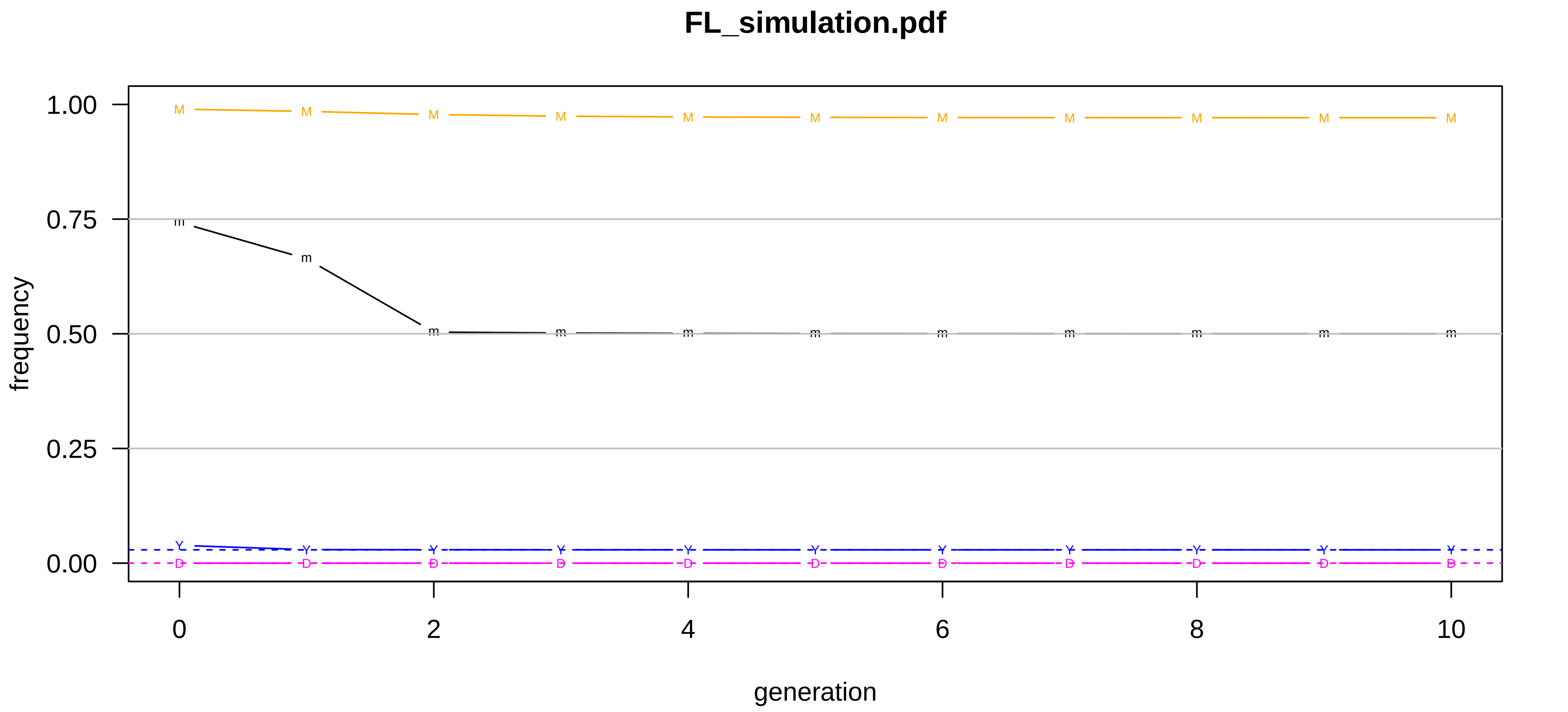

### GA_simulation.pdf

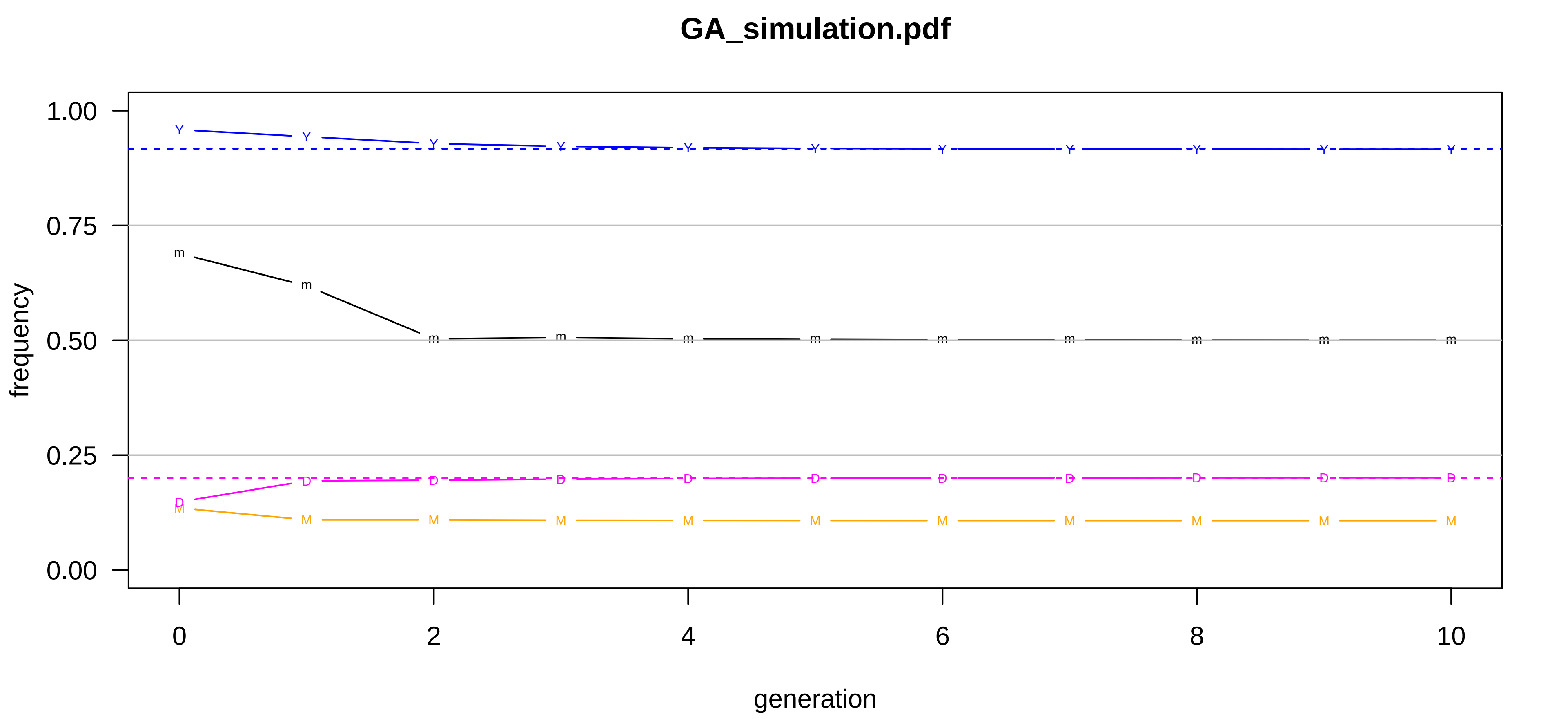

### KS_simulation.pdf

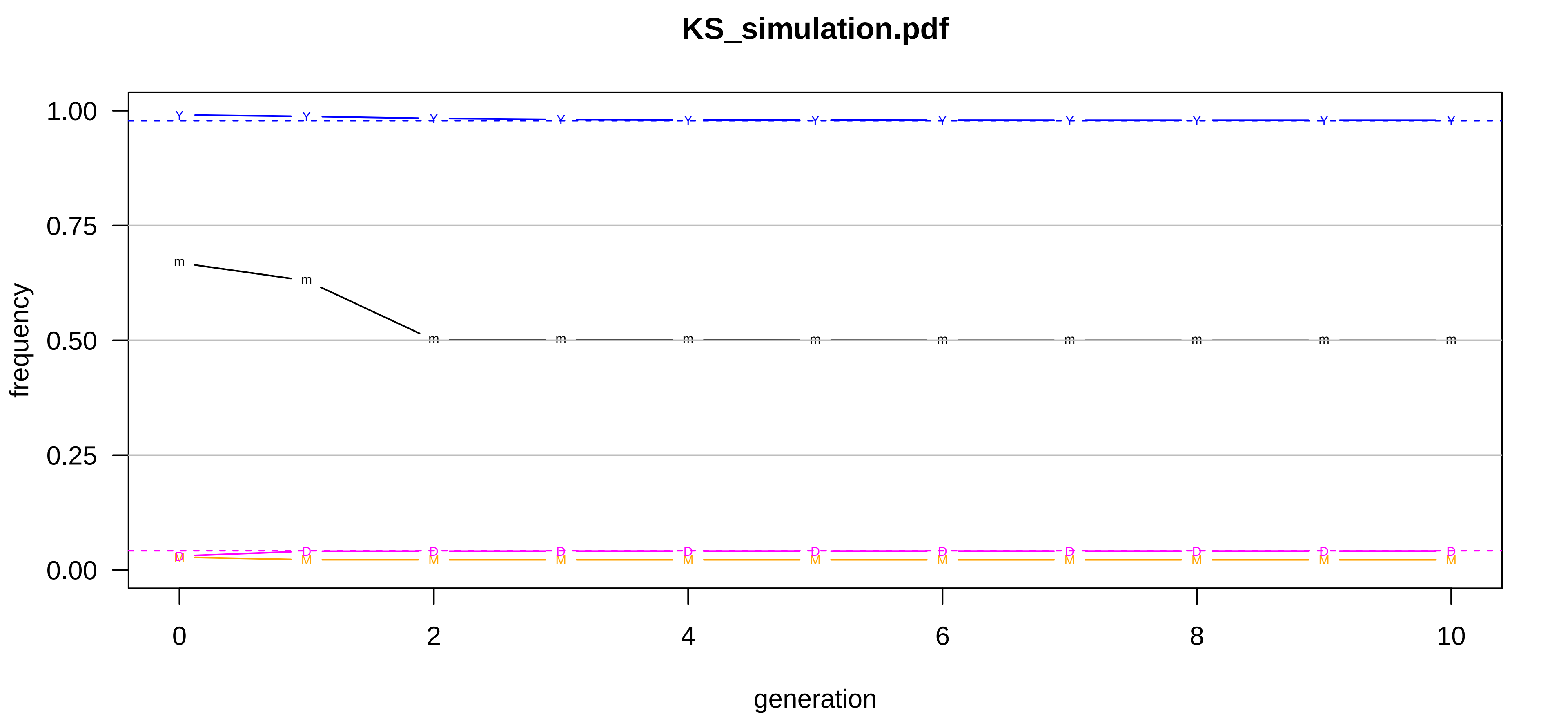

### NC_simulation.pdf

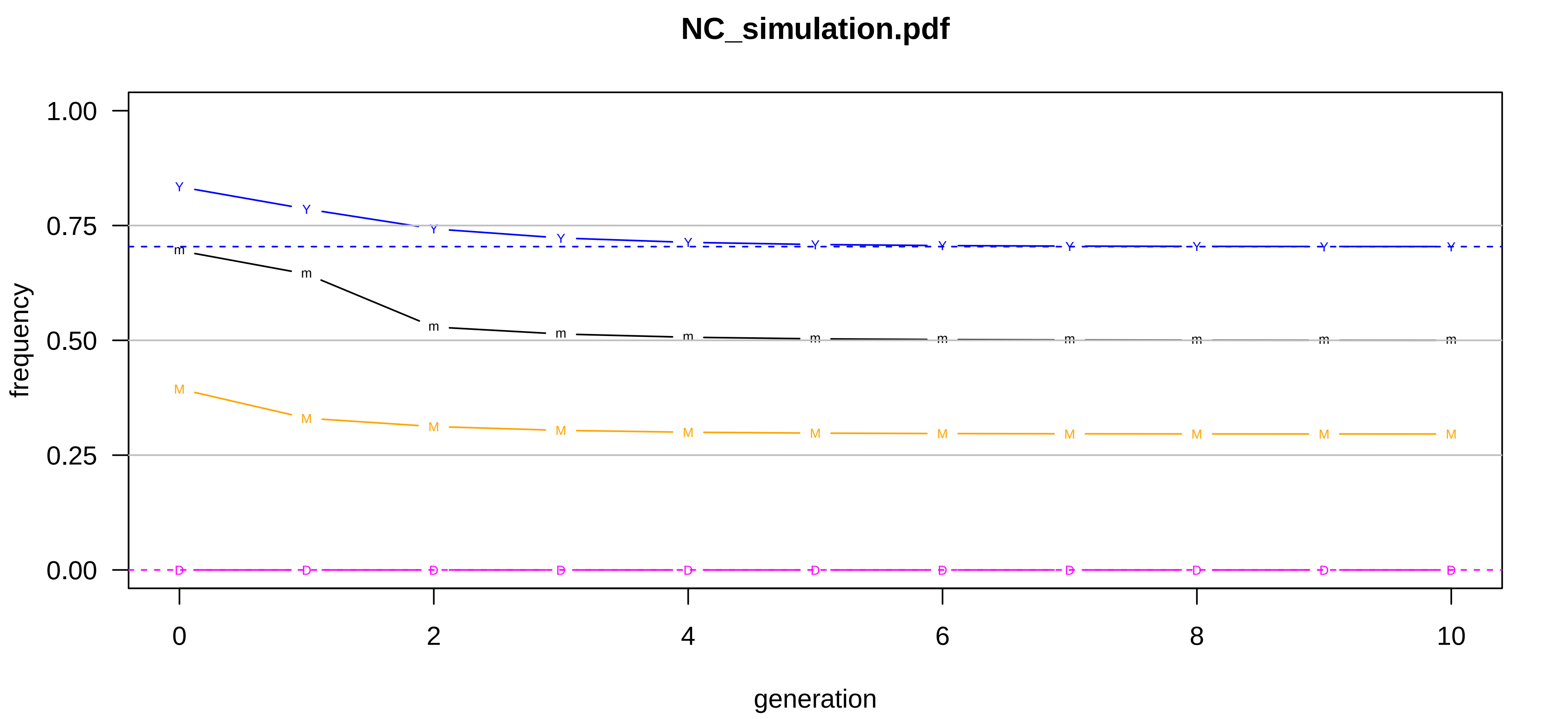

### NE_simulation.pdf

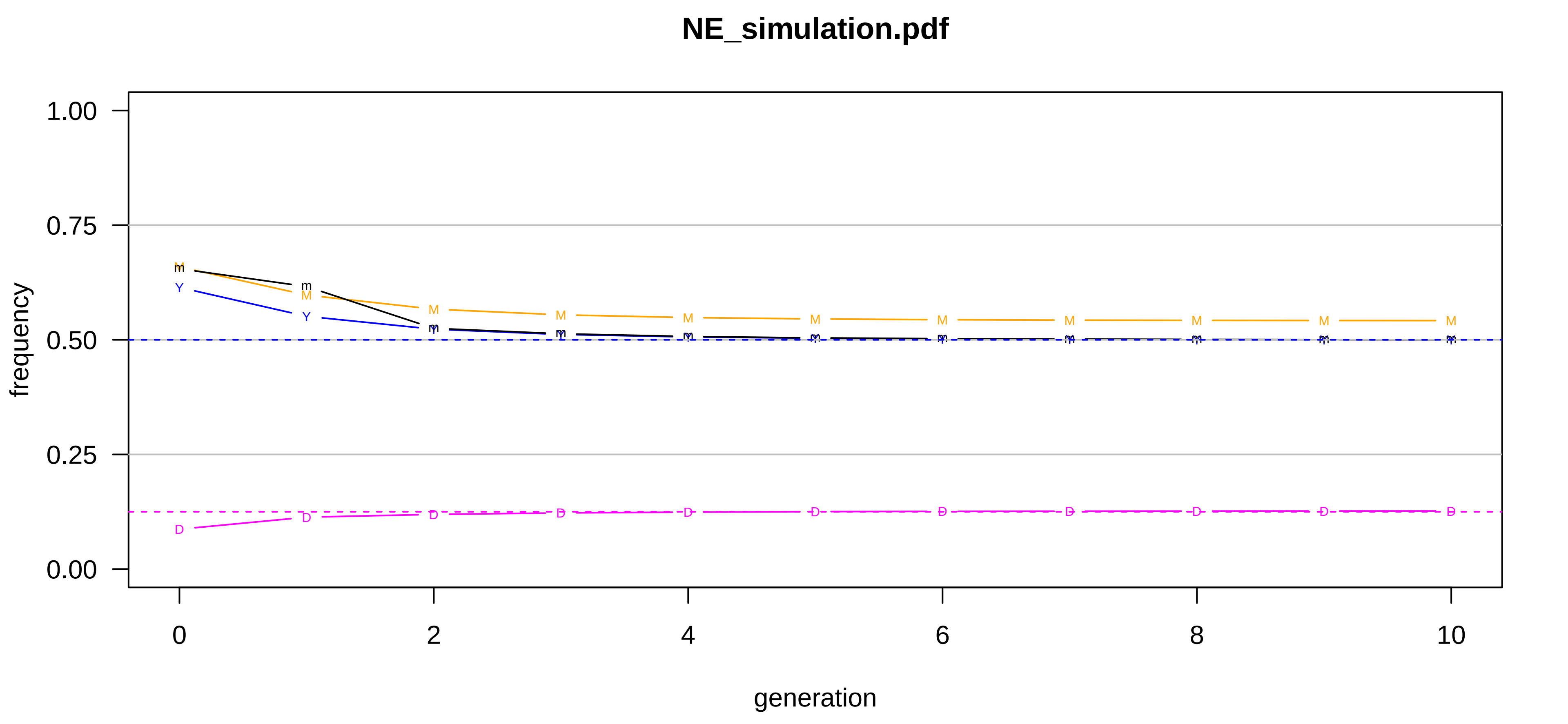

### PA_simulation.pdf

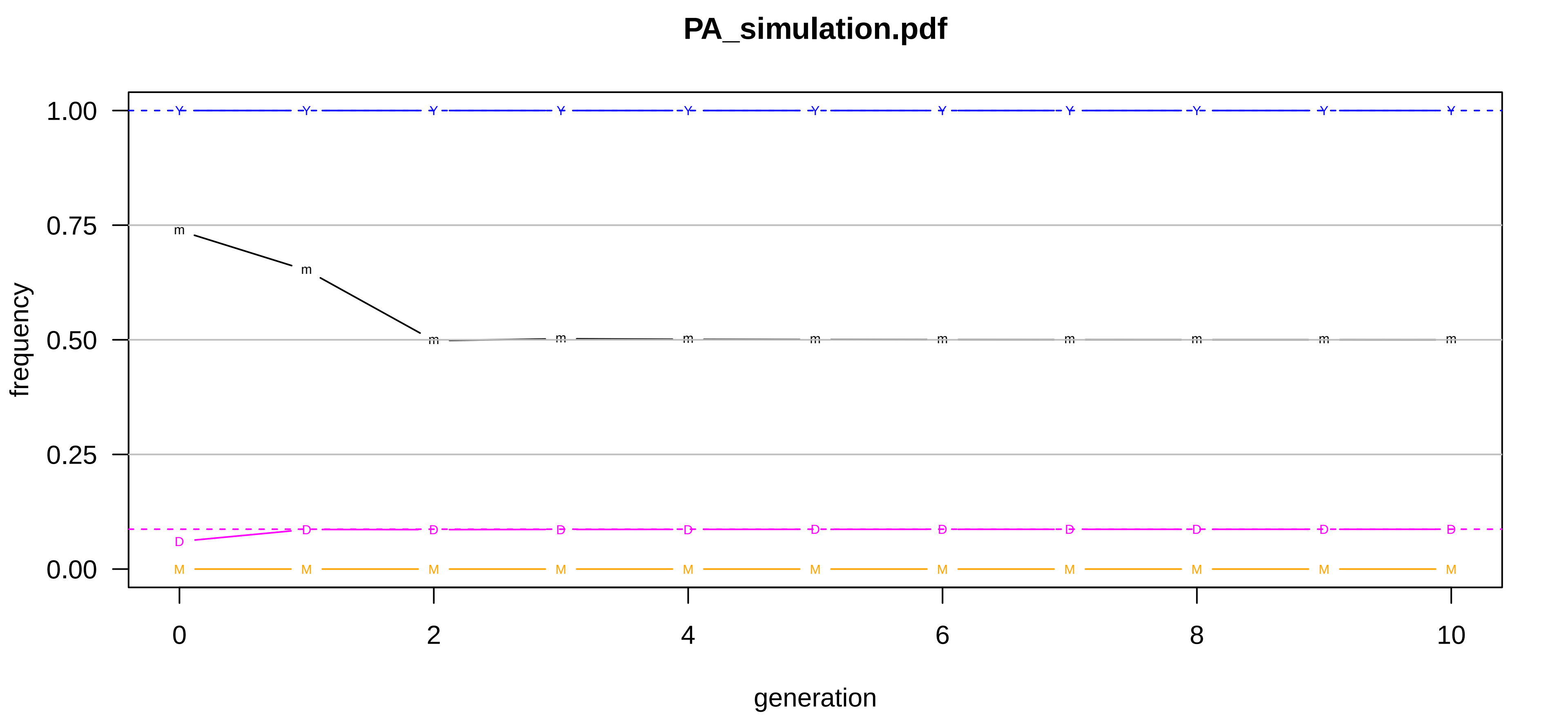

### TN_simulation.pdf

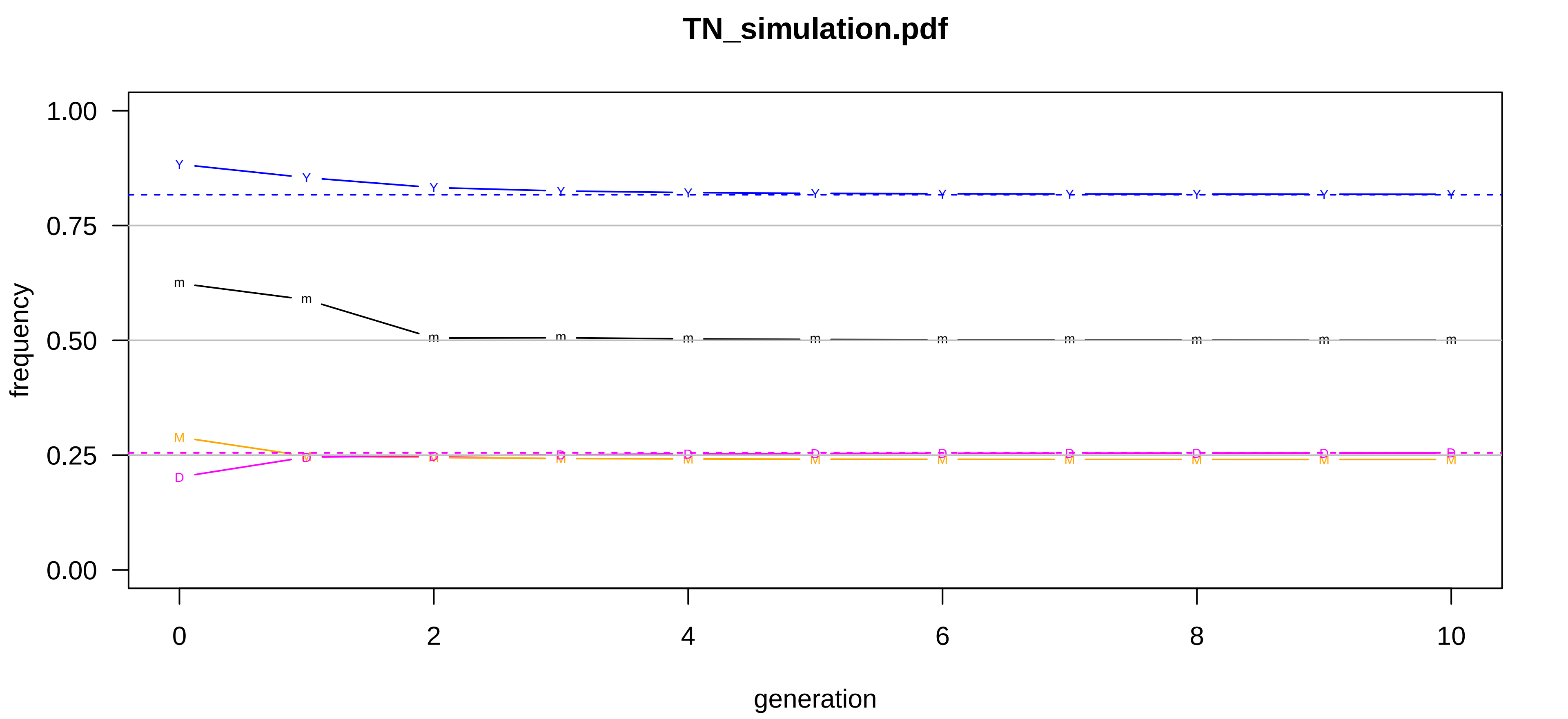

### TX_simulation.pdf

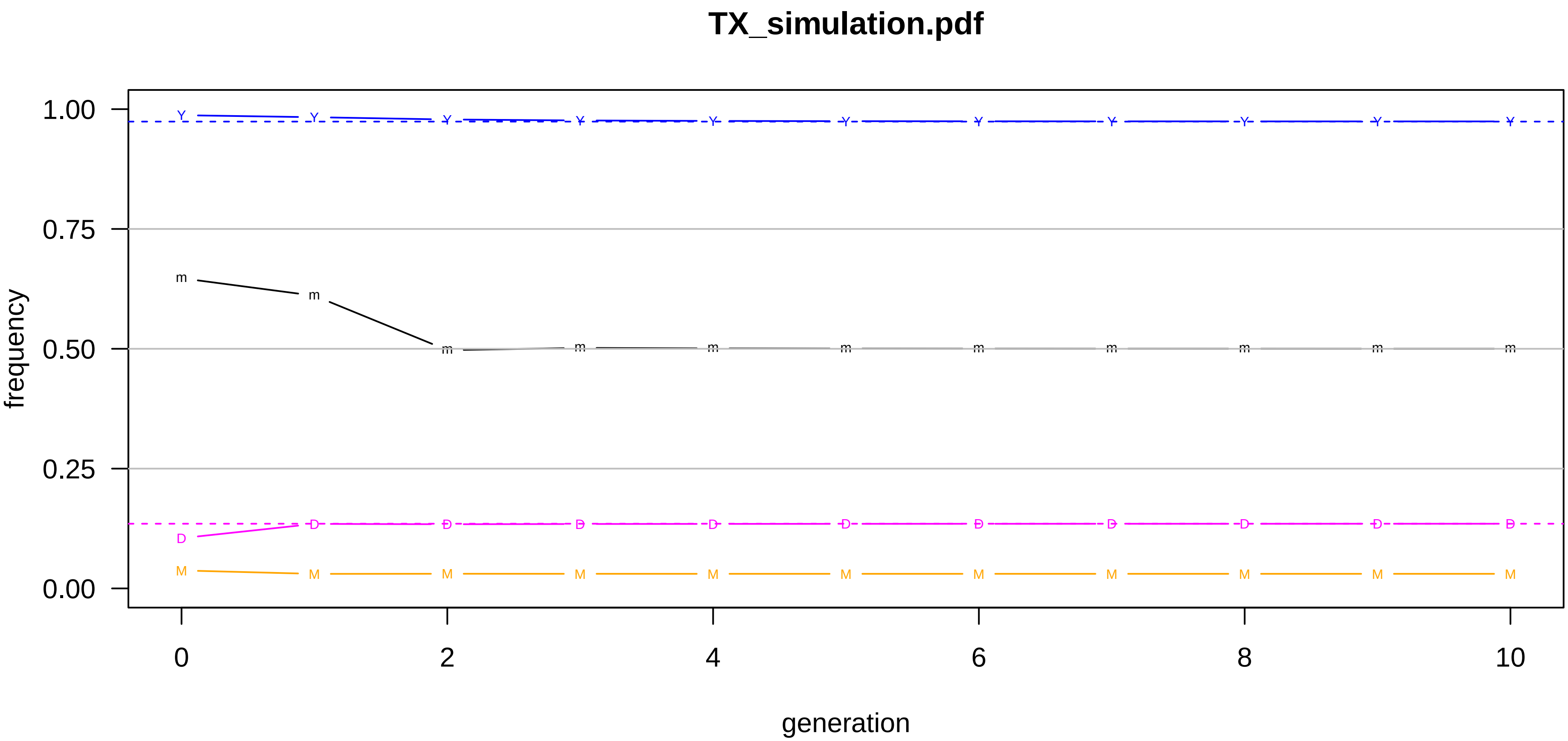
