## Supplemental Code for "Frequencies of house fly proto-Y chromosomes across populations are predicted by temperature heterogeneity within populations": SupCode_2024_04.R

```
setwd("./SupData/")
male_freqs <- read.delim("male-freqs.tsv")
female_freqs <- read.delim("female-freqs.tsv")

male_freqs$NotYM[is.na(male_freqs$NotYM)] <- 0
female_freqs$traDMnotY[is.na(female_freqs$traDMnotY)] <- 0
female_freqs$traDYM[is.na(female_freqs$traDYM)] <- 0
female_freqs$traDnoM[is.na(female_freqs$traDnoM)] <- 0
female_freqs$traD[is.na(female_freqs$traD)] <- 0


library(ggplot2)
# code states by west to east order using factor
states <- c("CA", "TX", "KS", "NE", "FL", "GA", "TN", "NC", "PA")

#male data
male_counts <- data.frame(
	state = factor(male_freqs$Pop, levels=states), # order states as factors sorted by west to east
	YM = c(rep("YM",length(male_freqs$YM)), rep("not YM",length(male_freqs$NotYM))), 
	counts = c(male_freqs$YM, male_freqs$NotYM)
)

#female data
female_counts <- data.frame(
	state = factor(female_freqs$Pop, levels=states), # order states as factors sorted by west to east
	gene = c(rep("tra",2*length(female_freqs$Pop)), rep("Mdmd",2*length(female_freqs$Pop))), 
	allele = factor(
		c(rep("tra^D",length(female_freqs$Pop)), rep("tra+",length(female_freqs$Pop)), rep("Mdmd",length(female_freqs$Pop)), rep("not Mdmd",length(female_freqs$Pop))), 
		levels=c("not Mdmd", "Mdmd", "tra+", "tra^D")
	), 
	counts = c(
		female_freqs$traD, 
		female_freqs$tra,
		female_freqs$traDYM + female_freqs$traDMnotY, 
		female_freqs$tra + female_freqs$traDnoM
	)
)


###########
# Plot collection sites on map
library(usmap)
library(sf)
library(sp)
library(cowplot)

collection_sites <- data.frame(
	Pop = male_freqs$Pop, 
	lat = c(33.858630, 29.7946429, 34.273960, 39.186691, 35.722480, 40.813618, 40.7944504, 35.772283, 30.67), 
	lon = c(-117.020450, -82.4102420, -83.594450, -96.566597, -78.680820, -96.702599, -77.8616386, -83.851272, -96.35)
)
transformed_data <- usmap_transform(collection_sites)

sex_counts <- rbind(
	data.frame(
		sex="male", 
		state = male_counts$state, 
		gene = "XY", 
		allele = male_counts$YM, 
		counts = male_counts$counts
	),
	cbind(sex="female", female_counts)
)
sex_counts$facets <- c(
	gl(1, 18, labels=expression(Y^M~"in"~males)),
	gl(1, 18, labels=expression(Md-tra^D~"in"~females)), 
	gl(1, 18, labels=expression(Mdmd~"in"~females))
)
sex_counts$labs <- factor(sex_counts$allele, levels=c("not YM", "YM", "tra+", "tra^D", "not Mdmd", "Mdmd" )) # order levels for legend

sex_counts_p <- ggplot(sex_counts, aes(fill=labs, x=state, y=counts)) + 
	geom_bar(stat="identity",position="stack", color="black") + 
	scale_y_continuous("number of flies") + 
	scale_fill_manual(labels=c(expression(not~Y^M), expression(Y^M), expression(Md-tra^"+"), expression(Md-tra^D), expression(not~Mdmd), expression(Mdmd)), values = c("YM"="blue", "not YM"="white", "tra+"="white", "tra^D"="magenta", "Mdmd"="black", "not Mdmd"="white")) +
	scale_x_discrete("") + 
	facet_wrap(~facets, ncol=1, labeller = "label_parsed", scales="free_y") +  
	theme_bw() + 
	theme(legend.title=element_blank(), legend.text.align = 0) 

USMap_p <- plot_usmap(regions = "states", exclude=c("AK","HI")) + 
	geom_sf(
  		data = transformed_data,
  		color = "black", size=3)
#  geom_point(data = transformed_data, color = "black", size = 3) +
#  theme(panel.background=element_blank())


# Make with cowplot so that can have separate legends for each assay/sex 
male_counts_p <- ggplot(subset(sex_counts, sex=="male"), aes(fill=labs, x=state, y=counts)) + 
	geom_bar(stat="identity",position="stack", color="black") + 
	scale_fill_manual(labels=c(expression(not~Y^M), expression(Y^M)), values = c("YM"="blue", "not YM"="white")) +
	scale_y_continuous("") + 
	scale_x_discrete("") + 
	facet_wrap(~facets, ncol=1, labeller = "label_parsed", scales="free_y") +  
	theme_bw() + 
	theme(legend.title=element_blank(), legend.text.align = 0, plot.margin = unit(c(0.1, 0, 0, 0.1), "cm")) #, axis.text.x=element_blank(), axis.ticks.x=element_blank()

female_counts_p1 <- ggplot(subset(sex_counts, gene=="tra"), aes(fill=labs, x=state, y=counts)) + 
	geom_bar(stat="identity",position="stack", color="black") + 
	scale_fill_manual(labels=c(expression(Md-tra^"+"), expression(Md-tra^D)), values = c("tra+"="white", "tra^D"="magenta")) +
	scale_y_continuous("number of flies") + 
	scale_x_discrete("") + 
	facet_wrap(~facets, ncol=1, labeller = "label_parsed", scales="free_y") +  
	theme_bw() + 
	theme(legend.title=element_blank(), legend.text.align = 0, plot.margin = unit(c(0.1, 0, 0, 0.1), "cm")) #, axis.text.x=element_blank(), axis.ticks.x=element_blank(), plot.margin = unit(c(0, 0, 0, 0), "cm"))) 

female_counts_p2 <- ggplot(subset(sex_counts, gene=="Mdmd"), aes(fill=labs, x=state, y=counts)) + 
	geom_bar(stat="identity",position="stack", color="black") + 
	scale_fill_manual(labels=c(expression(not~Mdmd), expression(Mdmd)), values = c("not Mdmd"="white", "Mdmd"="black")) +
	scale_y_continuous("") + 
	scale_x_discrete("") + 
	facet_wrap(~facets, ncol=1, labeller = "label_parsed", scales="free_y") +  
	theme_bw() + 
	theme(legend.title=element_blank(), legend.text.align = 0, plot.margin = unit(c(0.1, 0, 0, 0.1), "cm")) #, plot.margin = unit(c(0, 0, 0.2, 0), "cm"))) 


pdf("map_counts_v2.pdf", width=4, height=6)
ggdraw(plot_grid(male_counts_p, female_counts_p1, female_counts_p2, USMap_p, ncol = 1, rel_heights=c(1,1,1,2))) + 
  draw_line(x=c(0.15, 0.18), y=c(0.43, 0.15)) + # CA
  draw_line(x=c(0.22, 0.53), y=c(0.43, 0.09)) + # TX
  draw_line(x=c(0.28, 0.51), y=c(0.43, 0.205)) + # KS
  draw_line(x=c(0.345, 0.5125), y=c(0.43, 0.23)) + # NE
  draw_line(x=c(0.41, 0.76), y=c(0.43, 0.1)) + # FL
  draw_line(x=c(0.475, 0.7175), y=c(0.43, 0.16)) + # GA
  draw_line(x=c(0.54, 0.7175), y=c(0.43, 0.18)) + # TN
  draw_line(x=c(0.61, 0.8), y=c(0.43, 0.19)) + # NC
  draw_line(x=c(0.67, 0.79), y=c(0.43, 0.26)) # PA   
dev.off()


##########
# Recursion simulations to try to match observed frequencies

# This code simulates a randomly mating population 
# with a given frequency of traD and YM males
# and determines the frequency of YM and IIIM chromosomes
# required to maintain traD frequency and YM male frequency


# 1 female f1 M0 F0 Y0	III /III	tra+/tra+	X/X
# 2 female f2 M0 F1 Y0	III /III	traD/tra+	X/X
# 3 female f3 M1 F1 Y0	IIIM/III	traD/tra+	X/X
# 4 female f4 M2 F1 Y0	IIIM/IIIM	traD/tra+	X/X
# 5 female f5 M0 F1 Y1	III /III	traD/tra+	X/Y
# 6 female f6 M1 F1 Y1	IIIM/III	traD/tra+	X/Y
# 7 female f7 M2 F1 Y1	IIIM/IIIM	traD/tra+	X/Y
# 8 female f8 M0 F1 Y2	III /III	traD/tra+	Y/Y
# 9 female f9 M1 F1 Y2	IIIM/III	traD/tra+	Y/Y
#10 male   m1 M1 F0 Y0	IIIM/III	tra+/tra+	X/X
#11 male   m2 M2 F0 Y0	IIIM/IIIM	tra+/tra+	X/X
#12 male   m3 M0 F0 Y1	III /III	tra+/tra+	Y/X
#13 male   m4 M1 F0 Y1	IIIM/III	tra+/tra+	Y/X
#14 male   m5 M2 F0 Y1	IIIM/IIIM	tra+/tra+	Y/X
#15 male   m6 M0 F0 Y2	III /III	tra+/tra+	Y/Y
#16 male   m7 M1 F0 Y2	IIIM/III	tra+/tra+	Y/Y
#17 male   m8 M2 F0 Y2	IIIM/IIIM	tra+/tra+	Y/Y
#18 female f10M2 F1 Y2	IIIM/IIIM	traD/tra+	Y/Y


###############

recursions <- function(counts){ 

	f1 <- counts[1]
	f2 <- counts[2]
	f3 <- counts[3]
	f4 <- counts[4]
	f5 <- counts[5]
	f6 <- counts[6]
	f7 <- counts[7]
	f8 <- counts[8]
	f9 <- counts[9]

	m1 <- counts[10]
	m2 <- counts[11]
	m3 <- counts[12]
	m4 <- counts[13]
	m5 <- counts[14]
	m6 <- counts[15]
	m7 <- counts[16]
	m8 <- counts[17]
	f10 <- counts[18]

# new
	unlist(c(
		f1*m1*.5 + f1*m3*.5 + f1*m4*.25 + f2*m1*.25 + f2*m3*.25 + f2*m4*.125 + f3*m1*.125 + f3*m3*.125 + f3*m4*.0625 + f5*m1*.125 + f5*m3*.125 + f5*m4*.0625 + f6*m1*.0625 + f6*m3*.0625 + f6*m4*.03125, 
		f2*m1*.25 + f2*m3*.25 + f2*m4*.125 + f3*m1*.125 + f3*m3*.125 + f3*m4*.0625 + f5*m1*.125 + f5*m3*.125 + f5*m4*.0625 + f6*m1*.0625 + f6*m3*.0625 + f6*m4*.03125, 
		f2*m1*.25 + f2*m2*.5 + f2*m4*.125 + f2*m5*.25 + f3*m1*.25 + f3*m2*.25 + f3*m3*.125 + f3*m4*.125 + f3*m5*.125 + f4*m1*.25 + f4*m3*.25 + f4*m4*.125 + f5*m1*.125 + f5*m2*.25 + f5*m4*.0625 + f5*m5*.125 + f6*m1*.125 + f6*m2*.125 + f6*m3*.0625 + f6*m4*.0625 + f6*m5*.0625 + f7*m1*.125 + f7*m3*.125 + f7*m4*.0625, 
		f3*m1*.125 + f3*m2*.25 + f3*m4*.0625 + f3*m5*.125 + f4*m1*.25 + f4*m2*.5 + f4*m4*.125 + f4*m5*.25 + f6*m1*.0625 + f6*m2*.125 + f6*m4*.03125 + f6*m5*.0625 + f7*m1*.125 + f7*m2*.25 + f7*m4*.0625 + f7*m5*.125, 
		f2*m3*.25 + f2*m4*.125 + f2*m6*.5 + f2*m7*.25 + f3*m3*.125 + f3*m4*.0625 + f3*m6*.25 + f3*m7*.125 + f5*m1*.125 + f5*m3*.25 + f5*m4*.125 + f5*m6*.25 + f5*m7*.125 + f6*m1*.0625 + f6*m3*.125 + f6*m4*.0625 + f6*m6*.125 + f6*m7*.0625 + f8*m1*.25 + f8*m3*.25 + f8*m4*.125 + f9*m1*.125 + f9*m3*.125 + f9*m4*.0625, 
		f10*m1*.25 + f10*m3*.25 + f10*m4*.125 + f2*m4*.125 + f2*m5*.25 + f2*m7*.25 + f2*m8*.5 + f3*m3*.125 + f3*m4*.125 + f3*m5*.125 + f3*m6*.25 + f3*m7*.25 + f3*m8*.25 + f4*m3*.25 + f4*m4*.125 + f4*m6*.5 + f4*m7*.25 + f5*m1*.125 + f5*m2*.25 + f5*m4*.125 + f5*m5*.25 + f5*m7*.125 + f5*m8*.25 + f6*m1*.125 + f6*m2*.125 + f6*m3*.125 + f6*m4*.125 + f6*m5*.125 + f6*m6*.125 + f6*m7*.125 + f6*m8*.125 + f7*m1*.125 + f7*m3*.25 + f7*m4*.125 + f7*m6*.25 + f7*m7*.125 + f8*m1*.25 + f8*m2*.5 + f8*m4*.125 + f8*m5*.25 + f9*m1*.25 + f9*m2*.25 + f9*m3*.125 + f9*m4*.125 + f9*m5*.125, 
		f10*m1*.25 + f10*m2*.5 + f10*m4*.125 + f10*m5*.25 + f3*m4*.0625 + f3*m5*.125 + f3*m7*.125 + f3*m8*.25 + f4*m4*.125 + f4*m5*.25 + f4*m7*.25 + f4*m8*.5 + f6*m1*.0625 + f6*m2*.125 + f6*m4*.0625 + f6*m5*.125 + f6*m7*.0625 + f6*m8*.125 + f7*m1*.125 + f7*m2*.25 + f7*m4*.125 + f7*m5*.25 + f7*m7*.125 + f7*m8*.25 + f9*m1*.125 + f9*m2*.25 + f9*m4*.0625 + f9*m5*.125, 
		f5*m3*.125 + f5*m4*.0625 + f5*m6*.25 + f5*m7*.125 + f6*m3*.0625 + f6*m4*.03125 + f6*m6*.125 + f6*m7*.0625 + f8*m3*.25 + f8*m4*.125 + f8*m6*.5 + f8*m7*.25 + f9*m3*.125 + f9*m4*.0625 + f9*m6*.25 + f9*m7*.125, 
		f10*m3*.25 + f10*m4*.125 + f10*m6*.5 + f10*m7*.25 + f5*m4*.0625 + f5*m5*.125 + f5*m7*.125 + f5*m8*.25 + f6*m3*.0625 + f6*m4*.0625 + f6*m5*.0625 + f6*m6*.125 + f6*m7*.125 + f6*m8*.125 + f7*m3*.125 + f7*m4*.0625 + f7*m6*.25 + f7*m7*.125 + f8*m4*.125 + f8*m5*.25 + f8*m7*.25 + f8*m8*.5 + f9*m3*.125 + f9*m4*.125 + f9*m5*.125 + f9*m6*.25 + f9*m7*.25 + f9*m8*.25, 

		f1*m1*.5 + f1*m2*1 + f1*m4*.25 + f1*m5*.5 + f2*m1*.25 + f2*m2*.5 + f2*m4*.125 + f2*m5*.25 + f3*m1*.25 + f3*m2*.25 + f3*m3*.125 + f3*m4*.125 + f3*m5*.125 + f4*m1*.25 + f4*m3*.25 + f4*m4*.125 + f5*m1*.125 + f5*m2*.25 + f5*m4*.0625 + f5*m5*.125 + f6*m1*.125 + f6*m2*.125 + f6*m3*.0625 + f6*m4*.0625 + f6*m5*.0625 + f7*m1*.125 + f7*m3*.125 + f7*m4*.0625, 
		f3*m1*.125 + f3*m2*.25 + f3*m4*.0625 + f3*m5*.125 + f4*m1*.25 + f4*m2*.5 + f4*m4*.125 + f4*m5*.25 + f6*m1*.0625 + f6*m2*.125 + f6*m4*.03125 + f6*m5*.0625 + f7*m1*.125 + f7*m2*.25 + f7*m4*.0625 + f7*m5*.125, 
		f1*m3*.5 + f1*m4*.25 + f1*m6*1 + f1*m7*.5 + f2*m3*.25 + f2*m4*.125 + f2*m6*.5 + f2*m7*.25 + f3*m3*.125 + f3*m4*.0625 + f3*m6*.25 + f3*m7*.125 + f5*m1*.125 + f5*m3*.25 + f5*m4*.125 + f5*m6*.25 + f5*m7*.125 + f6*m1*.0625 + f6*m3*.125 + f6*m4*.0625 + f6*m6*.125 + f6*m7*.0625 + f8*m1*.25 + f8*m3*.25 + f8*m4*.125 + f9*m1*.125 + f9*m3*.125 + f9*m4*.0625, 
		f1*m4*.25 + f1*m5*.5 + f1*m7*.5 + f1*m8*1 + f10*m1*.25 + f10*m3*.25 + f10*m4*.125 + f2*m4*.125 + f2*m5*.25 + f2*m7*.25 + f2*m8*.5 + f3*m3*.125 + f3*m4*.125 + f3*m5*.125 + f3*m6*.25 + f3*m7*.25 + f3*m8*.25 + f4*m3*.25 + f4*m4*.125 + f4*m6*.5 + f4*m7*.25 + f5*m1*.125 + f5*m2*.25 + f5*m4*.125 + f5*m5*.25 + f5*m7*.125 + f5*m8*.25 + f6*m1*.125 + f6*m2*.125 + f6*m3*.125 + f6*m4*.125 + f6*m5*.125 + f6*m6*.125 + f6*m7*.125 + f6*m8*.125 + f7*m1*.125 + f7*m3*.25 + f7*m4*.125 + f7*m6*.25 + f7*m7*.125 + f8*m1*.25 + f8*m2*.5 + f8*m4*.125 + f8*m5*.25 + f9*m1*.25 + f9*m2*.25 + f9*m3*.125 + f9*m4*.125 + f9*m5*.125, 
		f10*m1*.25 + f10*m2*.5 + f10*m4*.125 + f10*m5*.25 + f3*m4*.0625 + f3*m5*.125 + f3*m7*.125 + f3*m8*.25 + f4*m4*.125 + f4*m5*.25 + f4*m7*.25 + f4*m8*.5 + f6*m1*.0625 + f6*m2*.125 + f6*m4*.0625 + f6*m5*.125 + f6*m7*.0625 + f6*m8*.125 + f7*m1*.125 + f7*m2*.25 + f7*m4*.125 + f7*m5*.25 + f7*m7*.125 + f7*m8*.25 + f9*m1*.125 + f9*m2*.25 + f9*m4*.0625 + f9*m5*.125, 
		f5*m3*.125 + f5*m4*.0625 + f5*m6*.25 + f5*m7*.125 + f6*m3*.0625 + f6*m4*.03125 + f6*m6*.125 + f6*m7*.0625 + f8*m3*.25 + f8*m4*.125 + f8*m6*.5 + f8*m7*.25 + f9*m3*.125 + f9*m4*.0625 + f9*m6*.25 + f9*m7*.125, 
		f10*m3*.25 + f10*m4*.125 + f10*m6*.5 + f10*m7*.25 + f5*m4*.0625 + f5*m5*.125 + f5*m7*.125 + f5*m8*.25 + f6*m3*.0625 + f6*m4*.0625 + f6*m5*.0625 + f6*m6*.125 + f6*m7*.125 + f6*m8*.125 + f7*m3*.125 + f7*m4*.0625 + f7*m6*.25 + f7*m7*.125 + f8*m4*.125 + f8*m5*.25 + f8*m7*.25 + f8*m8*.5 + f9*m3*.125 + f9*m4*.125 + f9*m5*.125 + f9*m6*.25 + f9*m7*.25 + f9*m8*.25, 
		f10*m4*.125 + f10*m5*.25 + f10*m7*.25 + f10*m8*.5 + f6*m4*.03125 + f6*m5*.0625 + f6*m7*.0625 + f6*m8*.125 + f7*m4*.0625 + f7*m5*.125 + f7*m7*.125 + f7*m8*.25 + f9*m4*.0625 + f9*m5*.125 + f9*m7*.125 + f9*m8*.25, 
		f10*m4*.125 + f10*m5*.25 + f10*m7*.25 + f10*m8*.5 + f6*m4*.03125 + f6*m5*.0625 + f6*m7*.0625 + f6*m8*.125 + f7*m4*.0625 + f7*m5*.125 + f7*m7*.125 + f7*m8*.25 + f9*m4*.0625 + f9*m5*.125 + f9*m7*.125 + f9*m8*.25
	))


# old
#	unlist(c(
#		f1*m1*.5 + f1*m3*.5 + f1*m4*.25 + f2*m1*.25 + f2*m3*.25 + f2*m4*.125 + f3*m1*.25 + f3*m3*.125 + f3*m4*.0625  + f5*m1*.125 + f5*m3*.125 + f5*m4*.0625 + f6*m1*.0625 + f6*m3*.0625 + f6*m4*.03125,
#		f2*m1*.25 + f2*m3*.25 + f2*m4*.125 + f3*m1*.125 + f3*m3*.125 + f3*m4*.0625 + f5*m1*.125 + f5*m3*.125 + f5*m4*.0625 + f6*m1*.0625 + f6*m3*.0625 + f6*m4*.03125,
#		f2*m1*.25 + f2*m2*.5 + f2*m4*.125 + f2*m5*.25 + f3*m1*.25 + f3*m2*.25 + f3*m3*.125 + f3*m4*.125 + f3*m5*.125 + f4*m1*.25 + f4*m3*.25 + f4*m4*.125 + f5*m1*.125 + f5*m2*.25 + f5*m4*.0625 + f5*m5*.125 + f6*m1*.125 + f6*m2*.125 + f6*m3*.0625 + f6*m4*.0625 + f6*m5*.0625 + f7*m1*.125 + f7*m3*.125 + f7*m4*.0625,
#		f3*m1*.125 + f3*m2*.25 + f3*m4*.0625 + f3*m5*.125 + f4*m1*.25 + f4*m2*.5 + f4*m4*.125 + f4*m5*.25 + f6*m1*.0625 + f6*m2*.125 + f6*m4*.03125 + f6*m5*.0625 + f7*m1*.125 + f7*m2*.25 + f7*m4*.0625 + f7*m5*.125,
#		f2*m3*.25 + f2*m4*.125 + f2*m6*.5 + f2*m7*.25 + f3*m3*.125 + f3*m4*.0625 + f3*m6*.25 + f3*m7*.125 + f5*m1*.125 + f5*m3*.25 + f5*m4*.125 + f5*m6*.25 + f5*m7*.125 + f6*m1*.0625 + f6*m3*.125 + f6*m4*.0625 + f6*m6*.125 + f6*m7*.0625 + f8*m1*.25 + f8*m3*.25 + f8*m4*.125 + f9*m1*.125 + f9*m3*.125 + f9*m4*.0625,
#		f2*m4*.125 + f2*m5*.25 + f2*m7*.25 + f2*m8*.5 + f3*m3*.125 + f3*m4*.125 + f3*m5*.125 + f3*m6*.25 + f3*m7*.25 + f3*m8*.25 + f4*m3*.25 + f4*m4*.125 + f4*m6*.5 + f4*m7*.25 + f5*m1*.125 + f5*m2*.25 + f5*m4*.125 + f5*m5*.25 + f5*m7*.125 + f5*m8*.25 + f6*m1*.125 + f6*m2*.125 + f6*m3*.125 + f6*m4*.125 + f6*m5*.125 + f6*m6*.125 + f6*m7*.125 + f6*m8*.125 + f7*m1*.125 + f7*m3*.25 + f7*m4*.125 + f7*m6*.25 + f7*m7*.125 + f8*m1*.25 + f8*m2*.5 + f8*m4*.125 + f8*m5*.25 + f9*m1*.25 + f9*m2*.25 + f9*m3*.125 + f9*m4*.125 + f9*m5*.125 + f10*m1*.25 + f10*m3*.25 + f10*m4*.125,
#		f3*m4*.0625 + f3*m5*.125 + f3*m7*.125 + f3*m8*.25 + f4*m4*.125 + f4*m5*.25 + f4*m7*.25 + f4*m8*.5 + f6*m1*.0625 + f6*m2*.125 + f6*m4*.0625 + f6*m5*.125 + f6*m7*.0625 + f6*m8*.125 + f7*m1*.125 + f7*m2*.25 + f7*m4*.125 + f7*m5*.25 + f7*m7*.125 + f7*m8*.25 + f9*m1*.125 + f9*m2*.25 + f9*m4*.0625 + f9*m5*.125 + f10*m1*.25 + f10*m2*.5 + f10*m4*.125 + f10*m5*.25,
#		f5*m3*.125 + f5*m4*.0625 + f5*m6*.25 + f5*m7*.125 + f6*m3*.0625 + f6*m4*.03125 + f6*m6*.125 + f6*m7*.0625 + f8*m3*.25 + f8*m4*.125 + f8*m6*.5 + f8*m7*.25 + f9*m3*.125 + f9*m4*.0625 + f9*m6*.25 + f9*m7*.125,
#		f5*m4*.0625 + f5*m5*.125 + f5*m7*.125 + f5*m8*.25 + f6*m3*.0625 + f6*m4*.0625 + f6*m5*.0625 + f6*m6*.125 + f6*m7*.125 + f6*m8*.125 + f7*m3*.125 + f7*m4*.0625 + f7*m6*.25 + f7*m7*.125 + f8*m4*.125 + f8*m5*.25 + f8*m7*.25 + f8*m8*.5 + f9*m3*.125 + f9*m4*.125 + f9*m5*.125 + f9*m6*.25 + f9*m7*.25 + f9*m8*.25 + f10*m3*.25 + f10*m4*.125 + f10*m6*.5 + f10*m7*.25,
#
#		f1*m1*.5 + f1*m2*1 + f1*m4*.25 + f1*m5*.5 + f2*m1*.25 + f2*m2*.5 + f2*m4*.125 + f2*m5*.25 + f3*m1*.125 + f3*m2*.25 + f3*m3*.125 + f3*m4*.125 + f3*m5*.125 + f4*m1*.25 + f4*m3*.25 + f4*m4*.125 + f5*m1*.125 + f5*m2*.25 + f5*m4*.0625 + f5*m5*.125 + f6*m1*.125 + f6*m2*.125 + f6*m3*.0625 + f6*m4*.0625 + f6*m5*.0625 + f7*m1*.125 + f7*m3*.125 + f7*m4*.0625,
#		f3*m1*.125 + f3*m2*.25 + f3*m4*.0625 + f3*m5*.125 + f4*m1*.25 + f4*m2*.5 + f4*m4*.125 + f4*m5*.25 + f6*m1*.0625 + f6*m2*.125 + f6*m4*.03125 + f6*m5*.0625 + f7*m1*.125 + f7*m2*.25 + f7*m4*.0625 + f7*m5*.125,
#		f1*m3*.5 + f1*m4*.25 + f1*m6*1 + f1*m7*.5 + f2*m3*.25 + f2*m4*.125 + f2*m6*.5 + f2*m7*.25 + f3*m3*.125 + f3*m4*.0625 + f3*m6*.25 + f3*m7*.125 + f5*m1*.125 + f5*m3*.25 + f5*m4*.125 + f5*m6*.25 + f5*m7*.125 + f6*m1*.0625 + f6*m3*.125 + f6*m4*.0625 + f6*m6*.125 + f6*m7*.0625 + f8*m1*.25 + f8*m3*.25 + f8*m4*.125 + f9*m1*.125 + f9*m3*.125 + f9*m4*.0625,
#		f1*m4*.25 + f1*m5*.5 + f1*m7*.5 + f1*m8*1 + f2*m4*.125 + f2*m5*.25 + f2*m7*.25 + f2*m8*.5 + f3*m3*.125 + f3*m4*.125 + f3*m5*.125 + f3*m6*.25 + f3*m7*.25 + f3*m8*.25 + f4*m3*.25 + f4*m4*.125 + f4*m6*.5 + f4*m7*.25 + f5*m1*.125 + f5*m2*.25 + f5*m4*.125 + f5*m5*.25 + f5*m7*.125 + f5*m8*.25 + f6*m1*.125 + f6*m2*.125 + f6*m3*.125 + f6*m4*.125 + f6*m5*.125 + f6*m6*.125 + f6*m7*.125 + f6*m8*.125 + f7*m1*.125 + f7*m3*.25 + f7*m4*.125 + f7*m6*.25 + f7*m7*.125 + f8*m1*.25 + f8*m2*.5 + f8*m4*.125 + f8*m5*.25 + f9*m1*.25 + f9*m2*.25 + f9*m3*.125 + f9*m4*.125 + f9*m5*.125 + f10*m1*.25 + f10*m3*.25 + f10*m4*.125,
#		f3*m4*.0625 + f3*m5*.125 + f3*m7*.125 + f3*m8*.25 + f4*m4*.125 + f4*m5*.25 + f4*m7*.25 + f4*m8*.5 + f6*m1*.0625 + f6*m2*.125 + f6*m4*.0625 + f6*m5*.125 + f6*m7*.0625 + f6*m8*.125 + f7*m1*.125 + f7*m2*.25 + f7*m4*.125 + f7*m5*.25 + f7*m7*.125 + f7*m8*.25 + f9*m1*.125 + f9*m2*.25 + f9*m4*.0625 + f9*m5*.125 + f10*m1*.25 + f10*m2*.5 + f10*m4*.125 + f10*m5*.25,
#		f5*m3*.125 + f5*m4*.0625 + f5*m6*.25 + f5*m7*.125 + f6*m3*.0625 + f6*m4*.03125 + f6*m6*.125 + f6*m7*.0625 + f8*m3*.25 + f8*m4*.125 + f8*m6*.5 + f8*m7*.25 + f9*m3*.125 + f9*m4*.0625 + f9*m6*.25 + f9*m7*.125,
#		f5*m4*.0625 + f5*m5*.125 + f5*m7*.125 + f5*m8*.25 + f6*m3*.0625 + f6*m4*.0625 + f6*m5*.0625 + f6*m6*.125 + f6*m7*.125 + f6*m8*.125 + f7*m3*.125 + f7*m4*.0625 + f7*m6*.25 + f7*m7*.125 + f8*m4*.125 + f8*m5*.25 + f8*m7*.25 + f8*m8*.5 + f9*m3*.125 + f9*m4*.125 + f9*m5*.125 + f9*m6*.25 + f9*m7*.25 + f9*m8*.25 + f10*m3*.25 + f10*m4*.125 + f10*m6*.5 + f10*m7*.25,
#		f6*m4*.03125 + f6*m5*.0625 + f6*m7*.0625 + f6*m8*.125 + f7*m4*.0625 + f7*m5*.125 + f7*m7*.125 + f7*m8*.25 + f9*m4*.0625 + f9*m5*.125 + f9*m7*.125 + f9*m8*.25 + f10*m4*.125 + f10*m5*.25 + f10*m7*.25 + f10*m8*.5,
#		f6*m4*.03125 + f6*m5*.0625 + f6*m7*.0625 + f6*m8*.125 + f7*m4*.0625 + f7*m5*.125 + f7*m7*.125 + f7*m8*.25 + f9*m4*.0625 + f9*m5*.125 + f9*m7*.125 + f9*m8*.25 + f10*m4*.125 + f10*m5*.25 + f10*m7*.25 + f10*m8*.5
#	))
}

genotype.freqs <- function(freqs){ # send function total genotype frequencies from recursions and it will calculate frequencies that sum to 1
	freqs/sum(freqs)
}

allele.freqs <- function(genotype_freqs){ # calculates allele frequencies from genotype frequencies
	X <- (sum(genotype_freqs[c(1,2,3,4,10,11)]) + 0.5*sum(genotype_freqs[c(5,6,7,12,13,14)])) / (sum(genotype_freqs))
	Y <- (sum(genotype_freqs[c(8,9,15,16,17,18)]) + 0.5*sum(genotype_freqs[c(5,6,7,12,13,14)])) / (sum(genotype_freqs))
	III  <- (sum(genotype_freqs[c(1,2,5,8,12,15)]) + 0.5*sum(genotype_freqs[c(3,6,9,10,13,16)])) / (sum(genotype_freqs))
	IIIM <- (sum(genotype_freqs[c(4,7,11,14,17,18)]) + 0.5*sum(genotype_freqs[c(3,6,9,10,13,16)])) / (sum(genotype_freqs))
	tra  <- (sum(genotype_freqs[c(1,10:17)]) + 0.5*sum(genotype_freqs[c(2:9,18)])) / (sum(genotype_freqs))
	traD <- (0.5*sum(genotype_freqs[c(2:9,18)])) / (sum(genotype_freqs))
	
	c(X, Y, III, IIIM, tra, traD)
}

#10 male   m1 M1 F0 Y0	IIIM/III	tra+/tra+	X/X
#11 male   m2 M2 F0 Y0	IIIM/IIIM	tra+/tra+	X/X
#12 male   m3 M0 F0 Y1	III /III	tra+/tra+	Y/X
#13 male   m4 M1 F0 Y1	IIIM/III	tra+/tra+	Y/X
#14 male   m5 M2 F0 Y1	IIIM/IIIM	tra+/tra+	Y/X
#15 male   m6 M0 F0 Y2	III /III	tra+/tra+	Y/Y
#16 male   m7 M1 F0 Y2	IIIM/III	tra+/tra+	Y/Y
#17 male   m8 M2 F0 Y2	IIIM/IIIM	tra+/tra+	Y/Y

allele.freqs.sex <- function(genotype_freqs){ # calculates allele frequencies in each sex 
	Y_male <- sum(genotype_freqs[c(12,13,14,15,16,17)]) / sum(genotype_freqs[10:17]) # frequency of males that have YM
	IIIM_male <- sum(genotype_freqs[c(10,11,13,14,16,17)]) / sum(genotype_freqs[10:17]) # frequency of males that have IIIM
	traD_female <- sum(genotype_freqs[c(2:9,18)]) / sum(genotype_freqs[c(1:9,18)]) # frequency of females that have traD
	c(Y_male, IIIM_male, traD_female)
}

sex.ratio <- function(genotype_freqs){ # calculates the frequency of males and females based on the genotype frequencies
	female <- sum(genotype_freqs[c(1:9,18)])/sum(genotype_freqs)
	male   <- sum(genotype_freqs[c(10:17)])/sum(genotype_freqs)
	
	c(female, male)
}
sex.chr <- function(genotype_freqs){
	X_fem <- (sum(genotype_freqs[c(1,2,3,4)]) + 0.5*sum(genotype_freqs[c(5,6,7)])) / (sum(genotype_freqs[c(1:4,10:11)]) + 0.5*sum(genotype_freqs[c(5:7,12:14)]))
	X_mal <- (sum(genotype_freqs[c(10,11)]) + 0.5*sum(genotype_freqs[c(12,13,14)])) / (sum(genotype_freqs[c(1:4,10:11)]) + 0.5*sum(genotype_freqs[c(5:7,12:14)]))
	Y_fem <- (sum(genotype_freqs[c(8,9,18)]) + 0.5*sum(genotype_freqs[c(5,6,7)])) / (sum(genotype_freqs[c(8:9,15:18)]) + 0.5*sum(genotype_freqs[c(5:7,12:14)]))
	Y_mal <- (sum(genotype_freqs[c(15,16,17)]) + 0.5*sum(genotype_freqs[c(12,13,14)])) / (sum(genotype_freqs[c(8:9,15:18)]) + 0.5*sum(genotype_freqs[c(5:7,12:14)]))
	c(X_fem, X_mal, Y_fem, Y_mal)
}
chrIII <- function(genotype_freqs){
	III_fem <- (sum(genotype_freqs[c(1,2,5,8)]) + 0.5*sum(genotype_freqs[c(3,6,9)])) / (sum(genotype_freqs[c(1,2,5,8,12,15)]) + 0.5*sum(genotype_freqs[c(3,6,9,10,13,16)]))
	III_mal <- (sum(genotype_freqs[c(12,15)]) + 0.5*sum(genotype_freqs[c(10,13,16)])) / (sum(genotype_freqs[c(1,2,5,8,12,15)]) + 0.5*sum(genotype_freqs[c(3,6,9,10,13,16)]))
	IIIM_fem <- (sum(genotype_freqs[c(4,7,18)]) + 0.5*sum(genotype_freqs[c(3,6,9)])) / (sum(genotype_freqs[c(4,7,18,11,14,17)]) + 0.5*sum(genotype_freqs[c(3,6,9,10,13,16)]))
	IIIM_mal <- (sum(genotype_freqs[c(11,14,17)]) + 0.5*sum(genotype_freqs[c(10,13,16)])) / (sum(genotype_freqs[c(4,7,18,11,14,17)]) + 0.5*sum(genotype_freqs[c(3,6,9,10,13,16)]))
	c(III_fem, III_mal, IIIM_fem, IIIM_mal)
}
tra <- function(genotype_freqs){
	tra_fem <- (sum(genotype_freqs[c(1)]) + 0.5*sum(genotype_freqs[c(2:9,18)])) / (sum(genotype_freqs[c(1,10:17)]) + 0.5*sum(genotype_freqs[c(2:9,18)]))
	tra_mal <- (sum(genotype_freqs[c(10:17)])) / (sum(genotype_freqs[c(1,10:17)]) + 0.5*sum(genotype_freqs[c(2:9,18)]))
	c(tra_fem, tra_mal)
}


# Run recursions with random mating
simulator <- function(initial_freqs, generations){
	return.df <- data.frame(
		generation = c(0:generations),
		f1 = NA, f2 = NA, f3 = NA, f4 = NA, f5 = NA, f6 = NA, f7 = NA, f8 = NA, f9 = NA,
		m1 = NA, m2 = NA, m3 = NA, m4 = NA, m5 = NA, m6 = NA, m7 = NA, m8 = NA, f10 = NA,
		X = NA, Y = NA, III = NA, IIIM = NA, tra = NA, traD = NA,
		female = NA, male = NA, 
		X_fem = NA, X_mal = NA, Y_fem = NA, Y_mal = NA, III_fem = NA, III_mal = NA, IIIM_fem = NA, IIIM_mal = NA, tra_fem = NA, tra_mal = NA
	)
	return.df[1,2:19] <- initial_freqs
	
	# run recursions
	for(i in 1:generations){
		return.df[i+1,2:19] <- genotype.freqs(as.numeric(recursions(return.df[i, 2:19])))
	}
	
	# calculate allele frequencies
	return.df[,20:25] <- t(apply(return.df[,2:19], 1, allele.freqs))
	
	# calculate sex ratio
	return.df[,26:27] <- t(apply(return.df[,2:19], 1, sex.ratio))
	
	# calculate frequency of X and Y chromosomes found in females and males
	return.df[,28:31] <- t(apply(return.df[,2:19], 1, sex.chr))
		
	# calculate frequency of III and IIIM chromosomes found in females and males
	return.df[,32:35] <- t(apply(return.df[,2:19], 1, chrIII))
	
	# calculate frequency of tra+ chromosomes found in females and males
	return.df[,36:37] <- t(apply(return.df[,2:19], 1, tra))
	
	return.df
}

# calculate genotype frequencies
geno.freqs <- function(allele_freq){
	geno_freqs <- data.frame(	
		Pop = allele_freq$Pop, 
		M0 = (allele_freq$III)^2, 
		M1 = 2*(allele_freq$III)*(allele_freq$IIIM),
		M2 = (allele_freq$IIIM)^2, 
		F0 = (allele_freq$tra)^2,
		F1 = (allele_freq$tra)*(allele_freq$traD), # Multiply tra*traD freq (not by 2) because traD can only be inherited from mother
		Y0 = (allele_freq$X)^2, 
		Y1 = 2*(allele_freq$X)*(allele_freq$YM),
		Y2 = (allele_freq$YM)^2
	)
	geno_freqs$f1 = geno_freqs$M0 * geno_freqs$F0 * geno_freqs$Y0
	geno_freqs$f2 = geno_freqs$M0 * geno_freqs$F1 * geno_freqs$Y0
	geno_freqs$f3 = geno_freqs$M1 * geno_freqs$F1 * geno_freqs$Y0
	geno_freqs$f4 = geno_freqs$M2 * geno_freqs$F1 * geno_freqs$Y0
	geno_freqs$f5 = geno_freqs$M0 * geno_freqs$F1 * geno_freqs$Y1
	geno_freqs$f6 = geno_freqs$M1 * geno_freqs$F1 * geno_freqs$Y1
	geno_freqs$f7 = geno_freqs$M2 * geno_freqs$F1 * geno_freqs$Y1
	geno_freqs$f8 = geno_freqs$M0 * geno_freqs$F1 * geno_freqs$Y2
	geno_freqs$f9 = geno_freqs$M1 * geno_freqs$F1 * geno_freqs$Y2
	geno_freqs$m1 = geno_freqs$M1 * geno_freqs$F0 * geno_freqs$Y0
	geno_freqs$m2 = geno_freqs$M2 * geno_freqs$F0 * geno_freqs$Y0
	geno_freqs$m3 = geno_freqs$M0 * geno_freqs$F0 * geno_freqs$Y1
	geno_freqs$m4 = geno_freqs$M1 * geno_freqs$F0 * geno_freqs$Y1
	geno_freqs$m5 = geno_freqs$M2 * geno_freqs$F0 * geno_freqs$Y1
	geno_freqs$m6 = geno_freqs$M0 * geno_freqs$F0 * geno_freqs$Y2
	geno_freqs$m7 = geno_freqs$M1 * geno_freqs$F0 * geno_freqs$Y2
	geno_freqs$m8 = geno_freqs$M2 * geno_freqs$F0 * geno_freqs$Y2
	geno_freqs$f10= geno_freqs$M2 * geno_freqs$F1 * geno_freqs$Y2
	geno_freqs
}
geno.freqs.norm <- function(geno_freq){
	geno_freqs_norm <- cbind(
		Pop = geno_freq$Pop,
		as.data.frame(t(apply(geno_freq[,10:27], 1, genotype.freqs))),
		X=NA, Y=NA, III=NA, IIIM=NA, tra=NA, traD=NA
	)
	geno_freqs_norm[,20:25] <- t(apply(geno_freqs_norm[,2:19], 1, allele.freqs))
	geno_freqs_norm
}


# Plot over time:
#	1. Frequency of alleles
#	2. Sex-ratio

# Include observed frequencies in plot
gen.plot.obs <- function(data.df, obs_Y, obs_traD, filename){
	data.df$YM_male <- NA
	data.df$IIIM_male <- NA
	data.df$traD_female <- NA
	data.df[,38:40] <- t(apply(data.df[,2:19], 1, allele.freqs.sex))
	pdf(filename, width=10, height=5)
#		plot(data.df$generation, data.df$X, ylim=c(0,1), type="b", pch="X", cex=0.5, col="red", xlab="generation", ylab="frequency", main=filename)
#		points(data.df$generation, data.df$Y, type="b", pch="Y", col="blue", cex=0.5)
#		plot(data.df$generation, data.df$Y, ylim=c(0,1), type="b", pch="Y", col="blue", cex=0.5, xlab="generation", ylab="frequency", main=filename, yaxt='n')
		plot(data.df$generation, data.df$YM_male, ylim=c(0,1), type="b", pch="Y", col="blue", cex=0.5, xlab="generation", ylab="frequency", main=filename, yaxt='n')
		axis(side=2, at=c(0,0.25, 0.5, 0.75, 1), las=1)
#		points(data.df$generation, data.df$III, type="b", pch=1, col="darkgreen", cex=0.5)
		points(data.df$generation, data.df$IIIM_male, type="b", pch="M", col="orange", cex=0.5)
#		points(data.df$generation, data.df$tra, type="b", pch="+", col="black", cex=0.5)
		points(data.df$generation, data.df$traD_female, type="b", pch="D", col="magenta", cex=0.5)
#		points(data.df$generation, data.df$female, type="b", pch="f", cex=0.5)
		points(data.df$generation, data.df$male, type="b", pch="m", cex=0.5)
		abline(h=c(0.5, 0.25, 0.75), lty=1, col="gray")			
		abline(h=obs_Y, lty=2, col=c("blue"))
		abline(h=obs_traD, lty=2, col=c("magenta"))
	dev.off()	
}


# YM males can be: 
#12 male   m3 M0 F0 Y1	III /III	tra+/tra+	Y/X
#13 male   m4 M1 F0 Y1	IIIM/III	tra+/tra+	Y/X
#14 male   m5 M2 F0 Y1	IIIM/IIIM	tra+/tra+	Y/X
#15 male   m6 M0 F0 Y2	III /III	tra+/tra+	Y/Y
#16 male   m7 M1 F0 Y2	IIIM/III	tra+/tra+	Y/Y
#17 male   m8 M2 F0 Y2	IIIM/IIIM	tra+/tra+	Y/Y

# traD females can be: 
# 2 female f2 M0 F1 Y0	III /III	traD/tra+	X/X
# 3 female f3 M1 F1 Y0	IIIM/III	traD/tra+	X/X
# 4 female f4 M2 F1 Y0	IIIM/IIIM	traD/tra+	X/X
# 5 female f5 M0 F1 Y1	III /III	traD/tra+	X/Y
# 6 female f6 M1 F1 Y1	IIIM/III	traD/tra+	X/Y
# 7 female f7 M2 F1 Y1	IIIM/IIIM	traD/tra+	X/Y
# 8 female f8 M0 F1 Y2	III /III	traD/tra+	Y/Y
# 9 female f9 M1 F1 Y2	IIIM/III	traD/tra+	Y/Y
#18 female f10M2 F1 Y2	IIIM/IIIM	traD/tra+	Y/Y


# generate frequencies of genotypes given observed frequencies of YM males and traD females

# minimum IIIM frequency is all males that are not YM: 
male_freqs$IIIM_min <- 1 - male_freqs$YMfreq


allele_freqs_v1 <- data.frame(
	Pop = male_freqs$Pop, 
	YM = male_freqs$YMfreq / 2, # YM males have 50% Y and 50% X (this is a lower bound on YM freq in males)
	X = 1 - male_freqs$YMfreq/2, # upper bound on X freq in males (see YM freq calc)
	IIIM = male_freqs$IIIM_min / 2, # IIIM males have 50% IIIM and 50% III (this is a lower bound on IIIM freq in males)
	III = 1 - male_freqs$IIIM_min/2, # upper bound on III freq in males (see IIIM freq calc)
	traD = female_freqs$traDfreq/4, # this is the frequency of traD population
	tra = 1 - female_freqs$traDfreq/4 # this is the frequency of tra in population
)

geno_freqs_v1 <- geno.freqs(allele_freqs_v1)
geno_freqs_v1_norm <- geno.freqs.norm(geno_freqs_v1)


# Compare observed and expected  
# test for within 0.002
obs.exp.df <- function(males, females, CA,FL,GA,KS,NC,NE,PA,TN,TX){
	return.df <- data.frame(
		pop = male_freqs$Pop, 
		YM_obs = male_freqs$YMfreq, 
		traD_obs = female_freqs$traDfreq, 
		YM_simulation = NA, 
		traD_simulation = NA
	)
	return.df[,4:5] <- rbind(
		allele.freqs.sex(CA[11,2:19])[c(1,3)], 
		allele.freqs.sex(FL[11,2:19])[c(1,3)],
		allele.freqs.sex(GA[11,2:19])[c(1,3)],
		allele.freqs.sex(KS[11,2:19])[c(1,3)],
		allele.freqs.sex(NC[11,2:19])[c(1,3)],
		allele.freqs.sex(NE[11,2:19])[c(1,3)],
		allele.freqs.sex(PA[11,2:19])[c(1,3)],
		allele.freqs.sex(TN[11,2:19])[c(1,3)],
		allele.freqs.sex(TX[11,2:19])[c(1,3)]
	)
	return.df$YM_diff <- return.df$YM_simulation - return.df$YM_obs 
	return.df$traD_diff <- return.df$traD_simulation - return.df$traD_obs

	return.df$YM_fit <- ifelse(abs(return.df$YM_diff) < 0.002, "fit", "*")
	return.df$traD_fit <- ifelse(abs(return.df$traD_diff) < 0.002, "fit", "*")
	
	return.df
}


# simulate 10 generations with starting frequencies
CA_v1_10 <- simulator(geno_freqs_v1_norm[1,2:19], 10)
FL_v1_10 <- simulator(geno_freqs_v1_norm[2,2:19], 10)
GA_v1_10 <- simulator(geno_freqs_v1_norm[3,2:19], 10)
KS_v1_10 <- simulator(geno_freqs_v1_norm[4,2:19], 10)
NC_v1_10 <- simulator(geno_freqs_v1_norm[5,2:19], 10)
NE_v1_10 <- simulator(geno_freqs_v1_norm[6,2:19], 10)
PA_v1_10 <- simulator(geno_freqs_v1_norm[7,2:19], 10)
TN_v1_10 <- simulator(geno_freqs_v1_norm[8,2:19], 10)
TX_v1_10 <- simulator(geno_freqs_v1_norm[9,2:19], 10)


# Compare observed and expected in v1
v1.df <- obs.exp.df(male_freqs, female_freqs, CA_v1_10, FL_v1_10, GA_v1_10, KS_v1_10, NC_v1_10, NE_v1_10, PA_v1_10, TN_v1_10, TX_v1_10)


# CA
# traD frequency is low
# YM male frequency is high
# start population with YM=0.1, traD=0.15, IIM=0.4

# FL
# good fit

# GA
# traD frequency is slightly high
# YM male frequency is slightly high
# start population with YM=0.4, traD=0.05, IIIM=0.04

# KS
# YM frequency is good fit
# traD frequency is slightly high
# start population with traD=0.01, IIIM=0.01, YM=0.475

# NC
# good fit

# NE
# traD frequency is a little high
# YM frequency is a little high
# start with YM=0.2, traD=0.025

# PA
# YM frequency is good
# traD frequency is high
# start population with YM=0.9, IIIM=0.1, traD=0.02

# TN
# traD frequency is slightly low
# YM male frequency is high
# start with YM=0.3, IIIM=0.1, traD=0.07

# TX
# traD frequency is high
# YM male frequency is good, but slightly high
# start with YM=0.45, IIIM=0.01, traD=0.03


#########
# update allele frequencies based on results from previous simulations
allele_freqs_v2 <- allele_freqs_v1

#1 CA: start population with YM=0.1, traD=0.15, IIM=0.4
allele_freqs_v2$YM[1] <- 0.1
allele_freqs_v2$X[1] <- 1 - allele_freqs_v2$YM[1]
allele_freqs_v2$traD[1] <- 0.15
allele_freqs_v2$tra[1] <- 1 - allele_freqs_v2$traD[1]
allele_freqs_v2$IIIM[1] <- 0.4
allele_freqs_v2$III[1] <- 1 - allele_freqs_v2$IIIM[1]

#2 FL: god fit

#3 GA: start population with YM=0.4, traD=0.05, IIIM=0.04
allele_freqs_v2$YM[3] <- 0.4
allele_freqs_v2$X[3] <- 1 - allele_freqs_v2$YM[3]
allele_freqs_v2$traD[3] <- 0.05
allele_freqs_v2$tra[3] <- 1 - allele_freqs_v2$traD[3]
allele_freqs_v2$IIIM[3] <- 0.04
allele_freqs_v2$III[3] <- 1 - allele_freqs_v2$IIIM[3]

#4 KS: start population with traD=0.01, IIIM=0.01, YM=0.475
allele_freqs_v2$YM[4] <- 0.474
allele_freqs_v2$X[4] <- 1 - allele_freqs_v2$YM[4]
allele_freqs_v2$IIIM[4] <- 0.01
allele_freqs_v2$III[4] <- 1 - allele_freqs_v2$IIIM[4]
allele_freqs_v2$traD[4] <- 0.01
allele_freqs_v2$tra[4] <- 1 - allele_freqs_v2$traD[4]

#5 NC: good fit

#6 NE: start with YM=0.2, traD=0.025
allele_freqs_v2$YM[6] <- 0.2
allele_freqs_v2$X[6] <- 1 - allele_freqs_v2$YM[6]
allele_freqs_v2$traD[6] <- 0.025
allele_freqs_v2$tra[6] <- 1 - allele_freqs_v2$traD[6]

#7 PA: start population with YM=0.9, IIIM=0.1, traD=0.02
allele_freqs_v2$YM[7] <- 0.9
allele_freqs_v2$X[7] <- 1 - allele_freqs_v2$YM[7]
allele_freqs_v2$IIIM[7] <- 0.1
allele_freqs_v2$III[7] <- 1 - allele_freqs_v2$IIIM[7]
allele_freqs_v2$traD[7] <- 0.02
allele_freqs_v2$tra[7] <- 1 - allele_freqs_v2$traD[7]

#8 TN: start with YM=0.3, IIIM=0.1, traD=0.07
allele_freqs_v2$YM[8] <- 0.3
allele_freqs_v2$X[8] <- 1 - allele_freqs_v2$YM[8]
allele_freqs_v2$IIIM[8] <- 0.1
allele_freqs_v2$III[8] <- 1 - allele_freqs_v2$IIIM[8]
allele_freqs_v2$traD[8] <- 0.07
allele_freqs_v2$tra[8] <- 1 - allele_freqs_v2$traD[8]

#9 TX: start with YM=0.45, IIIM=0.01, traD=0.03
allele_freqs_v2$YM[9] <- 0.45
allele_freqs_v2$X[9] <- 1 - allele_freqs_v2$YM[9]
allele_freqs_v2$IIIM[9] <- 0.01
allele_freqs_v2$III[9] <- 1 - allele_freqs_v2$IIIM[9]
allele_freqs_v2$traD[9] <- 0.03
allele_freqs_v2$tra[9] <- 1 - allele_freqs_v2$traD[9]


geno_freqs_v2 <- geno.freqs(allele_freqs_v2)
geno_freqs_v2_norm <- geno.freqs.norm(geno_freqs_v2)


# simulate 10 generations with new starting frequencies to see if they are stable
CA_v2_10 <- simulator(geno_freqs_v2_norm[1,2:19], 10)
GA_v2_10 <- simulator(geno_freqs_v2_norm[3,2:19], 10)
KS_v2_10 <- simulator(geno_freqs_v2_norm[4,2:19], 10)
NE_v2_10 <- simulator(geno_freqs_v2_norm[6,2:19], 10)
PA_v2_10 <- simulator(geno_freqs_v2_norm[7,2:19], 10)
TN_v2_10 <- simulator(geno_freqs_v2_norm[8,2:19], 10)
TX_v2_10 <- simulator(geno_freqs_v2_norm[9,2:19], 10)

v2.df <- obs.exp.df(male_freqs, female_freqs, CA_v2_10, FL_v1_10, GA_v2_10, KS_v2_10, NC_v1_10, NE_v2_10, PA_v2_10, TN_v2_10, TX_v2_10)


#1 CA: start population with YM=0.1, traD=0.15, IIM=0.4
# traD frequency good fit
# YM male frequency is too low
# start with higher YM (0.125) and lower IIIM (0.375)

# FL
# very good fit from naive

#3 GA: start population with YM=0.4, traD=0.05, IIIM=0.04
# YM too high
# traD too high
# YM=0.35, traD=0.02, IIIM=0.04

#4 KS: start population with traD=0.01, IIIM=0.01, YM=0.475
# YM frequency is slightly high
# traD frequency is high
# start population YM=0.47, traD=0.05, IIIM=0.015

# NC
# good fit from naive

#6 NE: start with YM=0.2, traD=0.025
# traD frequency is low and YM frequency is low
# go closer to v1
# start YM=0.24, traD=0.03, IIIM=0.275

#7 PA: start population with YM=0.9, IIIM=0.1, traD=0.02
# YM frequency is low and traD frequency is way too high in simulated population
# return YM to v1, and lower IIIM
# YM=0.5, traD=0.02, IIIM=0.05

#8 TN: start with YM=0.3, IIIM=0.1, traD=0.07
# traD frequency is low
# YM male frequency is low (make intmdt between v1 and v2)
# YM=0.35, traD=0.07, IIIM=0.1

#9 TX: start with YM=0.45, IIIM=0.01, traD=0.03
# traD frequency is too high in simulated population
# YM male frequency is slightly low
# YM=0.46, IIIM=0.01, traD=0.035


#########
# update allele frequencies based on results from previous simulations
allele_freqs_v3 <- allele_freqs_v2

#1 CA: higher YM (0.125) and lower IIIM (0.375)
allele_freqs_v3$YM[1] <- 0.125
allele_freqs_v3$X[1] <- 1 - allele_freqs_v3$YM[1]
allele_freqs_v3$traD[1] <- 0.15
allele_freqs_v3$tra[1] <- 1 - allele_freqs_v3$traD[1]
allele_freqs_v3$IIIM[1] <- 0.375
allele_freqs_v3$III[1] <- 1 - allele_freqs_v3$IIIM[1]

#2 FL: god fit

#3 GA: YM=0.35, traD=0.02, IIIM=0.04
allele_freqs_v3$YM[3] <- 0.35
allele_freqs_v3$X[3] <- 1 - allele_freqs_v3$YM[3]
allele_freqs_v3$traD[3] <- 0.02
allele_freqs_v3$tra[3] <- 1 - allele_freqs_v3$traD[3]
allele_freqs_v3$IIIM[3] <- 0.04
allele_freqs_v3$III[3] <- 1 - allele_freqs_v3$IIIM[3]

#4 KS: YM=0.47, traD=0.05, IIIM=0.015
allele_freqs_v3$YM[4] <- 0.47
allele_freqs_v3$X[4] <- 1 - allele_freqs_v3$YM[4]
allele_freqs_v3$IIIM[4] <- 0.015
allele_freqs_v3$III[4] <- 1 - allele_freqs_v3$IIIM[4]
allele_freqs_v3$traD[4] <- 0.05
allele_freqs_v3$tra[4] <- 1 - allele_freqs_v3$traD[4]

#5 NC: good fit

#6 NE: YM=0.24, traD=0.03, IIIM=0.275
allele_freqs_v3$YM[6] <- 0.24
allele_freqs_v3$X[6] <- 1 - allele_freqs_v3$YM[6]
allele_freqs_v3$traD[6] <- 0.03
allele_freqs_v3$tra[6] <- 1 - allele_freqs_v3$traD[6]
allele_freqs_v3$IIIM[6] <- 0.275
allele_freqs_v3$III[6] <- 1 - allele_freqs_v3$IIIM[6]

#7 PA: YM=0.5, traD=0.02, IIIM=0.05
allele_freqs_v3$YM[7] <- 0.5
allele_freqs_v3$X[7] <- 1 - allele_freqs_v3$YM[7]
allele_freqs_v3$IIIM[7] <- 0.05
allele_freqs_v3$III[7] <- 1 - allele_freqs_v3$IIIM[7]
allele_freqs_v3$traD[7] <- 0.02
allele_freqs_v3$tra[7] <- 1 - allele_freqs_v3$traD[7]

#8 TN: YM=0.35, traD=0.07, IIIM=0.1
allele_freqs_v3$YM[8] <- 0.35
allele_freqs_v3$X[8] <- 1 - allele_freqs_v3$YM[8]
allele_freqs_v3$IIIM[8] <- 0.1
allele_freqs_v3$III[8] <- 1 - allele_freqs_v3$IIIM[8]
allele_freqs_v3$traD[8] <- 0.07
allele_freqs_v3$tra[8] <- 1 - allele_freqs_v3$traD[8]

#9 TX:  YM=0.46, IIIM=0.1, traD=0.035
allele_freqs_v3$YM[9] <- 0.46
allele_freqs_v3$X[9] <- 1 - allele_freqs_v3$YM[9]
allele_freqs_v3$IIIM[9] <- 0.01
allele_freqs_v3$III[9] <- 1 - allele_freqs_v3$IIIM[9]
allele_freqs_v3$traD[9] <- 0.035
allele_freqs_v3$tra[9] <- 1 - allele_freqs_v3$traD[9]


geno_freqs_v3 <- geno.freqs(allele_freqs_v3)
geno_freqs_v3_norm <- geno.freqs.norm(geno_freqs_v3)


# simulate 10 generations 
CA_v3_10 <- simulator(geno_freqs_v3_norm[1,2:19], 10)
GA_v3_10 <- simulator(geno_freqs_v3_norm[3,2:19], 10)
KS_v3_10 <- simulator(geno_freqs_v3_norm[4,2:19], 10)
NE_v3_10 <- simulator(geno_freqs_v3_norm[6,2:19], 10)
PA_v3_10 <- simulator(geno_freqs_v3_norm[7,2:19], 10)
TN_v3_10 <- simulator(geno_freqs_v3_norm[8,2:19], 10)
TX_v3_10 <- simulator(geno_freqs_v3_norm[9,2:19], 10)

v3.df <- obs.exp.df(male_freqs, female_freqs, CA_v3_10, FL_v1_10, GA_v3_10, KS_v3_10, NC_v1_10, NE_v3_10, PA_v3_10, TN_v3_10, TX_v3_10)


#1 CA: 
# YM male frequency is too high
# traD slightly low
# YM=0.12, IIIM=0.38

# FL
# very good fit from naive

#3 GA: 
# YM slightly low
# traD low
# YM=0.36, traD=0.025, IIIM=0.05

#4 KS: 
# YM frequency is slightly low
# traD frequency is very high
# start population YM=0.471, traD=0.01, IIIM=0.01

# NC
# good fit from naive

#6 NE: 
# traD frequency is low and YM frequency is high
# lower IIIM
# start YM=0.24, traD=0.03, IIIM=0.26

#7 PA: 
# YM frequency is low and traD frequency is too high in simulated population
# YM=0.6, traD=0.02, IIIM=0.025

#8 TN: 
# traD frequency is low
# YM male frequency is slightly low
# YM=0.36, traD=0.072, IIIM=0.11

#9 TX: 
# traD frequency is slightly high in simulated population
# YM male frequency is slightly high
# YM=0.45, IIIM=0.01, traD=0.035


#########
# update allele frequencies based on results from previous simulations
allele_freqs_v4 <- allele_freqs_v3

#1 CA: YM=0.12, IIIM=0.38
allele_freqs_v4$YM[1] <- 0.12
allele_freqs_v4$X[1] <- 1 - allele_freqs_v4$YM[1]
#allele_freqs_v4$traD[1] <- 
allele_freqs_v4$tra[1] <- 1 - allele_freqs_v4$traD[1]
allele_freqs_v4$IIIM[1] <- 0.38
allele_freqs_v4$III[1] <- 1 - allele_freqs_v4$IIIM[1]

#2 FL: god fit

#3 GA: YM=0.36, traD=0.025, IIIM=0.05
allele_freqs_v4$YM[3] <- 0.36
allele_freqs_v4$X[3] <- 1 - allele_freqs_v4$YM[3]
allele_freqs_v4$traD[3] <- 0.025
allele_freqs_v4$tra[3] <- 1 - allele_freqs_v4$traD[3]
allele_freqs_v4$IIIM[3] <- 0.05
allele_freqs_v4$III[3] <- 1 - allele_freqs_v4$IIIM[3]

#4 KS: YM=0.471, traD=0.01, IIIM=0.01
allele_freqs_v4$YM[4] <- 0.471
allele_freqs_v4$X[4] <- 1 - allele_freqs_v4$YM[4]
allele_freqs_v4$IIIM[4] <- 0.01
allele_freqs_v4$III[4] <- 1 - allele_freqs_v4$IIIM[4]
allele_freqs_v4$traD[4] <- 0.01
allele_freqs_v4$tra[4] <- 1 - allele_freqs_v4$traD[4]

#5 NC: good fit

#6 NE: YM=0.24, traD=0.03, IIIM=0.26
allele_freqs_v4$YM[6] <- 0.24
allele_freqs_v4$X[6] <- 1 - allele_freqs_v4$YM[6]
allele_freqs_v4$traD[6] <- 0.03
allele_freqs_v4$tra[6] <- 1 - allele_freqs_v4$traD[6]
allele_freqs_v4$IIIM[6] <- 0.26
allele_freqs_v4$III[6] <- 1 - allele_freqs_v4$IIIM[6]

#7 PA: YM=0.6, traD=0.02, IIIM=0.025
allele_freqs_v4$YM[7] <- 0.6
allele_freqs_v4$X[7] <- 1 - allele_freqs_v4$YM[7]
allele_freqs_v4$IIIM[7] <- 0.025
allele_freqs_v4$III[7] <- 1 - allele_freqs_v4$IIIM[7]
allele_freqs_v4$traD[7] <- 0.02
allele_freqs_v4$tra[7] <- 1 - allele_freqs_v4$traD[7]

#8 TN: YM=0.36, traD=0.072, IIIM=0.11
allele_freqs_v4$YM[8] <- 0.36
allele_freqs_v4$X[8] <- 1 - allele_freqs_v4$YM[8]
allele_freqs_v4$IIIM[8] <- 0.11
allele_freqs_v4$III[8] <- 1 - allele_freqs_v4$IIIM[8]
allele_freqs_v4$traD[8] <- 0.072
allele_freqs_v4$tra[8] <- 1 - allele_freqs_v4$traD[8]

#9 TX: YM=0.45, IIIM=0.1, traD=0.035
allele_freqs_v4$YM[9] <- 0.45
allele_freqs_v4$X[9] <- 1 - allele_freqs_v4$YM[9]
allele_freqs_v4$IIIM[9] <- 0.01
allele_freqs_v4$III[9] <- 1 - allele_freqs_v4$IIIM[9]
allele_freqs_v4$traD[9] <- 0.035
allele_freqs_v4$tra[9] <- 1 - allele_freqs_v4$traD[9]


geno_freqs_v4 <- geno.freqs(allele_freqs_v4)
geno_freqs_v4_norm <- geno.freqs.norm(geno_freqs_v4)


# simulate 10 generations with naive starting frequencies to see if they are stable
CA_v4_10 <- simulator(geno_freqs_v4_norm[1,2:19], 10)
GA_v4_10 <- simulator(geno_freqs_v4_norm[3,2:19], 10)
KS_v4_10 <- simulator(geno_freqs_v4_norm[4,2:19], 10)
NE_v4_10 <- simulator(geno_freqs_v4_norm[6,2:19], 10)
PA_v4_10 <- simulator(geno_freqs_v4_norm[7,2:19], 10)
TN_v4_10 <- simulator(geno_freqs_v4_norm[8,2:19], 10)
TX_v4_10 <- simulator(geno_freqs_v4_norm[9,2:19], 10)

v4.df <- obs.exp.df(male_freqs, female_freqs, CA_v4_10, FL_v1_10, GA_v4_10, KS_v4_10, NC_v1_10, NE_v4_10, PA_v4_10, TN_v4_10, TX_v4_10)


## v5
allele_freqs_v5 <- allele_freqs_v4

#1 CA: 
# YM male frequency is slightly high
# traD slightly low
# YM=0.115, IIIM=0.39, traD=0.16
allele_freqs_v5$YM[1] <- 0.115
allele_freqs_v5$X[1] <- 1 - allele_freqs_v5$YM[1]
allele_freqs_v5$traD[1] <- 0.16
allele_freqs_v5$tra[1] <- 1 - allele_freqs_v5$traD[1]
allele_freqs_v5$IIIM[1] <- 0.39
allele_freqs_v5$III[1] <- 1 - allele_freqs_v5$IIIM[1]

# FL
# very good fit from naive

#3 GA: 
# YM slightly low
# traD low
# YM=0.37, traD=0.04, IIIM=0.1
allele_freqs_v5$YM[3] <- 0.37
allele_freqs_v5$X[3] <- 1 - allele_freqs_v5$YM[3]
allele_freqs_v5$traD[3] <- 0.04
allele_freqs_v5$tra[3] <- 1 - allele_freqs_v5$traD[3]
allele_freqs_v5$IIIM[3] <- 0.1
allele_freqs_v5$III[3] <- 1 - allele_freqs_v5$IIIM[3]

#4 KS: 
# YM frequency is slightly high
# traD frequency is slightly high
# start population YM=0.4705, traD=0.01, IIIM=0.009
allele_freqs_v5$YM[4] <- 0.4705
allele_freqs_v5$X[4] <- 1 - allele_freqs_v5$YM[4]
allele_freqs_v5$IIIM[4] <- 0.009
allele_freqs_v5$III[4] <- 1 - allele_freqs_v5$IIIM[4]
allele_freqs_v5$traD[4] <- 0.01
allele_freqs_v5$tra[4] <- 1 - allele_freqs_v5$traD[4]

# NC
# good fit from naive

#6 NE: 
# traD frequency is slightly high 
# YM frequency is good fit
# slightly lower IIIM
# start YM=0.24, traD=0.03, IIIM=0.26
allele_freqs_v5$YM[6] <- 0.24
allele_freqs_v5$X[6] <- 1 - allele_freqs_v5$YM[6]
allele_freqs_v5$traD[6] <- 0.03
allele_freqs_v5$tra[6] <- 1 - allele_freqs_v5$traD[6]
allele_freqs_v5$IIIM[6] <- 0.25
allele_freqs_v5$III[6] <- 1 - allele_freqs_v5$IIIM[6]

#7 PA: 
# YM frequency is low and traD frequency is too high in simulated population
# lower IIIM and lower traD, raise YM slightly
# YM=0.65, traD=0.015, IIIM=0.01
allele_freqs_v5$YM[7] <- 0.65
allele_freqs_v5$X[7] <- 1 - allele_freqs_v5$YM[7]
allele_freqs_v5$IIIM[7] <- 0.01
allele_freqs_v5$III[7] <- 1 - allele_freqs_v5$IIIM[7]
allele_freqs_v5$traD[7] <- 0.015
allele_freqs_v5$tra[7] <- 1 - allele_freqs_v5$traD[7]

#8 TN: 
# traD frequency is slightly low
# YM male frequency is slightly low
# YM=0.37, traD=0.075, IIIM=0.11
allele_freqs_v5$YM[8] <- 0.37
allele_freqs_v5$X[8] <- 1 - allele_freqs_v5$YM[8]
allele_freqs_v5$IIIM[8] <- 0.11
allele_freqs_v5$III[8] <- 1 - allele_freqs_v5$IIIM[8]
allele_freqs_v5$traD[8] <- 0.075
allele_freqs_v5$tra[8] <- 1 - allele_freqs_v5$traD[8]

#9 TX: 
# traD frequency is slightly high in simulated population
# YM male frequency is slightly high
# YM=0.44, IIIM=0.1, traD=0.0325
allele_freqs_v5$YM[9] <- 0.44
allele_freqs_v5$X[9] <- 1 - allele_freqs_v5$YM[9]
allele_freqs_v5$IIIM[9] <- 0.01
allele_freqs_v5$III[9] <- 1 - allele_freqs_v5$IIIM[9]
allele_freqs_v5$traD[9] <- 0.0325
allele_freqs_v5$tra[9] <- 1 - allele_freqs_v5$traD[9]


#############
# simulate 10 generations to see if they fit observations
geno_freqs_v5 <- geno.freqs(allele_freqs_v5)
geno_freqs_v5_norm <- geno.freqs.norm(geno_freqs_v5)

CA_v5_10 <- simulator(geno_freqs_v5_norm[1,2:19], 10)
GA_v5_10 <- simulator(geno_freqs_v5_norm[3,2:19], 10)
KS_v5_10 <- simulator(geno_freqs_v5_norm[4,2:19], 10)
NE_v5_10 <- simulator(geno_freqs_v5_norm[6,2:19], 10)
PA_v5_10 <- simulator(geno_freqs_v5_norm[7,2:19], 10)
TN_v5_10 <- simulator(geno_freqs_v5_norm[8,2:19], 10)
TX_v5_10 <- simulator(geno_freqs_v5_norm[9,2:19], 10)

v5.df <- obs.exp.df(male_freqs, female_freqs, CA_v5_10, FL_v1_10, GA_v5_10, KS_v5_10, NC_v1_10, NE_v5_10, PA_v5_10, TN_v5_10, TX_v5_10)


####### compare and update
allele_freqs_v6 <- allele_freqs_v5

#1 CA: 
# YM male frequency is slightly low
# traD slightly high
allele_freqs_v6$YM[1] <- 0.1175
allele_freqs_v6$X[1] <- 1 - allele_freqs_v6$YM[1]
allele_freqs_v6$traD[1] <- 0.15
allele_freqs_v6$tra[1] <- 1 - allele_freqs_v6$traD[1]
allele_freqs_v6$IIIM[1] <- 0.385
allele_freqs_v6$III[1] <- 1 - allele_freqs_v6$IIIM[1]

# FL
# very good fit from naive

#3 GA: 
# YM very low
# traD low
allele_freqs_v6$YM[3] <- 0.38
allele_freqs_v6$X[3] <- 1 - allele_freqs_v6$YM[3]
allele_freqs_v6$traD[3] <- 0.05
allele_freqs_v6$tra[3] <- 1 - allele_freqs_v6$traD[3]
allele_freqs_v6$IIIM[3] <- 0.15
allele_freqs_v6$III[3] <- 1 - allele_freqs_v6$IIIM[3]

#4 KS: 
# YM frequency is slightly high
# traD frequency is slightly high
allele_freqs_v6$YM[4] <- 0.47
allele_freqs_v6$X[4] <- 1 - allele_freqs_v6$YM[4]
allele_freqs_v6$IIIM[4] <- 0.009
allele_freqs_v6$III[4] <- 1 - allele_freqs_v6$IIIM[4]
allele_freqs_v6$traD[4] <- 0.01
allele_freqs_v6$tra[4] <- 1 - allele_freqs_v6$traD[4]

# NC
# good fit from naive

#6 NE: 
# traD is good fit
# YM is slightly high
allele_freqs_v6$YM[6] <- 0.235
allele_freqs_v6$X[6] <- 1 - allele_freqs_v6$YM[6]
allele_freqs_v6$traD[6] <- 0.03
allele_freqs_v6$tra[6] <- 1 - allele_freqs_v6$traD[6]
allele_freqs_v6$IIIM[6] <- 0.255
allele_freqs_v6$III[6] <- 1 - allele_freqs_v6$IIIM[6]

#7 PA: 
# YM frequency is slightly low 
# traD frequency is too high in simulated population
allele_freqs_v6$YM[7] <- 0.7
allele_freqs_v6$X[7] <- 1 - allele_freqs_v6$YM[7]
allele_freqs_v6$IIIM[7] <- 0.005
allele_freqs_v6$III[7] <- 1 - allele_freqs_v6$IIIM[7]
allele_freqs_v6$traD[7] <- 0.0125
allele_freqs_v6$tra[7] <- 1 - allele_freqs_v6$traD[7]

#8 TN: 
# traD frequency is slightly high
# YM male frequency is slightly low
allele_freqs_v6$YM[8] <- 0.37
allele_freqs_v6$X[8] <- 1 - allele_freqs_v6$YM[8]
allele_freqs_v6$IIIM[8] <- 0.105
allele_freqs_v6$III[8] <- 1 - allele_freqs_v6$IIIM[8]
allele_freqs_v6$traD[8] <- 0.075
allele_freqs_v6$tra[8] <- 1 - allele_freqs_v6$traD[8]

#9 TX: 
# traD frequency is slightly low in simulated population
# YM male frequency is slightly high
allele_freqs_v6$YM[9] <- 0.43
allele_freqs_v6$X[9] <- 1 - allele_freqs_v6$YM[9]
allele_freqs_v6$IIIM[9] <- 0.011
allele_freqs_v6$III[9] <- 1 - allele_freqs_v6$IIIM[9]
allele_freqs_v6$traD[9] <- 0.0325
allele_freqs_v6$tra[9] <- 1 - allele_freqs_v6$traD[9]


#############
# simulate 10 generations to see if they fit observations
geno_freqs_v6 <- geno.freqs(allele_freqs_v6)
geno_freqs_v6_norm <- geno.freqs.norm(geno_freqs_v6)

CA_v6_10 <- simulator(geno_freqs_v6_norm[1,2:19], 10)
GA_v6_10 <- simulator(geno_freqs_v6_norm[3,2:19], 10)
KS_v6_10 <- simulator(geno_freqs_v6_norm[4,2:19], 10)
NE_v6_10 <- simulator(geno_freqs_v6_norm[6,2:19], 10)
PA_v6_10 <- simulator(geno_freqs_v6_norm[7,2:19], 10)
TN_v6_10 <- simulator(geno_freqs_v6_norm[8,2:19], 10)
TX_v6_10 <- simulator(geno_freqs_v6_norm[9,2:19], 10)

v6.df <- obs.exp.df(male_freqs, female_freqs, CA_v6_10, FL_v1_10, GA_v6_10, KS_v6_10, NC_v1_10, NE_v6_10, PA_v6_10, TN_v6_10, TX_v6_10)


####### compare and update
allele_freqs_v7 <- allele_freqs_v6

#1 CA: 
# good fit! 

# FL
# very good fit from naive

#3 GA: 
# YM very low (raise YM)
# traD high (lower IIIM)
allele_freqs_v7$YM[3] <- 0.40
allele_freqs_v7$X[3] <- 1 - allele_freqs_v7$YM[3]
allele_freqs_v7$traD[3] <- 0.05
allele_freqs_v7$tra[3] <- 1 - allele_freqs_v7$traD[3]
allele_freqs_v7$IIIM[3] <- 0.125
allele_freqs_v7$III[3] <- 1 - allele_freqs_v7$IIIM[3]

#4 KS: 
# YM frequency is slightly high
# traD frequency is slightly high
allele_freqs_v7$YM[4] <- 0.46
allele_freqs_v7$X[4] <- 1 - allele_freqs_v7$YM[4]
allele_freqs_v7$IIIM[4] <- 0.009
allele_freqs_v7$III[4] <- 1 - allele_freqs_v7$IIIM[4]
allele_freqs_v7$traD[4] <- 0.01
allele_freqs_v7$tra[4] <- 1 - allele_freqs_v7$traD[4]

# NC
# good fit from naive

#6 NE: 
# Good fit! 

#7 PA: 
# YM frequency is slightly low
# traD frequency is too high in simulated population
allele_freqs_v7$YM[7] <- 0.71
allele_freqs_v7$X[7] <- 1 - allele_freqs_v7$YM[7]
allele_freqs_v7$IIIM[7] <- 0.001
allele_freqs_v7$III[7] <- 1 - allele_freqs_v7$IIIM[7]
allele_freqs_v7$traD[7] <- 0.01
allele_freqs_v7$tra[7] <- 1 - allele_freqs_v7$traD[7]

#8 TN: 
# traD frequency is slightly high
# YM male frequency is good fit
allele_freqs_v7$YM[8] <- 0.37
allele_freqs_v7$X[8] <- 1 - allele_freqs_v7$YM[8]
allele_freqs_v7$IIIM[8] <- 0.1025
allele_freqs_v7$III[8] <- 1 - allele_freqs_v7$IIIM[8]
allele_freqs_v7$traD[8] <- 0.075
allele_freqs_v7$tra[8] <- 1 - allele_freqs_v7$traD[8]

#9 TX: 
# traD frequency is slightly low in simulated population
# YM male frequency is slightly high
allele_freqs_v7$YM[9] <- 0.425
allele_freqs_v7$X[9] <- 1 - allele_freqs_v7$YM[9]
allele_freqs_v7$IIIM[9] <- 0.012
allele_freqs_v7$III[9] <- 1 - allele_freqs_v7$IIIM[9]
allele_freqs_v7$traD[9] <- 0.0325
allele_freqs_v7$tra[9] <- 1 - allele_freqs_v7$traD[9]


#############
# simulate 10 generations to see if they fit observations
geno_freqs_v7 <- geno.freqs(allele_freqs_v7)
geno_freqs_v7_norm <- geno.freqs.norm(geno_freqs_v7)

GA_v7_10 <- simulator(geno_freqs_v7_norm[3,2:19], 10)
KS_v7_10 <- simulator(geno_freqs_v7_norm[4,2:19], 10)
PA_v7_10 <- simulator(geno_freqs_v7_norm[7,2:19], 10)
TN_v7_10 <- simulator(geno_freqs_v7_norm[8,2:19], 10)
TX_v7_10 <- simulator(geno_freqs_v7_norm[9,2:19], 10)

v7.df <- obs.exp.df(male_freqs, female_freqs, CA_v6_10, FL_v1_10, GA_v7_10, KS_v7_10, NC_v1_10, NE_v6_10, PA_v7_10, TN_v7_10, TX_v7_10)


####### compare and update
allele_freqs_v8 <- allele_freqs_v7

#1 CA: 
# good fit! 

# FL
# very good fit from naive

#3 GA: 
# YM very low (raise YM)
# traD high (lower IIIM)
allele_freqs_v8$YM[3] <- 0.425
allele_freqs_v8$X[3] <- 1 - allele_freqs_v8$YM[3]
allele_freqs_v8$traD[3] <- 0.05
allele_freqs_v8$tra[3] <- 1 - allele_freqs_v8$traD[3]
allele_freqs_v8$IIIM[3] <- 0.1
allele_freqs_v8$III[3] <- 1 - allele_freqs_v8$IIIM[3]

#4 KS: 
# YM frequency is slightly high
# traD frequency is slightly high
allele_freqs_v8$YM[4] <- 0.45
allele_freqs_v8$X[4] <- 1 - allele_freqs_v8$YM[4]
allele_freqs_v8$IIIM[4] <- 0.009
allele_freqs_v8$III[4] <- 1 - allele_freqs_v8$IIIM[4]
allele_freqs_v8$traD[4] <- 0.01
allele_freqs_v8$tra[4] <- 1 - allele_freqs_v8$traD[4]

# NC
# good fit from naive

#6 NE: 
# Good fit! 

#7 PA: 
# YM frequency is good fit 
# traD frequency is too high in simulated population
# Revert back to values similar to v1 because that was best fit
allele_freqs_v8$YM[7] <- 0.5
allele_freqs_v8$X[7] <- 1 - allele_freqs_v8$YM[7]
allele_freqs_v8$IIIM[7] <- 0.0001
allele_freqs_v8$III[7] <- 1 - allele_freqs_v8$IIIM[7]
allele_freqs_v8$traD[7] <- 0.02
allele_freqs_v8$tra[7] <- 1 - allele_freqs_v8$traD[7]

#8 TN: 
# traD frequency is good fit
# YM male frequency is slightly high
allele_freqs_v8$YM[8] <- 0.365
allele_freqs_v8$X[8] <- 1 - allele_freqs_v8$YM[8]
allele_freqs_v8$IIIM[8] <- 0.1025
allele_freqs_v8$III[8] <- 1 - allele_freqs_v8$IIIM[8]
allele_freqs_v8$traD[8] <- 0.075
allele_freqs_v8$tra[8] <- 1 - allele_freqs_v8$traD[8]

#9 TX: 
# traD frequency is slightly low in simulated population
# YM male frequency is good fit, but might need to lower slightly to get other freqs to obs
allele_freqs_v8$YM[9] <- 0.42
allele_freqs_v8$X[9] <- 1 - allele_freqs_v8$YM[9]
allele_freqs_v8$IIIM[9] <- 0.0125
allele_freqs_v8$III[9] <- 1 - allele_freqs_v8$IIIM[9]
allele_freqs_v8$traD[9] <- 0.035
allele_freqs_v8$tra[9] <- 1 - allele_freqs_v8$traD[9]


#############
# simulate 10 generations to see if they fit observations
geno_freqs_v8 <- geno.freqs(allele_freqs_v8)
geno_freqs_v8_norm <- geno.freqs.norm(geno_freqs_v8)

GA_v8_10 <- simulator(geno_freqs_v8_norm[3,2:19], 10)
KS_v8_10 <- simulator(geno_freqs_v8_norm[4,2:19], 10)
PA_v8_10 <- simulator(geno_freqs_v8_norm[7,2:19], 10)
TN_v8_10 <- simulator(geno_freqs_v8_norm[8,2:19], 10)
TX_v8_10 <- simulator(geno_freqs_v8_norm[9,2:19], 10)

v8.df <- obs.exp.df(male_freqs, female_freqs, CA_v6_10, FL_v1_10, GA_v8_10, KS_v8_10, NC_v1_10, NE_v6_10, PA_v8_10, TN_v8_10, TX_v8_10)


####### compare and update
allele_freqs_v9 <- allele_freqs_v8

#1 CA: 
# good fit! 

# FL
# very good fit from naive

#3 GA: 
# YM  low (raise YM)
# traD high (lower IIIM)
allele_freqs_v9$YM[3] <- 0.44
allele_freqs_v9$X[3] <- 1 - allele_freqs_v9$YM[3]
allele_freqs_v9$traD[3] <- 0.05
allele_freqs_v9$tra[3] <- 1 - allele_freqs_v9$traD[3]
allele_freqs_v9$IIIM[3] <- 0.05
allele_freqs_v9$III[3] <- 1 - allele_freqs_v9$IIIM[3]

#4 KS: 
# YM frequency is slightly high
# traD frequency is slightly high
allele_freqs_v9$YM[4] <- 0.44
allele_freqs_v9$X[4] <- 1 - allele_freqs_v9$YM[4]
allele_freqs_v9$IIIM[4] <- 0.009
allele_freqs_v9$III[4] <- 1 - allele_freqs_v9$IIIM[4]
allele_freqs_v9$traD[4] <- 0.01
allele_freqs_v9$tra[4] <- 1 - allele_freqs_v9$traD[4]

# NC
# good fit from naive

#6 NE: 
# Good fit! 

#7 PA: 
# YM frequency is good fit 
# traD frequency is too high (lower IIIM more)
allele_freqs_v9$YM[7] <- 0.5
allele_freqs_v9$X[7] <- 1 - allele_freqs_v9$YM[7]
allele_freqs_v9$IIIM[7] <- 0.00005
allele_freqs_v9$III[7] <- 1 - allele_freqs_v9$IIIM[7]
allele_freqs_v9$traD[7] <- 0.02
allele_freqs_v9$tra[7] <- 1 - allele_freqs_v9$traD[7]

#8 TN: 
# traD frequency is slightly low - raise IIIM and traD
# YM male frequency is good fit, but lower lower YM if raising IIIM
allele_freqs_v9$YM[8] <- 0.3625
allele_freqs_v9$X[8] <- 1 - allele_freqs_v9$YM[8]
allele_freqs_v9$IIIM[8] <- 0.103
allele_freqs_v9$III[8] <- 1 - allele_freqs_v9$IIIM[8]
allele_freqs_v9$traD[8] <- 0.08
allele_freqs_v9$tra[8] <- 1 - allele_freqs_v9$traD[8]

#9 TX: 
# traD frequency is slightly low - raise traD and IIIM
# YM male frequency is good fit 
allele_freqs_v9$YM[9] <- 0.42
allele_freqs_v9$X[9] <- 1 - allele_freqs_v9$YM[9]
allele_freqs_v9$IIIM[9] <- 0.0127
allele_freqs_v9$III[9] <- 1 - allele_freqs_v9$IIIM[9]
allele_freqs_v9$traD[9] <- 0.037
allele_freqs_v9$tra[9] <- 1 - allele_freqs_v9$traD[9]


#############
# simulate 10 generations to see if they fit observations
geno_freqs_v9 <- geno.freqs(allele_freqs_v9)
geno_freqs_v9_norm <- geno.freqs.norm(geno_freqs_v9)

GA_v9_10 <- simulator(geno_freqs_v9_norm[3,2:19], 10)
KS_v9_10 <- simulator(geno_freqs_v9_norm[4,2:19], 10)
PA_v9_10 <- simulator(geno_freqs_v9_norm[7,2:19], 10)
TN_v9_10 <- simulator(geno_freqs_v9_norm[8,2:19], 10)
TX_v9_10 <- simulator(geno_freqs_v9_norm[9,2:19], 10)

v9.df <- obs.exp.df(male_freqs, female_freqs, CA_v6_10, FL_v1_10, GA_v9_10, KS_v9_10, NC_v1_10, NE_v6_10, PA_v9_10, TN_v9_10, TX_v9_10)


####### compare and update
allele_freqs_v10 <- allele_freqs_v9

#1 CA: 
# good fit! 

# FL
# very good fit from naive

#3 GA: 
# YM  good fit
# traD slightly high (lower IIIM)
allele_freqs_v10$YM[3] <- 0.44
allele_freqs_v10$X[3] <- 1 - allele_freqs_v10$YM[3]
allele_freqs_v10$traD[3] <- 0.04
allele_freqs_v10$tra[3] <- 1 - allele_freqs_v10$traD[3]
allele_freqs_v10$IIIM[3] <- 0.05
allele_freqs_v10$III[3] <- 1 - allele_freqs_v10$IIIM[3]

#4 KS: 
# YM frequency is good slightly high - lower YM
# traD frequency is good fit
allele_freqs_v10$YM[4] <- 0.43
allele_freqs_v10$X[4] <- 1 - allele_freqs_v10$YM[4]
allele_freqs_v10$IIIM[4] <- 0.009
allele_freqs_v10$III[4] <- 1 - allele_freqs_v10$IIIM[4]
allele_freqs_v10$traD[4] <- 0.01
allele_freqs_v10$tra[4] <- 1 - allele_freqs_v10$traD[4]

# NC
# good fit from naive

#6 NE: 
# Good fit! 

#7 PA: 
# YM frequency is good fit 
# traD frequency is too high (lower IIIM more)
allele_freqs_v10$YM[7] <- 0.5
allele_freqs_v10$X[7] <- 1 - allele_freqs_v10$YM[7]
allele_freqs_v10$IIIM[7] <- 0.00001
allele_freqs_v10$III[7] <- 1 - allele_freqs_v10$IIIM[7]
allele_freqs_v10$traD[7] <- 0.02
allele_freqs_v10$tra[7] <- 1 - allele_freqs_v10$traD[7]

#8 TN: 
# traD frequency is good fit (slightly high) - lower IIIM 
# YM male frequency is good fit - but lower YM to get traD down
# v6 and v7 were the best fit
allele_freqs_v10$YM[8] <- 0.37
allele_freqs_v10$X[8] <- 1 - allele_freqs_v10$YM[8]
allele_freqs_v10$IIIM[8] <- 0.102
allele_freqs_v10$III[8] <- 1 - allele_freqs_v10$IIIM[8]
allele_freqs_v10$traD[8] <- 0.09
allele_freqs_v10$tra[8] <- 1 - allele_freqs_v10$traD[8]

#9 TX: 
# traD frequency is good fit  
# YM male frequency is good fit 


#############
# simulate 10 generations to see if they fit observations
geno_freqs_v10 <- geno.freqs(allele_freqs_v10)
geno_freqs_v10_norm <- geno.freqs.norm(geno_freqs_v10)

GA_v10_10 <- simulator(geno_freqs_v10_norm[3,2:19], 10)
KS_v10_10 <- simulator(geno_freqs_v10_norm[4,2:19], 10)
PA_v10_10 <- simulator(geno_freqs_v10_norm[7,2:19], 10)
TN_v10_10 <- simulator(geno_freqs_v10_norm[8,2:19], 10)

v10.df <- obs.exp.df(male_freqs, female_freqs, CA_v6_10, FL_v1_10, GA_v10_10, KS_v10_10, NC_v1_10, NE_v6_10, PA_v10_10, TN_v10_10, TX_v9_10)


####### compare and update
allele_freqs_v11 <- allele_freqs_v10

#1 CA: 
# good fit! 

# FL
# very good fit from naive

#3 GA: 
# YM  slightly low - raise YM
# traD slightly low (raise IIIM and traD)
allele_freqs_v11$YM[3] <- 0.445
allele_freqs_v11$X[3] <- 1 - allele_freqs_v11$YM[3]
allele_freqs_v11$traD[3] <- 0.045
allele_freqs_v11$tra[3] <- 1 - allele_freqs_v11$traD[3]
allele_freqs_v11$IIIM[3] <- 0.075
allele_freqs_v11$III[3] <- 1 - allele_freqs_v11$IIIM[3]

#4 KS: 
# YM frequency is slightly high - slightly lower YM
# traD frequency is good fit (slightly raise IIIM to compensate for lower YM)
allele_freqs_v11$YM[4] <- 0.425
allele_freqs_v11$X[4] <- 1 - allele_freqs_v11$YM[4]
allele_freqs_v11$IIIM[4] <- 0.0095
allele_freqs_v11$III[4] <- 1 - allele_freqs_v11$IIIM[4]
allele_freqs_v11$traD[4] <- 0.01
allele_freqs_v11$tra[4] <- 1 - allele_freqs_v11$traD[4]

# NC
# good fit from naive

#6 NE: 
# Good fit! 

#7 PA: 
# YM frequency is good fit 
# traD frequency is too high (lower IIIM more, and lower traD)
allele_freqs_v11$YM[7] <- 0.5
allele_freqs_v11$X[7] <- 1 - allele_freqs_v11$YM[7]
allele_freqs_v11$IIIM[7] <- 0.000001
allele_freqs_v11$III[7] <- 1 - allele_freqs_v11$IIIM[7]
allele_freqs_v11$traD[7] <- 0.015
allele_freqs_v11$tra[7] <- 1 - allele_freqs_v11$traD[7]

#8 TN: 
# traD frequency is high
# YM male frequency is high
# revert to v6-8
#   Pop     YM      X   IIIM    III   traD    tra
#v6  TN 0.3700 0.6300 0.1050 0.8950 0.0750 0.9250
#v7  TN 0.3700 0.6300 0.1025 0.8975 0.0750 0.9250
#v8  TN 0.3650 0.6350 0.1025 0.8975 0.0750 0.9250
allele_freqs_v11$YM[8] <- 0.3675
allele_freqs_v11$X[8] <- 1 - allele_freqs_v11$YM[8]
allele_freqs_v11$IIIM[8] <- 0.103
allele_freqs_v11$III[8] <- 1 - allele_freqs_v11$IIIM[8]
allele_freqs_v11$traD[8] <- 0.075
allele_freqs_v11$tra[8] <- 1 - allele_freqs_v11$traD[8]

#9 TX: 
# good fit


#############
# simulate 10 generations to see if they fit observations
geno_freqs_v11 <- geno.freqs(allele_freqs_v11)
geno_freqs_v11_norm <- geno.freqs.norm(geno_freqs_v11)

GA_v11_10 <- simulator(geno_freqs_v11_norm[3,2:19], 10)
KS_v11_10 <- simulator(geno_freqs_v11_norm[4,2:19], 10)
PA_v11_10 <- simulator(geno_freqs_v11_norm[7,2:19], 10)
TN_v11_10 <- simulator(geno_freqs_v11_norm[8,2:19], 10)

v11.df <- obs.exp.df(male_freqs, female_freqs, CA_v6_10, FL_v1_10, GA_v11_10, KS_v11_10, NC_v1_10, NE_v6_10, PA_v11_10, TN_v11_10, TX_v9_10)


####### compare and update
allele_freqs_v12 <- allele_freqs_v11

#1 CA: 
# good fit! 

# FL
# very good fit from v1

#3 GA: 
# YM too low
# traD too high
# revert to v9 and v10
#    Pop     YM      X    IIIM     III   traD    tra
#v9   GA 0.4400 0.5600 0.05000 0.95000 0.0500 0.9500 # traD too high
#v10  GA 0.4400 0.5600 0.05000 0.95000 0.0400 0.9600 # traD too low
allele_freqs_v12$YM[3] <- 0.44
allele_freqs_v12$X[3] <- 1 - allele_freqs_v12$YM[3]
allele_freqs_v12$traD[3] <- 0.045
allele_freqs_v12$tra[3] <- 1 - allele_freqs_v12$traD[3]
allele_freqs_v12$IIIM[3] <- 0.05
allele_freqs_v12$III[3] <- 1 - allele_freqs_v12$IIIM[3]

#4 KS: 
# Good fit! 

# NC
# good fit from v1

#6 NE: 
# Good fit! 

#7 PA: 
# YM frequency is good fit 
# traD frequency is too low (raise IIIM and traD)
allele_freqs_v12$YM[7] <- 0.5
allele_freqs_v12$X[7] <- 1 - allele_freqs_v12$YM[7]
allele_freqs_v12$IIIM[7] <- 0.000005
allele_freqs_v12$III[7] <- 1 - allele_freqs_v12$IIIM[7]
allele_freqs_v12$traD[7] <- 0.0175
allele_freqs_v12$tra[7] <- 1 - allele_freqs_v12$traD[7]

#8 TN: 
# traD frequency is good fit, but raise if lowering YM
# YM male frequency is slightly high, lower YM
allele_freqs_v12$YM[8] <- 0.365
allele_freqs_v12$X[8] <- 1 - allele_freqs_v12$YM[8]
allele_freqs_v12$IIIM[8] <- 0.103
allele_freqs_v12$III[8] <- 1 - allele_freqs_v12$IIIM[8]
allele_freqs_v12$traD[8] <- 0.076
allele_freqs_v12$tra[8] <- 1 - allele_freqs_v12$traD[8]


#9 TX: 
# good fit


#############
# simulate 10 generations to see if they fit observations
geno_freqs_v12 <- geno.freqs(allele_freqs_v12)
geno_freqs_v12_norm <- geno.freqs.norm(geno_freqs_v12)

GA_v12_10 <- simulator(geno_freqs_v12_norm[3,2:19], 10)
PA_v12_10 <- simulator(geno_freqs_v12_norm[7,2:19], 10)
TN_v12_10 <- simulator(geno_freqs_v12_norm[8,2:19], 10)

v12.df <- obs.exp.df(male_freqs, female_freqs, CA_v6_10, FL_v1_10, GA_v12_10, KS_v11_10, NC_v1_10, NE_v6_10, PA_v12_10, TN_v12_10, TX_v9_10)


####### compare and update
allele_freqs_v13 <- allele_freqs_v12

#1 CA: 
# good fit! 

# FL
# very good fit from v1

#3 GA: 
# YM too low - very slight raise YM
# traD too low (raise traD and IIIM)
allele_freqs_v13$YM[3] <- 0.441
allele_freqs_v13$X[3] <- 1 - allele_freqs_v13$YM[3]
allele_freqs_v13$traD[3] <- 0.0475
allele_freqs_v13$tra[3] <- 1 - allele_freqs_v13$traD[3]
allele_freqs_v13$IIIM[3] <- 0.055
allele_freqs_v13$III[3] <- 1 - allele_freqs_v13$IIIM[3]

#4 KS: 
# Good fit! 

# NC
# good fit from v1

#6 NE: 
# Good fit! 

#7 PA: 
# YM frequency is good fit 
# traD frequency is too high (lower IIIM )
allele_freqs_v13$YM[7] <- 0.5
allele_freqs_v13$X[7] <- 1 - allele_freqs_v13$YM[7]
allele_freqs_v13$IIIM[7] <- 0.0000025
allele_freqs_v13$III[7] <- 1 - allele_freqs_v13$IIIM[7]
allele_freqs_v13$traD[7] <- 0.0175
allele_freqs_v13$tra[7] <- 1 - allele_freqs_v13$traD[7]

#8 TN: 
# good fit! 

#9 TX: 
# good fit! 


#############
# simulate 10 generations to see if they fit observations
geno_freqs_v13 <- geno.freqs(allele_freqs_v13)
geno_freqs_v13_norm <- geno.freqs.norm(geno_freqs_v13)

GA_v13_10 <- simulator(geno_freqs_v13_norm[3,2:19], 10)
PA_v13_10 <- simulator(geno_freqs_v13_norm[7,2:19], 10)

v13.df <- obs.exp.df(male_freqs, female_freqs, CA_v6_10, FL_v1_10, GA_v13_10, KS_v11_10, NC_v1_10, NE_v6_10, PA_v13_10, TN_v12_10, TX_v9_10)


####### compare and update
allele_freqs_v14 <- allele_freqs_v13

#1 CA: 
# good fit! 

# FL
# very good fit from v1

#3 GA: 
# YM too low - raise slightly
# traD slightly high - lower traD and and IIIM
allele_freqs_v14$YM[3] <- 0.442
allele_freqs_v14$X[3] <- 1 - allele_freqs_v14$YM[3]
allele_freqs_v14$traD[3] <- 0.0474
allele_freqs_v14$tra[3] <- 1 - allele_freqs_v14$traD[3]
allele_freqs_v14$IIIM[3] <- 0.054
allele_freqs_v14$III[3] <- 1 - allele_freqs_v14$IIIM[3]

#4 KS: 
# Good fit! 

# NC
# good fit from v1

#6 NE: 
# Good fit! 

#7 PA: 
# YM frequency is good fit 
# traD frequency is too high (lower IIIM )
allele_freqs_v14$YM[7] <- 0.5
allele_freqs_v14$X[7] <- 1 - allele_freqs_v14$YM[7]
allele_freqs_v14$IIIM[7] <- 0.000002
allele_freqs_v14$III[7] <- 1 - allele_freqs_v14$IIIM[7]
allele_freqs_v14$traD[7] <- 0.0175
allele_freqs_v14$tra[7] <- 1 - allele_freqs_v14$traD[7]

#8 TN: 
# good fit

#9 TX: 
# good fit! 


#############
# simulate 10 generations to see if they fit observations
geno_freqs_v14 <- geno.freqs(allele_freqs_v14)
geno_freqs_v14_norm <- geno.freqs.norm(geno_freqs_v14)

GA_v14_10 <- simulator(geno_freqs_v14_norm[3,2:19], 10)
PA_v14_10 <- simulator(geno_freqs_v14_norm[7,2:19], 10)

v14.df <- obs.exp.df(male_freqs, female_freqs, CA_v6_10, FL_v1_10, GA_v14_10, KS_v11_10, NC_v1_10, NE_v6_10, PA_v14_10, TN_v12_10, TX_v9_10)


####### compare and update
allele_freqs_v15 <- allele_freqs_v14

#1 CA: 
# good fit! 

# FL
# very good fit from v1

#3 GA: 
# YM too low - raise slightly
# traD slightly high - lower traD and and IIIM
allele_freqs_v15$YM[3] <- 0.443
allele_freqs_v15$X[3] <- 1 - allele_freqs_v15$YM[3]
allele_freqs_v15$traD[3] <- 0.0473
allele_freqs_v15$tra[3] <- 1 - allele_freqs_v15$traD[3]
allele_freqs_v15$IIIM[3] <- 0.053
allele_freqs_v15$III[3] <- 1 - allele_freqs_v15$IIIM[3]

#4 KS: 
# Good fit! 

# NC
# good fit from v1

#6 NE: 
# Good fit! 

#7 PA: 
# YM frequency is good fit 
# traD frequency is slightly high (lower IIIM and traD)
allele_freqs_v15$YM[7] <- 0.5
allele_freqs_v15$X[7] <- 1 - allele_freqs_v15$YM[7]
allele_freqs_v15$IIIM[7] <- 0.0000015
allele_freqs_v15$III[7] <- 1 - allele_freqs_v15$IIIM[7]
allele_freqs_v15$traD[7] <- 0.01725
allele_freqs_v15$tra[7] <- 1 - allele_freqs_v15$traD[7]

#8 TN: 
# good fit

#9 TX: 
# good fit! 


#############
# simulate 10 generations to see if they fit observations
geno_freqs_v15 <- geno.freqs(allele_freqs_v15)
geno_freqs_v15_norm <- geno.freqs.norm(geno_freqs_v15)

GA_v15_10 <- simulator(geno_freqs_v15_norm[3,2:19], 10)
PA_v15_10 <- simulator(geno_freqs_v15_norm[7,2:19], 10)

v15.df <- obs.exp.df(male_freqs, female_freqs, CA_v6_10, FL_v1_10, GA_v15_10, KS_v11_10, NC_v1_10, NE_v6_10, PA_v15_10, TN_v12_10, TX_v9_10)


####### compare and update
allele_freqs_v16 <- allele_freqs_v15

#1 CA: 
# good fit! 

# FL
# very good fit from v1

#3 GA: 
# YM slightly low - raise slightly
# traD slightly high - lower traD and and IIIM
allele_freqs_v16$YM[3] <- 0.444
allele_freqs_v16$X[3] <- 1 - allele_freqs_v16$YM[3]
allele_freqs_v16$traD[3] <- 0.0472
allele_freqs_v16$tra[3] <- 1 - allele_freqs_v16$traD[3]
allele_freqs_v16$IIIM[3] <- 0.052
allele_freqs_v16$III[3] <- 1 - allele_freqs_v16$IIIM[3]

#4 KS: 
# Good fit! 

# NC
# good fit from v1

#6 NE: 
# Good fit! 

#7 PA: 
# YM frequency is good fit 
# traD frequency is slightly high (lower IIIM and traD)
allele_freqs_v16$YM[7] <- 0.5
allele_freqs_v16$X[7] <- 1 - allele_freqs_v16$YM[7]
allele_freqs_v16$IIIM[7] <- 0.00000125
allele_freqs_v16$III[7] <- 1 - allele_freqs_v16$IIIM[7]
allele_freqs_v16$traD[7] <- 0.0172
allele_freqs_v16$tra[7] <- 1 - allele_freqs_v16$traD[7]

#8 TN: 
# good fit

#9 TX: 
# good fit! 


#############
# simulate 10 generations to see if they fit observations
geno_freqs_v16 <- geno.freqs(allele_freqs_v16)
geno_freqs_v16_norm <- geno.freqs.norm(geno_freqs_v16)

GA_v16_10 <- simulator(geno_freqs_v16_norm[3,2:19], 10)
PA_v16_10 <- simulator(geno_freqs_v16_norm[7,2:19], 10)

v16.df <- obs.exp.df(male_freqs, female_freqs, CA_v6_10, FL_v1_10, GA_v16_10, KS_v11_10, NC_v1_10, NE_v6_10, PA_v16_10, TN_v12_10, TX_v9_10)


####### compare and update
allele_freqs_v17 <- allele_freqs_v16

#1 CA: 
# good fit! 

# FL
# very good fit from v1

#3 GA: 
# YM slightly low - raise slightly
# traD slightly high - lower traD and and IIIM
allele_freqs_v17$YM[3] <- 0.4445
allele_freqs_v17$X[3] <- 1 - allele_freqs_v17$YM[3]
allele_freqs_v17$traD[3] <- 0.0471
allele_freqs_v17$tra[3] <- 1 - allele_freqs_v17$traD[3]
allele_freqs_v17$IIIM[3] <- 0.051
allele_freqs_v17$III[3] <- 1 - allele_freqs_v17$IIIM[3]

#4 KS: 
# Good fit! 

# NC
# good fit from v1

#6 NE: 
# Good fit! 

#7 PA: 
# YM frequency is good fit 
# traD frequency is slightly high (lower IIIM and traD)
allele_freqs_v17$YM[7] <- 0.5
allele_freqs_v17$X[7] <- 1 - allele_freqs_v17$YM[7]
allele_freqs_v17$IIIM[7] <- 0.0000012
allele_freqs_v17$III[7] <- 1 - allele_freqs_v17$IIIM[7]
allele_freqs_v17$traD[7] <- 0.0171
allele_freqs_v17$tra[7] <- 1 - allele_freqs_v17$traD[7]

#8 TN: 
# good fit

#9 TX: 
# good fit! 


#############
# simulate 10 generations to see if they fit observations
geno_freqs_v17 <- geno.freqs(allele_freqs_v17)
geno_freqs_v17_norm <- geno.freqs.norm(geno_freqs_v17)

GA_v17_10 <- simulator(geno_freqs_v17_norm[3,2:19], 10)
PA_v17_10 <- simulator(geno_freqs_v17_norm[7,2:19], 10)

v17.df <- obs.exp.df(male_freqs, female_freqs, CA_v6_10, FL_v1_10, GA_v17_10, KS_v11_10, NC_v1_10, NE_v6_10, PA_v17_10, TN_v12_10, TX_v9_10)


####### compare and update
allele_freqs_v18 <- allele_freqs_v17

#1 CA: 
# good fit! 

# FL
# very good fit from v1

#3 GA: 
# YM slightly low - raise slightly
# traD slightly high - lower traD and and IIIM
# looks like v13-16 (v12 has traD low)
allele_freqs_v18$YM[3] <- 0.445
allele_freqs_v18$X[3] <- 1 - allele_freqs_v18$YM[3]
allele_freqs_v18$traD[3] <- 0.047
allele_freqs_v18$tra[3] <- 1 - allele_freqs_v18$traD[3]
allele_freqs_v18$IIIM[3] <- 0.0505
allele_freqs_v18$III[3] <- 1 - allele_freqs_v18$IIIM[3]

#4 KS: 
# Good fit! 

# NC
# good fit from v1

#6 NE: 
# Good fit! 

#7 PA: 
# YM frequency is good fit 
# traD frequency is slightly high (lower IIIM and traD)
# looks like v12-v16 (v11 has traD low)
allele_freqs_v18$YM[7] <- 0.5
allele_freqs_v18$X[7] <- 1 - allele_freqs_v18$YM[7]
allele_freqs_v18$IIIM[7] <- 0.000001
allele_freqs_v18$III[7] <- 1 - allele_freqs_v18$IIIM[7]
allele_freqs_v18$traD[7] <- 0.0165
allele_freqs_v18$tra[7] <- 1 - allele_freqs_v18$traD[7]

#8 TN: 
# good fit

#9 TX: 
# good fit! 


#############
# simulate 10 generations to see if they fit observations
geno_freqs_v18 <- geno.freqs(allele_freqs_v18)
geno_freqs_v18_norm <- geno.freqs.norm(geno_freqs_v18)

GA_v18_10 <- simulator(geno_freqs_v18_norm[3,2:19], 10)
PA_v18_10 <- simulator(geno_freqs_v18_norm[7,2:19], 10)

v18.df <- obs.exp.df(male_freqs, female_freqs, CA_v6_10, FL_v1_10, GA_v18_10, KS_v11_10, NC_v1_10, NE_v6_10, PA_v18_10, TN_v12_10, TX_v9_10)


####### compare and update
allele_freqs_v19 <- allele_freqs_v18

#1 CA: 
# good fit! 

# FL
# very good fit from v1

#3 GA: 
# YM good fit - raise slightly if lowering IIIM
# traD slightly high - lower traD and and IIIM
# looks like v13-17 (v12 has traD low)
# subset(rbind(allele_freqs_v12, allele_freqs_v13, allele_freqs_v14, allele_freqs_v15, allele_freqs_v16, allele_freqs_v17, allele_freqs_v18), Pop=="GA")
allele_freqs_v19$YM[3] <- 0.4455
allele_freqs_v19$X[3] <- 1 - allele_freqs_v19$YM[3]
allele_freqs_v19$traD[3] <- 0.046
allele_freqs_v19$tra[3] <- 1 - allele_freqs_v19$traD[3]
allele_freqs_v19$IIIM[3] <- 0.050
allele_freqs_v19$III[3] <- 1 - allele_freqs_v19$IIIM[3]


#4 KS: 
# Good fit! 

# NC
# good fit from v1

#6 NE: 
# Good fit! 

#7 PA: 
# YM frequency is good fit 
# traD frequency is slightly high (lower traD)
# looks like v12-v17 (v11 has traD low)
# subset(rbind(allele_freqs_v11, allele_freqs_v12, allele_freqs_v13, allele_freqs_v14, allele_freqs_v15, allele_freqs_v16, allele_freqs_v17, allele_freqs_v18), Pop=="PA")
allele_freqs_v19$YM[7] <- 0.5
allele_freqs_v19$X[7] <- 1 - allele_freqs_v19$YM[7]
allele_freqs_v19$IIIM[7] <- 0.000001
allele_freqs_v19$III[7] <- 1 - allele_freqs_v19$IIIM[7]
allele_freqs_v19$traD[7] <- 0.01575
allele_freqs_v19$tra[7] <- 1 - allele_freqs_v19$traD[7]


#8 TN: 
# good fit

#9 TX: 
# good fit! 


#############
# simulate 10 generations to see if they fit observations
geno_freqs_v19 <- geno.freqs(allele_freqs_v19)
geno_freqs_v19_norm <- geno.freqs.norm(geno_freqs_v19)

GA_v19_10 <- simulator(geno_freqs_v19_norm[3,2:19], 10)
PA_v19_10 <- simulator(geno_freqs_v19_norm[7,2:19], 10)

v19.df <- obs.exp.df(male_freqs, female_freqs, CA_v6_10, FL_v1_10, GA_v19_10, KS_v11_10, NC_v1_10, NE_v6_10, PA_v19_10, TN_v12_10, TX_v9_10)


####### compare and update
allele_freqs_v20 <- allele_freqs_v19

#1 CA: 
# good fit! 

# FL
# very good fit from v1

#3 GA: 
# YM good fit (slightly low) - raise slightly
# traD good fit

#4 KS: 
# Good fit! 

# NC
# good fit from v1

#6 NE: 
# Good fit! 

#7 PA: 
# good fit

#8 TN: 
# good fit

#9 TX: 
# good fit! 


#######
# Perform final simulation for record

CA_final <- simulator(geno_freqs_v19_norm[1,2:19], 10)
FL_final <- simulator(geno_freqs_v19_norm[2,2:19], 10)
GA_final <- simulator(geno_freqs_v19_norm[3,2:19], 10)
KS_final <- simulator(geno_freqs_v19_norm[4,2:19], 10)
NC_final <- simulator(geno_freqs_v19_norm[5,2:19], 10)
NE_final <- simulator(geno_freqs_v19_norm[6,2:19], 10)
PA_final <- simulator(geno_freqs_v19_norm[7,2:19], 10)
TN_final <- simulator(geno_freqs_v19_norm[8,2:19], 10)
TX_final <- simulator(geno_freqs_v19_norm[9,2:19], 10)

vFinal.df <- obs.exp.df(male_freqs, female_freqs, CA_final, FL_final, GA_final, KS_final, NC_final, NE_final, PA_final, TN_final, TX_final)


# Plot simulation results for final frequencies
gen.plot.obs(CA_final, male_freqs$YMfreq[1], female_freqs$traDfreq[1], "CA_simulation.pdf")
gen.plot.obs(FL_final, male_freqs$YMfreq[2], female_freqs$traDfreq[2], "FL_simulation.pdf")
gen.plot.obs(GA_final, male_freqs$YMfreq[3], female_freqs$traDfreq[3], "GA_simulation.pdf")
gen.plot.obs(KS_final, male_freqs$YMfreq[4], female_freqs$traDfreq[4], "KS_simulation.pdf")
gen.plot.obs(NC_final, male_freqs$YMfreq[5], female_freqs$traDfreq[5], "NC_simulation.pdf")
gen.plot.obs(NE_final, male_freqs$YMfreq[6], female_freqs$traDfreq[6], "NE_simulation.pdf")
gen.plot.obs(PA_final, male_freqs$YMfreq[7], female_freqs$traDfreq[7], "PA_simulation.pdf")
gen.plot.obs(TN_final, male_freqs$YMfreq[8], female_freqs$traDfreq[8], "TN_simulation.pdf")
gen.plot.obs(TX_final, male_freqs$YMfreq[9], female_freqs$traDfreq[9], "TX_simulation.pdf")


# Create object with final simulated frequencies for each population
# and observed frequencies
final_freqs <- cbind(
	subset(male_freqs, select=c(Pop, YMfreq)), 
	subset(female_freqs, select=c(traDfreq, Mfreq, Myfreq)),
	rbind(CA_final[11,], FL_final[11,], GA_final[11,], KS_final[11,], NC_final[11,], NE_final[11,], PA_final[11,], TN_final[11,], TX_final[11,]), 
	Y_male = NA, IIIM_male=NA, traD_female = NA
)
final_freqs[, 43:45] <- t(apply(final_freqs[,7:24], 1, allele.freqs.sex))

# calculate difference between obs and simulated frequencies
final_freqs$Ym_OE <- final_freqs$YMfreq - final_freqs$Y_male
final_freqs$traD_OE <- final_freqs$traDfreq - final_freqs$traD_female

# % deviation between obs and simulated
final_freqs$Ym_percDev <- 100*final_freqs$Ym_OE/final_freqs$YMfreq 
final_freqs$traD_precDev <- 100*final_freqs$traD_OE/final_freqs$traDfreq


# Plot frequency of YM, IIIM, Multi-M males traD in each population
states <- c("CA", "TX", "KS", "NE", "FL", "GA", "TN", "NC", "PA")

final_freqs$state <- factor(final_freqs$Pop, levels=states)

# % muti-M males
#10 male   m1 M1 F0 Y0	IIIM/III	tra+/tra+	X/X
#11 male   m2 M2 F0 Y0	IIIM/IIIM	tra+/tra+	X/X
#12 male   m3 M0 F0 Y1	III /III	tra+/tra+	Y/X
#13 male   m4 M1 F0 Y1	IIIM/III	tra+/tra+	Y/X
#14 male   m5 M2 F0 Y1	IIIM/IIIM	tra+/tra+	Y/X
#15 male   m6 M0 F0 Y2	III /III	tra+/tra+	Y/Y
#16 male   m7 M1 F0 Y2	IIIM/III	tra+/tra+	Y/Y
#17 male   m8 M2 F0 Y2	IIIM/IIIM	tra+/tra+	Y/Y
multi.M <- function(genotypes){
	100*sum(genotypes[c(11,13,14,15,16,17)]) / (sum(genotypes[10:17]))
}
final_freqs$multi_M <- apply(final_freqs[,7:24], 1, multi.M)

# Export Final Frequencies
write.table(final_freqs, file="final_freqs.tsv", quote=FALSE, sep="\t", row.names=FALSE)


# Make plot 
library(cowplot)
library(ggplot2)

# code states by west to east order using factor
states <- c("CA", "TX", "KS", "NE", "FL", "GA", "TN", "NC", "PA")
final_freqs$states <-  factor(final_freqs$state, levels=states) # order states as factors sorted by west to east


YM_freq <- ggplot(final_freqs, aes(x=states, y=Y)) + 
	geom_bar(stat="identity", color="blue", fill="blue") + 
	scale_y_continuous(expression(freq~Y^M)) + 
	theme_bw() +
	theme(axis.title.x=element_blank()) 

IIIM_freq <- ggplot(final_freqs, aes(x=states, y=IIIM)) + 
	geom_bar(stat="identity", color="orange", fill="orange") + 
	scale_y_continuous(expression(freq~III^M)) + 
	theme_bw()  +
	theme(axis.title.x=element_blank()) 


traD_freq <- ggplot(final_freqs, aes(x=states, y=traD)) + 
	geom_bar(stat="identity", color="magenta", fill="magenta") + 
	scale_y_continuous(expression(freq~Md-tra^D)) + 
	theme_bw()  +
	theme(axis.title.x=element_blank()) 


pdf("final_freqs.pdf", width=3, height=4)
plot_grid(YM_freq, IIIM_freq, traD_freq, ncol = 1, align = "v") 
dev.off()


# scatter plots of traD freq vs multi-M
pdf("traD_multiM.pdf", width=4, height=4)
	plot(x=final_freqs$multi_M, y=100*final_freqs$traD_female, type='n', xlab="% Multi-M males", ylab=expression('%'~tra^D~females))
	text(x=final_freqs$multi_M, y=100*final_freqs$traD_female, label=final_freqs$Pop)
dev.off()


# Calculate correlation of tra^D frequency and multi-M frequency
traD.multiM <- lm(100*traD_female ~ multi_M, data=final_freqs)
traD.multiM
#Coefficients:
#(Intercept)      multi_M  
#      1.982        2.249  
summary(traD.multiM)
#Residuals:
#    Min      1Q  Median      3Q     Max 
#-2.0281 -1.8374 -0.3673  1.5891  2.8431 
#Coefficients:
#            Estimate Std. Error t value Pr(>|t|)    
#(Intercept)   1.9823     0.9917   1.999   0.0858 .  
#multi_M       2.2490     0.1267  17.745 4.45e-07 ***
#---
#Signif. codes:  0 ‘***’ 0.001 ‘**’ 0.01 ‘*’ 0.05 ‘.’ 0.1 ‘ ’ 1
#Residual standard error: 2.157 on 7 degrees of freedom
#Multiple R-squared:  0.9783,	Adjusted R-squared:  0.9751 
#F-statistic: 314.9 on 1 and 7 DF,  p-value: 4.451e-07


########
# Compare with published frequencies
#								Chino1982	Chino2014	CA 2021	NC 2002		NC 2006	NC 2007		NC 2021		
# m3	X/Y; III/III	males	68.00%		76.00%		NA		77.65%		77.80%	95.30%		NA			
# m1	X/X; IIIM/III	males	0.00%		8.00%		NA		20.00%		19.40%	2.30%		NA			
# m4	X/Y; IIIM/III	males	12.00%		4.00%		NA		2.35%		1.40%	0.00%		NA			
# m6	Y/Y; III/III	males	-			12.00%		NA		0.00%		1.40%	2.30%		NA			
#		Y^M				males	92.00%		27.38%		NA		80.60%		97.60%	70.41%		NA
#		III^M			males	12.00%		83.42%		NA		20.80%		2.30%	29.61%		NA
#		Md-traD			females	-			33.60%		NA		5.90%		4.20%			
#								Meisel et al. 2016			Hamm et al. 2005	Hamm and Scott 2008			

freq.times <- function(finals){
	c(	100*finals[18] / sum(finals[16:23]), 
		100*finals[16] / sum(finals[16:23]), 
		100*finals[19] / sum(finals[16:23]), 
		100*finals[21] / sum(finals[16:23]), 
		100*finals$Y_male, 
		100*finals$IIIM_male, 
		100*finals$traD_female
	)
}

freq_time.df <- data.frame(
	Genotype = c("X/YM; III/III", "X/X; IIIM/III", "X/YM; IIIM/III", "YM/YM; III/III", "Y^M", "III^M", "Md-traD"),
	Sex = c(rep("male", 6), "female"), 
	Chino_1982 = c(68.00, 0.00, 12.00, rep(NA, 4)), 
	Chino_2014 = c(76.00, 8.00, 4.00, 12.00, 92.00, 12.00, 33.60),
	CA_2021 = unlist(freq.times(final_freqs[1,])), 
	NC_2002 = c(77.65, 20.00, 2.35, 0.00, 80.00, 22.35, 5.90), 
	NC_2006 = c(77.80, 19.40, 1.40, 1.40, 80.60, 20.80, NA), 
	NC_2007 = c(95.30, 2.30, 0.00, 2.30, 97.60, 2.30, 4.20),
	NC_2021 = unlist(freq.times(final_freqs[5,]))
)


write.table(freq_time.df, file="freq_time.tsv", quote=FALSE, sep="\t", row.names=FALSE)


##########
# Analyze climate data
# final_freqs <- read.delim(file="final_freqs.tsv")

# calculate frequency of males that have both YM and IIIM
# 1 female f1 M0 F0 Y0	III /III	tra+/tra+	X/X
# 2 female f2 M0 F1 Y0	III /III	traD/tra+	X/X
# 3 female f3 M1 F1 Y0	IIIM/III	traD/tra+	X/X
# 4 female f4 M2 F1 Y0	IIIM/IIIM	traD/tra+	X/X
# 5 female f5 M0 F1 Y1	III /III	traD/tra+	X/Y
# 6 female f6 M1 F1 Y1	IIIM/III	traD/tra+	X/Y
# 7 female f7 M2 F1 Y1	IIIM/IIIM	traD/tra+	X/Y
# 8 female f8 M0 F1 Y2	III /III	traD/tra+	Y/Y
# 9 female f9 M1 F1 Y2	IIIM/III	traD/tra+	Y/Y
#10 male   m1 M1 F0 Y0	IIIM/III	tra+/tra+	X/X
#11 male   m2 M2 F0 Y0	IIIM/IIIM	tra+/tra+	X/X
#12 male   m3 M0 F0 Y1	III /III	tra+/tra+	Y/X
#13 male   m4 M1 F0 Y1	IIIM/III	tra+/tra+	Y/X
#14 male   m5 M2 F0 Y1	IIIM/IIIM	tra+/tra+	Y/X
#15 male   m6 M0 F0 Y2	III /III	tra+/tra+	Y/Y
#16 male   m7 M1 F0 Y2	IIIM/III	tra+/tra+	Y/Y
#17 male   m8 M2 F0 Y2	IIIM/IIIM	tra+/tra+	Y/Y
#18 female f10M2 F1 Y2	IIIM/IIIM	traD/tra+	Y/Y
Y3.freq <- function(datas){
	sum(datas[c(13,14,16,17)]) / sum(datas[10:17])
}
final_freqs$Y3_freq <- apply(final_freqs[,7:24], 1, Y3.freq)


# Coordinates
#				CA			FL			GA			KS			NC			NE			PA			TN			TX     
#	lat = c(  33.858630,  29.7946429,  34.273960,  39.186691,  35.722480,  40.813618,  40.7944504,  35.772283,  30.67), 
#	lon = c(-117.020450, -82.4102420, -83.594450, -96.566597, -78.680820, -96.702599, -77.8616386, -83.851272, -96.35)


CA_monthly <- read.csv("CA-normals-monthly-1991-2020-2023-07-27T17-04-38.csv")
FL_monthly <- read.csv("FL-normals-monthly-1991-2020-2023-07-27T17-09-41.csv")
GA_monthly <- read.csv("GA-normals-monthly-1991-2020-2023-07-27T17-24-59.csv")
KS_monthly <- read.csv("KS-normals-monthly-1991-2020-2023-07-27T17-24-28.csv")
NC_monthly <- read.csv("NC-normals-monthly-1991-2020-2023-07-27T17-28-26.csv")
NE_monthly <- read.csv("NE-normals-monthly-1991-2020-2023-07-27T17-29-51.csv")
PA_monthly <- read.csv("PA-normals-monthly-1991-2020-2023-07-27T17-42-49.csv")
TN_monthly <- read.csv("TN-normals-monthly-1991-2020-2023-07-31T12-25-45.csv")
TX_monthly <- read.csv("TX-normals-monthly-1991-2020-2023-07-31T12-31-44.csv")

CA_annual <- read.csv("CA-normals-annualseasonal-1991-2020-2023-07-27T17-11-15.csv")
FL_annual <- read.csv("FL-normals-annualseasonal-1991-2020-2023-07-27T17-12-08.csv")
GA_annual <- read.csv("GA-normals-annualseasonal-1991-2020-2023-07-27T17-25-22.csv")
KS_annual <- read.csv("KS-normals-annualseasonal-1991-2020-2023-07-27T17-25-54.csv")
NC_annual <- read.csv("NC-normals-annualseasonal-1991-2020-2023-07-27T17-28-03.csv")
NE_annual <- read.csv("NE-normals-annualseasonal-1991-2020-2023-07-27T17-30-12.csv")
PA_annual <- read.csv("PA-normals-annualseasonal-1991-2020-2023-07-27T17-42-21.csv")
TN_annual <- read.csv("TN-normals-annualseasonal-1991-2020-2023-07-31T12-26-15.csv")
TX_annual <- read.csv("TX-normals-annualseasonal-1991-2020-2023-07-31T12-30-01.csv")

Annuals <- cbind(
	Pop = final_freqs$Pop, 
	rbind(CA_annual, FL_annual, GA_annual, KS_annual, NC_annual, NE_annual, PA_annual, TN_annual, TX_annual)
)

# Analyzing: 
# ANN.TAVG.NORMAL						Tmean 		Annual mean average daily temperature
# ANN.TMIN.NORMAL						Tmin 		Annual mean minimum daily temperature
# ANN.TMAX.NORMAL						Tmax 		Annual mean maximum daily temperature
# ANN.TMAX.NORMAL - ANN.TMIN.NORMAL		DailyTR 	Annual mean daily temperature range
# ANN.PRCP.NORMAL						Precip		Annual Precipitation
# max(MLY.TAVG.NORMAL)					Tactive 	Mean temperature of warmest month
# cv(MLY.TAVG.NORMAL)					Season1 	Coefficient of variation of average monthly temperatures
# 0.5*((max(MLY.TMAX.NORMAL) - min(MLY.TMAX.NORMAL)) + (max(MLY.TMIN.NORMAL) - min(MLY.TMIN.NORMAL)))
#										Season2 	Average of difference between highest and lowest monthly maximum temperature
#						 						and difference between highest and lowest monthly minimum temperature
# mean(monthly$MLY.TAVG.NORMAL[5:7])	SummerMean	Mean average monthly temperature during May-July 
# mean(monthly$MLY.TMAX.NORMAL[5:7])	SummerMax	Mean maximum monthly temperature during May-July 
# mean(monthly$MLY.TMIN.NORMAL[5:7])	SummerMin	Mean minimum monthly temperature during May-July 

# Not analyzing: 
# Alt Altitude
# Lat Latitude relative to mean latitude per study
# Humidity Annual mean relative humidity
# Frost Annual number of days with ground frost
# Sunshine Annual percentage of sunshine hours

# convert to celcius
# C = (°F − 32) × 5/9
F.to.C <- function(Farn){
	(Farn - 32) * 5/9
}


# Summer uses May, June, July weather only

monthlies <- function(monthly){
	c(
	 	Tactive = max(F.to.C(monthly$MLY.TAVG.NORMAL)), # Tactive
		Season1 = sd(F.to.C(monthly$MLY.TAVG.NORMAL)) / mean(F.to.C(monthly$MLY.TAVG.NORMAL)), # Season1
		Season2 = 0.5*((max(F.to.C(monthly$MLY.TMAX.NORMAL)) - min(F.to.C(monthly$MLY.TMAX.NORMAL))) + (max(F.to.C(monthly$MLY.TMIN.NORMAL)) - min(F.to.C(monthly$MLY.TMIN.NORMAL)))), #Season2
		SummerMean = mean(F.to.C(monthly$MLY.TAVG.NORMAL[5:7])),
		SummerMax = mean(F.to.C(monthly$MLY.TMAX.NORMAL[5:7])),
		SummerMin = mean(F.to.C(monthly$MLY.TMIN.NORMAL[5:7]))
	)
}


final_freqs$Tmean <- F.to.C(Annuals$ANN.TAVG.NORMAL)
final_freqs$Tmin <- F.to.C(Annuals$ANN.TMIN.NORMAL)
final_freqs$Tmax <- F.to.C(Annuals$ANN.TMAX.NORMAL)
final_freqs$DailyTR <- F.to.C(Annuals$ANN.TMAX.NORMAL) - F.to.C(Annuals$ANN.TMIN.NORMAL)
final_freqs$Precip <- 2.54*Annuals$ANN.PRCP.NORMAL # convert to cm
final_freqs <- cbind(final_freqs, 
	rbind(
		monthlies(CA_monthly),
		monthlies(FL_monthly),
		monthlies(GA_monthly),
		monthlies(KS_monthly),
		monthlies(NC_monthly),
		monthlies(NE_monthly),
		monthlies(PA_monthly),
		monthlies(TN_monthly),
		monthlies(TX_monthly)
	)
)


# Calculate correlations between climate factors and allele/genotype frequencies
correlations <- function(genetic, environ){
	c(
		cor.test(genetic, environ, method="spearman")$estimate, 
		p = cor.test(genetic, environ, method="spearman")$p.value 	
	)
}

gen.freqs <- subset(final_freqs, select=c(Y, IIIM, traD, multi_M, Y3_freq))
climates  <- subset(final_freqs, select=c(Tmean, Tmin, Tmax, DailyTR, Precip, Tactive, Season1, Season2, SummerMean, SummerMax, SummerMin))


cor(gen.freqs, climates)


correlations.df <- data.frame( # create dummy object
	genetic = rep(NA, length(gen.freqs)*length(climates)),
	climate = rep(NA, length(gen.freqs)*length(climates)),
	rho = rep(NA, length(gen.freqs)*length(climates)),
	p = rep(NA, length(gen.freqs)*length(climates))
) 
counter <- 1
for(i in 1:length(gen.freqs)){
	for(j in 1:length(climates)){
		correlations.df[counter, 1] <- colnames(gen.freqs)[i]
		correlations.df[counter, 2] <- colnames(climates)[j]
		correlations.df[counter, 3:4] <- correlations(gen.freqs[,i], climates[,j])
		counter <- counter + 1
	}
}


# Export all correlations
write.table(correlations.df, "eco_correlations.tsv", quote=FALSE, sep="\t", row.names=FALSE)


correlations.df[order(correlations.df$p), ]
subset(correlations.df, p<0.05)
# DailyTR is the only thing with a significant correlation with a genotype (YM and IIIM frequencies)


# Principal component analysis of annual climate data
row.names(climates) <- c("CA","FL","GA","KS","NC","NE","PA","TN","TX")
climate_pca <- prcomp(climates[,1:8], scale.=TRUE)


# Plot PCA and DailyTR correlation
library(ggfortify)
climate_pca.p <- autoplot(climate_pca, data=climates[,c(1:8)], shape = FALSE, loadings = TRUE, loadings.label = TRUE) + 
					theme_bw() + 
				theme(axis.text=element_text(size=12), axis.title=element_text(size=12)) 

plot_freqs <- cbind(
	Pop = final_freqs$Pop, 
	gen.freqs, 
	climates
)

DailyTR_YM <- ggplot(plot_freqs, aes(x=DailyTR, y=Y)) + 
				geom_text(aes(label=Pop)) + 
				scale_y_continuous(expression(Y^M~"frequency")) + 
				scale_x_continuous("") + 
				theme_bw() + 
				theme(axis.text=element_text(size=12), axis.title=element_text(size=12)) 

DailyTR_IIIM <- ggplot(plot_freqs, aes(x=DailyTR, y=IIIM)) + 
				geom_text(aes(label=Pop)) + 
				scale_y_continuous(expression(III^M~"frequency")) + 
				scale_x_continuous("") + 
				theme_bw() + 
				theme(axis.text=element_text(size=12), axis.title=element_text(size=12)) 


pdf("PCA_DailyTR.pdf", width=12, height=4)
ggdraw(plot_grid(climate_pca.p, DailyTR_YM, DailyTR_IIIM, labels="AUTO", ncol=3, rel_widths=c(1,1,1)) + 
			draw_label("annual mean daily temperature range (Celcius)", x=.66, y=.05)) # manually draw x-axis label
dev.off()


# First 3 PCAs explain >98% of variance 
summary(climate_pca)
# Importance of components:
#                           PC1    PC2     PC3     PC4     PC5      PC6       PC7
# Standard deviation     2.2903 1.3958 0.81990 0.33752 0.14185 0.01560 0.002342 6.014e-17
# Proportion of Variance 0.6557 0.2435 0.08403 0.01424 0.00252 0.00003 0.000000 0.000e+00
# Cumulative Proportion  0.6557 0.8992 0.98321 0.99745 0.99997 1.00000 1.000000 1.000e+00


# Test for correlation between first three PCs and chromosome frequencies
corr_pca <- data.frame( # create dummy object
	genetic = rep(NA, length(gen.freqs)*3),
	climate = rep(NA, length(gen.freqs)*3),
	rho = rep(NA, length(gen.freqs)*3),
	p = rep(NA, length(gen.freqs)*3)
) 
counter <- 1
for(i in 1:length(gen.freqs)){
	for(j in 1:3){ # only cycle through the first 3 PCAs
		corr_pca[counter, 1] <- colnames(gen.freqs)[i]
		corr_pca[counter, 2] <- c("PC1", "PC2", "PC3")[j]
		corr_pca[counter, 3:4] <- correlations(gen.freqs[,i], climate_pca$x[,j])
		counter <- counter + 1
	}
}


corr_pca
# PC2 is significantly correlated with Y^M frequency (p=0.0214)


# Test for interaction between PC1 and PC2 in linear model 
freqs_pca <- cbind(
	Pop = final_freqs$Pop, 
	gen.freqs, 
	climate_pca$x
)
anova(lm(Y ~ PC1 * PC2, data = freqs_pca))
anova(lm(IIIM ~ PC1 * PC2, data = freqs_pca))


###########
# exclude CA population

# Calculate correlations between climate factors and allele/genotype frequencies

gen.freqs_noCA <- gen.freqs[2:9,]
climates_noCA  <- climates[2:9,]

correlations_noCA.df <- data.frame( # create dummy object
	genetic = rep(NA, length(gen.freqs_noCA)*length(climates_noCA)),
	climate = rep(NA, length(gen.freqs_noCA)*length(climates_noCA)),
	rho = rep(NA, length(gen.freqs_noCA)*length(climates_noCA)),
	p = rep(NA, length(gen.freqs_noCA)*length(climates_noCA))
) 
counter <- 1
for(i in 1:length(gen.freqs_noCA)){
	for(j in 1:length(climates_noCA)){
		correlations_noCA.df[counter, 1] <- colnames(gen.freqs_noCA)[i]
		correlations_noCA.df[counter, 2] <- colnames(climates_noCA)[j]
		correlations_noCA.df[counter, 3:4] <- correlations(gen.freqs_noCA[,i], climates_noCA[,j])
		counter <- counter + 1
	}
}

correlations_noCA.df[order(correlations_noCA.df$p), ]
subset(correlations_noCA.df, p<0.05)
# DailyTR is still significantly correlated with YM frequency! 


# Principal component analysis of annual climate data
row.names(climates_noCA) <- c("FL","GA","KS","NC","NE","PA","TN","TX")
climate_pca_noCA <- prcomp(climates_noCA[,1:8], scale.=TRUE)


# Plot PCA and DailyTR correlation
pdf("climate_pca_noCA.pdf", width=5, height=5)
autoplot(climate_pca_noCA, data=climates_noCA[,c(1:8)], shape = FALSE, loadings = TRUE, loadings.label = TRUE) + 
	theme_bw() + 
	theme(axis.text=element_text(size=12), axis.title=element_text(size=12)) 
dev.off()


summary(climate_pca_noCA)
# Importance of components:
#                           PC1    PC2     PC3     PC4     PC5      PC6      PC7       PC8
# Standard deviation     2.3948 1.3306 0.58142 0.39063 0.05903 0.007138 0.002251 2.107e-16
# Proportion of Variance 0.7169 0.2213 0.04226 0.01907 0.00044 0.000010 0.000000 0.000e+00
# Cumulative Proportion  0.7169 0.9382 0.98048 0.99956 0.99999 1.000000 1.000000 1.000e+00
```
